# Pyranthiones/Pyrones: “Click and Release” Donors for Subcellular Hydrogen Sulfide Delivery and Labeling

**DOI:** 10.1101/2023.06.19.545622

**Authors:** Wei Huang, Jorge Escorihuela, Scott T. Laughlin

## Abstract

Hydrogen sulfide (H_2_S), one of the most important gasotransmitters, plays a critical role in endogenous signaling pathways of many diseases. However, developing H_2_S donors with both tunable release kinetics and high release efficiency for subcellular delivery has been challenging. Here, we describe a click and release reaction between pyrone/pyranthiones and bicyclononyne (BCN). This reaction features a release of CO_2_/COS with second-order rate constants comparable to Strain-Promoted Azide-Alkyne Cycloaddition reactions (SPAACs). Interestingly, pyranthiones showed enhanced reaction rates compared to their pyrone counterparts. We investigated pyrone biorthogonality and demonstrated their utility in protein labeling applications. Moreover, we synthesized substituted pyranthiones with H_2_S release kinetics that can address the range of physiologically relevant H_2_S dynamics in cells and achieved quantitative H_2_S release efficiency *in vitro*. Finally, we explored the potential of pyranthiones as H_2_S/COS donors for mitochondrial-targeted H_2_S delivery in living cells.

## Main

Bioorthogonal click and release reactions are powerful tools for the delivery of cargo with good stability and controllable release^[1]^. For example, extensive work has been done on the activation of anticancer prodrugs using the Staudinger-Bertozzi ligation^[2]^, strain-promoted 1,3-dipolar cycloadditions between azides and *trans*-cyclooctene^[3]^, and inverse-electron-demand Diels-Alder cycloadditions (IEDDA) between tetrazines and *trans*-cyclooctenes^[4]^. Recently, there has been growing interest in the development of bioorthogonal activation of prodrugs for the delivery of gasotransmitters, including carbon monoxide, hydrogen sulfide, and nitric oxide, which are now recognized to play vital roles in endogenous signaling pathways^[5]^.

Dysregulation of hydrogen sulfide (H_2_S), one of the endogenous gasotransmitters for oxidative/reductive stress regulation, is linked to cardiovascular, gastrointestinal, neurological, and endocrine diseases^[6–8]^.

One likely mechanism for H_2_S-linked disease development is its role in regulating energy production in mitochondria^[9]^. It has been reported that the sulfide level in human plasma is ∼3 μM with free hydrogen sulfide levels of 0.2–0.8 μM^[10]^. Unlike conventional H_2_S donors that employ inorganic sulfide salts and an instantaneous H_2_S concentration surge, or organic small molecule donors like AP39, which generate H_2_S with un-tunable release rates triggered by bio-thiols, Pluth and coworkers reported a click and release strategy for controllable H_2_S release^[11]^. They modified *trans*-cyclooctene with a benzylic thiocarbamate as a masked H_2_S donor, which undergoes self-immolation to produce carbonyl sulfide (COS) after reaction with a tetrazine. The generated COS can then be converted into H_2_S by carbonic anhydrase, a ubiquitous enzyme inside cells^[12]^. Though promising *in vitro*, this COS/H_2_S donor’s slow-release rate and susceptibility to tetrazine reduction by the released H_2_S stymied its straightforward application in living cells. Later, Taran and coworkers showed that 1,3-dithiolium-4-olates (DTO) could be an alternative H_2_S/COS donor when reacting with strained alkynes^[13]^. However, the release of H_2_S was not demonstrated. More recently, inspired by Taran’s work, Liang and coworkers developed a mitochondrial targeting H_2_S/COS donor by introducing a diphenylamino substituent on DTO^[14]^. This approach had the added benefit of generating a green fluorophore after reaction with bicyclononynes (BCN). Albeit useful, the reaction between DTO and BCN provides modest H_2_S release efficiency (∼40%) and limited potential for tuning the H_2_S release kinetics, which are two pillars of H_2_S donor development.

2-Pyrones have been widely used as building blocks for enantioselective total synthesis as well as bioactive compounds like antibiotics^[15,16]^. Proceeding via Diels-Alder (DA) reaction with alkynes or alkenes followed by a retro DA reaction by releasing carbon dioxide (CO_2_), aromatic compounds can be synthesized efficiently from 2-pyrones^[17]^. Further, 2-pyrones are important cellular metabolites that can be generated by several biosynthetic pathways^[15]^. Nevertheless, this useful synthon has received little attention as a potential bioorthogonal handle due to the required harsh reaction conditions like high temperature or pressure^[18,19]^. We envisioned that the requirement for harsh reaction conditions could be mitigated though the addition of electron withdrawing groups that accelerate the reaction of 2-pyrones with alkenes and alkynes. Moreover, the related compounds, pyranthiones, could be potential H_2_S/COS donors when reacting with electron-rich and/or strained alkynes/alkenes.

Our initial interest in pyrones/pyranthiones stemmed from their potential use as reaction partners with newly developed, modular activatable cyclopropenes^[20,21]^ and cyclopropene-modified neurotransmitters for bioorthogonal detection^[22]^. Thus, to begin exploring functionalized pyrones and pyranthiones as bioorthogonal reagents, we combined an ester functionalized pyrone and a cyclopropene commonly used in bioorthogonal studies ^[23]^, but no reaction was observed (**Figure 1A**). In contrast, the reaction proceeded smoothly when a strained alkyne, BCN, was used (**Figure 1B**). We reasoned that spontaneous release of CO_2_ driven by aromatization might accelerate the reaction. To further explore enhanced reaction rates and the potential for bioorthogonal gasotransmitter release, we designed and synthesized four pyrone/pyranthione analogues bearing different electron withdrawing groups (**Figure 2**). These compounds enabled us to examine the relationship between reactivity and biorthogonality and explore pyranthiones as potential H_2_S/COS donors with both tunable release kinetics as well as high release efficiency in subcellular H_2_S delivery.

**Figure 1.**
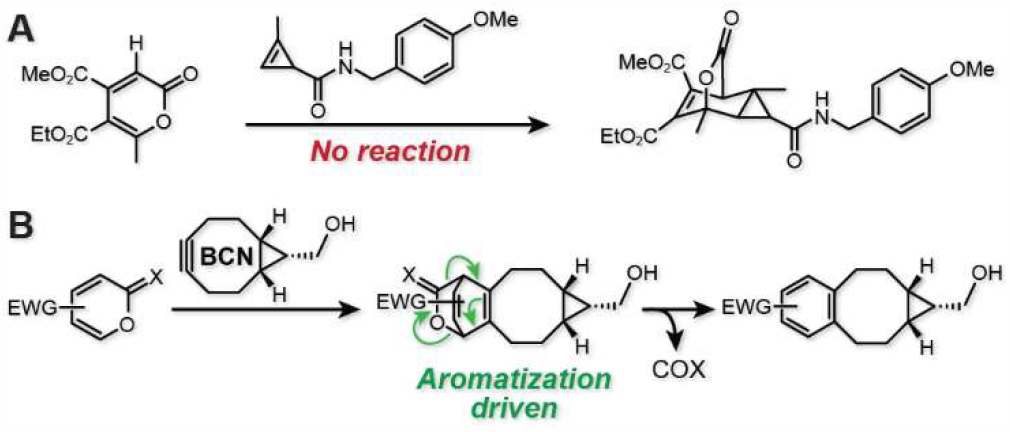
Inverse electron demand Diels-Alder cycloadditions between strained alkene/alkyne and electron deficient 2-pyrone. **A**, A common cyclopropene used in bioorthogonal reactions showed no reaction towards 2-pyrone. **B**, *endo*-BCN underwent cycloaddition with 2-pyrone/pyranthione followed by retro-Diels-Alder with concomitant release of CO_2_ or COS driven by aromatization.

**Figure 2.**
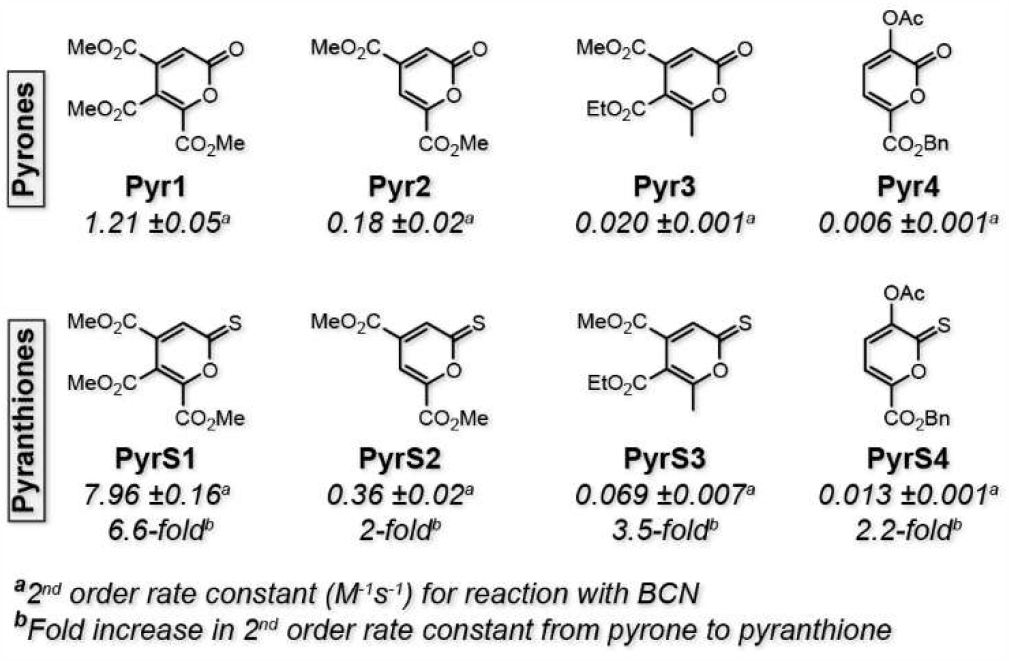
Kinetics studies for the IEDDA cycloaddition between 2-pyrones/pyranthiones and BCN. Second-order rate constants were derived from pseudo first-order rates using 10 to 20 : 1 ratio of BCN to 2-pyrone/pyranthione in 50% H_2_O/CH_3_CN. Rates are shown as mean values from linear fitting between pseudo first-order rates versus concentration of BCN ± standard deviation.

To demonstrate how the electron withdrawing groups influence the reactivity of 2-pyrones, we synthesized four pyrone derivatives, Pyr1, Pyr2, Pyr3, and Pyr4, with different numbers of ester groups appending to various positions (**Figure 2**). The corresponding pyranthione counterparts were also readily synthesized by refluxing pyrone with Lawesson’s reagent in toluene with decent yields (30% to 71%) (**Supporting Information** Synthetic details). Next, we determined the 2^nd^ order rate constants between *endo-*BCN-OH (BCN) and 2-pyrones by following the decay of the peaks of 2-pyrones using HPLC or ^1^H NMR (**Figure 2** and **S1**). Not surprisingly, Pyr1, with the highest number of ester groups, showed the highest reaction rates, achieving up to 1.2 M^-1^·s^-1^, comparable to SPAACs. In contrast, more sluggish kinetics were observed for pyrones with a smaller number of electron withdrawing groups, showing a 2^nd^ order rate constant of 0.18, 0.02, and 0.006 M^-1^S^-1^ for Pyr2, Pyr3, and Pyr4, respectively. In addition, a 9-fold difference of reaction rate between Pyr2 and Pyr3 implied the importance of the substitution position as well as steric effects on the reactivity of 2-pyrones. The kinetics between pyranthione the counterparts and BCN were also determined by following the decay of the characteristic absorption at 412 nm (**Figure S2, S3**, and **S4**). Similar reactivity trends were observed for pyranthione derivatives, showing 2^nd^ order rate constants of 7.96, 0.36, 0.069, and 0.013 M^-1^S^-1^ for PyrS1, PyrS2, PyrS3, PyrS4, respectively. All pyranthione derivatives showed enhanced kinetics (2- to 6-fold) compared with their 2-pyrone counterparts, which is counterintuitive since pyranthiones are considered more aromatic than pyrones^[18]^.

To begin exploring the rationale for the reactivity difference between pyrone and pyranthione we performed a computational study using DFT calculations with Gaussian 16 (**Supporting Information** Computational details). The computed transition state structures at the M06-2X/6-311+G(d,p) level of theory and the activation energy (Δ*E*^‡^), activation free energy (Δ*G*^‡^), and free energy of the reaction (Δ*G*_*rxn*_) are shown in **Figure 3A**. The activation free energies for 2-pyrones (Pyr1, Pyr2, Pyr3, and Pyr4) were higher than that of the homologous pyranthione derivatives (PyrS1, PyrS2, PyrS3 and PyrS4) and followed the same trend observed experimentally. The cycloaddition reaction proceeds in all cases through non-synchronous transition states with activation free energy (Δ*G*^‡^) in the range of 15.1–21.0 kcal/mol and highly exergonic processes (ΔG_R_ < 0). Pyr1 and PyrS1, with the highest number of ester groups, show earlier transition states and a longer forming C−C bond distance than other pyrone or pyranthione derivatives, respectively, and accordingly lower activation free energies.

**Figure 3.**
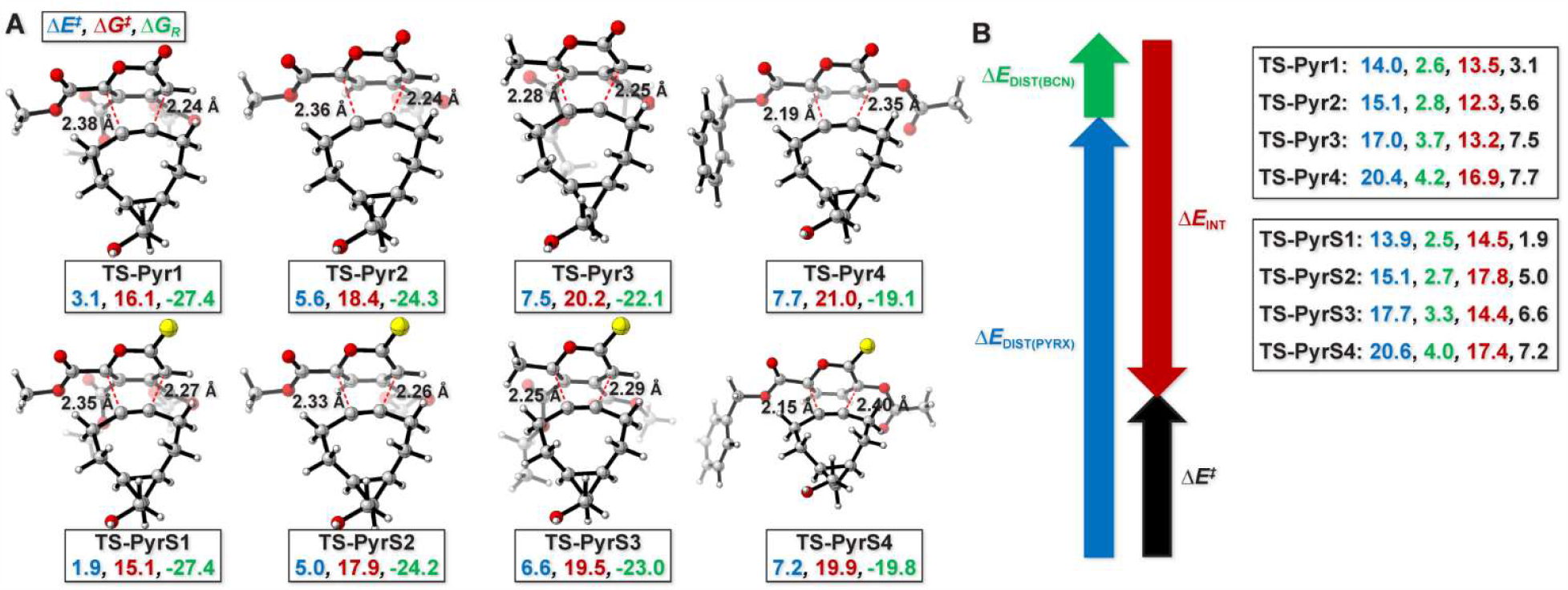
**A**, Optimized transition state structures at the M06-2X/6-311+G(d,p) level of theory with bond forming distances in Å. Values of activation energy (Δ*E*^‡^), activation free energy (Δ*G*^‡^), and free energy of the reaction (Δ*G*_*rxn*_) are shown below each structure in blue, red, and green, respectively, in kcal mol^−1^. **B**, Analysis of distortion, interaction, and activation (blue: distortion energy of PyrX (X = 1–4, S1–S4), green: distortion energy of BCN, red: interaction energy, black: activation energy). Energies are given in kcal mol^−1^.

In order to quantitatively analyze the factors behind the reactivity, we used the Activation Strain Model (ASM)^[24]^, which assess the relative contributions of the distortion (Δ*E*_dist_) and interaction (Δ*E*_int_) terms upon the approach of BCN to the pyrone or pyranthione. The distortion energy can be decomposed into the distortion energy of both reactants, *i*.*e*., the distortion energy of PyrX (X = 1–4 or S1–S4), and the distortion energy of BCN (**Figure 3B**). The change in BCN distortion is small across the series of pyrone or pyranthione derivatives (1.5–1.6 kcal/mol, green values). PyrX distortion varies more (blue values), between 6.4 and 6.7 kcal/mol, for pyrones or pyranthiones, respectively. However, the change in interaction energies (red values) is most significant, and pyranthiones have higher interaction activation energies compared to the homologous pyrones, which is reflected in lower activation free energies.

Next, we began to test the biorthogonality and stability of pyrones and pyranthiones since successful biolabeling using clickable handles always requires biorthogonality towards bio-nucleophiles to diminish nonspecific reactions. To this end, we chose two of the most reactive pyrones, Pyr1 and Pyr2, and tested the biorthogonality towards three naturally occurring amino acids, glutathione, lysine, and serine by HPLC assay (**Figure S5, S6**). We found that Pyr1 reacted rapidly with glutathione, and, more slowly, with both lysine and serine, likely resulting from the ring’s electron deficiency priming it for nucleophilic attack. In comparison, Pyr2 showed negligible reaction towards these amino acids, highlighting the trade-off between reactivity and biorthogonality in this system.

Next, we sought to move forward with biological studies leveraging the Pyr2 scaffold, which necessitated its functionalization. We started the synthesis from α-ketoglutaric acid to prepare a pyrone derivative bearing a hydroxy group, compound HO-Pyr2 (**Scheme S1**). However, activation of the hydroxyl group by treating with phosgene followed by reaction with amino linker did not yield fruitful results (**Scheme S2**). Instead, we attached a bromopropane linker by treating HO-Pyr2 with 1,3-dibromopropane and potassium carbonate at 80 oC, affording Br-Pyr2 in 52% yield. Substitution of the Br for Az produced azide derivative Az-Pyr2 in 60% yield and provided a convenient clickable handle for subsequent bioconjugation reactions (**Scheme S1**).

To demonstrate the utility of the pyrone-BCN reaction, we explored *in vitro* protein labeling with in-gel fluorescence detection (**Figure 4**). To accomplish this, we synthesized a fluorophore-pyrone conjugate, sulforhodamine B-PEG4-pyrone using copper catalyzed azide-alkyne click chemistry, indicating the compatibility of the pyrone-BCN reaction with 1,3-dipolar cycloadditions. Separately, we prepared BCN-modified lysozyme by incubation with BCN *N*-hydroxysuccinimide ester in PBS and confirmed the BCN modification by mass spectrometry (**Figure S7**). Then we performed protein labeling experiments by treating BCN-functionalized lysozyme with sulforhodamine B-PEG4-pyrone (50-400 μM) in PBS for 4h at room temperature and analyzed by in-gel fluorescence (**Supporting Information** Protein labeling experiments). Labeling results showed the reaction proceeded in a concentration dependent manner, with the highest fluorescence observed reacting with sulforhodamine B-PEG4-pyrone (400 μM). The success of *in vitro* protein labeling indicated that the 2-pyrones could be a promising bioorthogonal reactive handle for labeling *in vitro* and, potentially, *in vivo*.

**Figure 4.**
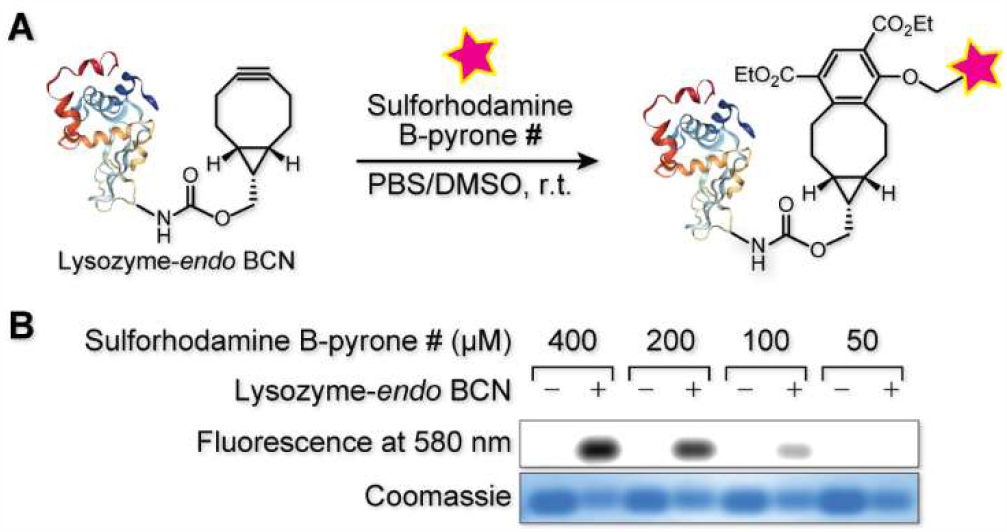
Demonstration of protein labeling using the ligation between BCN and pyrone-fluorophore conjugate. **A**, BCN modified lysozyme was labeled with sulforhodamine B-pyrone. **B**, In-gel fluorescence analysis of lysozyme-BCN or lysozyme (1.3 mg/mL) incubated with various concentrations of sulforhodamine B-pyrone (50–400 μM) in PBS at room temperature for 4h.

Inspired by the promising results of protein labeling using pyrone analogues, we next set up to examine the capabilities of pyranthione as H_2_S/COS donors *in vitro* (**Figure 5A**). We first used a fluorescence assay based on a turn-on fluorescent probe for H_2_S, NDI^[25]^, to evaluate bioorthogonal H_2_S release with physiological concentrations of H_2_S donor (**Supporting Information** Fluorescent assay of H_2_S release). PyrS2 was treated with BCN at submillimolar concentrations (100 μM) in the presence of Mito-NDI (10 μM) and carbonic anhydrase (50 μg/mL). The release of H_2_S was monitored by the increasing green fluorescence of NDI backbone freed by H_2_S substitution at 528 nm. As depicted in **Figure 5B**, PyrS2 showed steady H_2_S release while no significant changes of fluorescence were observed for 2-pyrone or amino acids control (**Figure S9**). Notably, the release rate of H_2_S seemed to be much slower without the presence of CA, suggesting that its important role in converting COS to H_2_S, which is consistent with previous reports^[12]^. H_2_S release with tunable rates were also performed with PyrS1, PyrS3 and PyrS4. However, both PyrS1 and PyrS4 did not show significant fluorescent signal after 12h incubation at room temperature (**Figure S10**). We reasoned that the vulnerability of PyrS1 to nucleophilic attack and the relatively slow kinetics of PyrS4 reaction with BCN might be responsible for the absence of fluorescence signals in this assay. Interestingly, the PyrS3 showed rapid turn on fluorescence, which was determined to be from the degradation in DMSO (**Figure S10** and **S11**).

**Figure 5.**
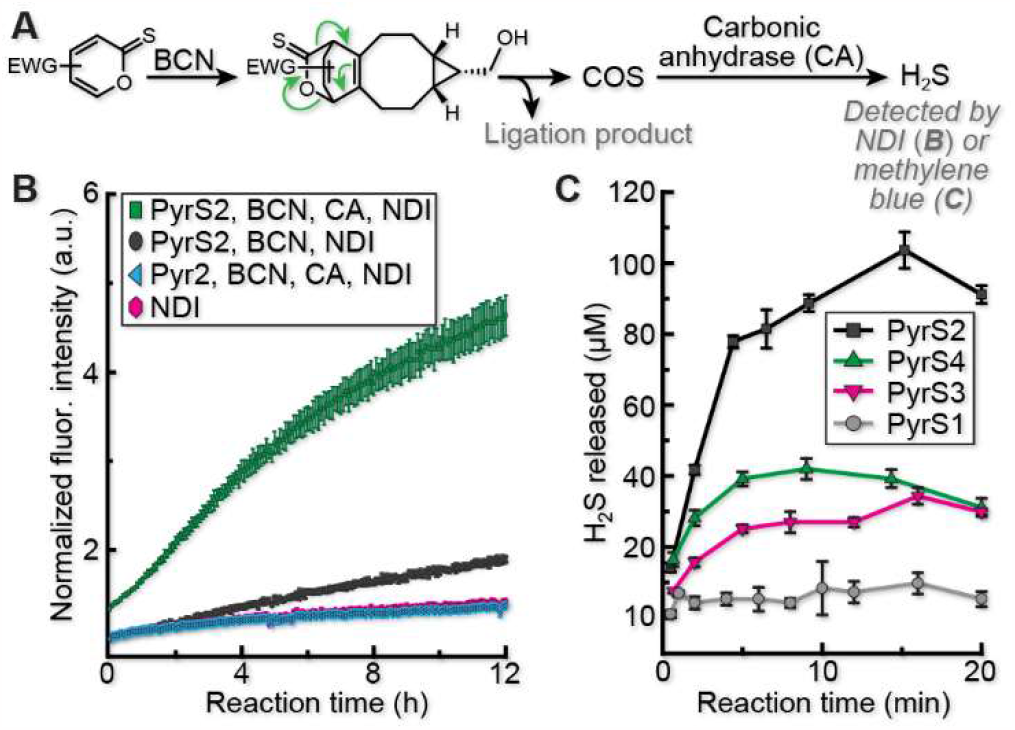
Demonstration of H_2_S release *in vitro*. **A**, Schematic demonstration of H_2_S release and detection after the ligation between pyranthiones and BCN. **B**, Turn-on fluorescent assay for the detection of generated H_2_S when incubation of PyrS2 (100 μM) with BCN (100 μM) using NDI (10 μM) as the H_2_S probe in the presence of carbonic anhydrase (50 μg/mL) in PBS at room temperature. **C**, Quantification of H_2_S released by using methylene blue assay after incubation of pyranthiones (100 μM) with BCN (1 mM) in the presence of carbonic anhydrase (50 μg/mL) in PBS at room temperature. Error bars in B and C represent standard deviation from three replicates.

Next, to further quantify the H_2_S releasing efficiency of pyranthiones, we used a methylene blue assay^[14]^ (**Supporting Information** Quantification of H_2_S release). As shown in **Figure 5C**, PyrS2 (100 μM), when treated with 1 mM of BCN, showed an almost quantitative H_2_S release efficiency in 15 minutes at room temperature, demonstrating a great potential for H_2_S/COS donors. In contrast, PyrS3 and PyrS5 exhibited decreased H_2_S releasing kinetics, which agreed well with our kinetics data for these compounds.

To explore the application of pyranthiones in subcellular targeted H_2_S delivery, we synthesized a pyranthione derivative, Br-PyrS2, by treating Br-Pyr2 with phosphorus pentasulfide at 80 oC in toluene with 41% yield (**Scheme S3**). Interestingly, this reaction did not show any conversion when Lawesson’s reagent was used, indicating the lower reactivity of Br-Pyr2 compared with Pyr2. To prepare a mitochondrial targeting pyranthione, PPh_3_-PyrS2, we tried refluxing Br-PyrS2 with triphenylphosphine in acetonitrile. However, we failed to isolate the product due to the domination of side reactions at high temperatures (above 80 oC) and low conversion with relative lower temperatures (60 oC to 70 oC). To facilitate the reaction’s success at lower temperatures, we synthesized an iodinated pyranthione, I-PyrS2, by treating Br-PyrS6 with sodium iodine at 40 °C. PPh_3_-PyrS2 was then prepared by reacting I-PyrS6 with PPh_3_ at 40 °C with a yield of 16% over two steps (**Scheme S4**). A mitochondrial targeting pyrone derivative, PPh_3_-Pyr2, was also synthesized using similar procedures (**Scheme S2**) and is an ideal control for live cell imaging of H_2_S.

Finally, to evaluate bioorthogonal click and release H_2_S generation in living cells, we exposed HeLa cells to mitochondria-targeted PPh_3_-PyrS2 or control PPh_3_-Pyr2, with or without BCN, and imaged the production of H_2_S using the fluorescent indicator NDI (**Figure 6**). In the controls lacking BCN or with PPh_3_-Pyr2 instead of PPh_3_-PyrS2, we observed minimal fluorescence in the cells (**Figure 6A, B**). However, living HeLa cells exposed to PPh_3_-PyrS2 and BCN showed significant fluorescence turn on of the H_2_S fluorescent indicator NDI (**Figure 6C**), and the NDI fluorescence colocalized with a fluorescent marker for the mitochondria (Mitotracker Red), indicating we have successfully delivered H_2_S in the mitochondria.

**Figure 6.**
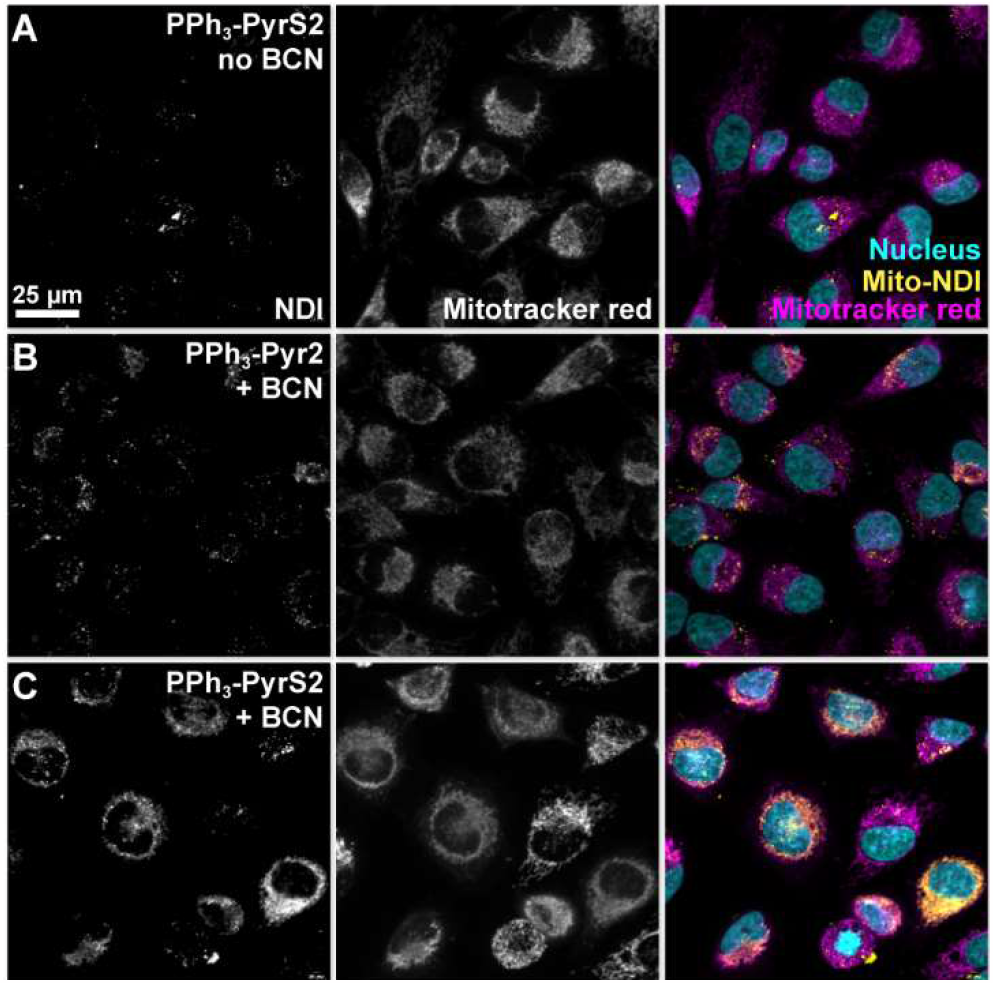
Live cell imaging of H_2_S delivery in mitochondria. HeLa cells were treated with PPh_3_-Pyr2 (40 μM) and NDI (mitochondrial targeting H_2_S probe, 10 μM) for 30 min, followed by treatment with BCN (200 μM) for 2.5 h, washed, co-stained by MitoTracker Red (200 nM) for 21 min, Hoechst 33342 (1 μg/mL) for 15 min at 37 oC, washed, and then imaged by confocal microscopy.

In conclusion, we have reported a “click and release” pair based on an inversed electron demand Diels-Alder cycloaddition between pyrone/pyranthione and BCN by introducing strong electron withdrawing groups on dienes (pyrones/pyranthiones). We found that this type of reaction is likely driven by aromatization induced retro Diels-Alder reaction with the commitment release of CO_2_/COS, which proceeded with second order rate constants comparable to SPAACs. We demonstrated the utility of pyrones/pyranthiones in protein labeling, *in vitro* H_2_S release with quantitative efficiency, and H_2_S delivery to mitochondria in living cells. This click and release reaction enriches the click reaction toolkits and provides a tunable and high efficiency method for subcellular H_2_S delivery in living cells. It should be noted that, during the preparation of this manuscript, a similar strategy^[26]^ for H_2_S generation, albeit in fixed cells, was reported.

## Notes

### Competing Interest Statement

The authors have declared no competing interest.

