## Supporting Information for "Pyranthiones/Pyrones: “Click and Release” Donors for Subcellular Hydrogen Sulfide Delivery and Labeling"

**Table of Contents**

List of abbreviations ..... S3

General materials and methods ..... S4

Computational details ..... S5

Synthetic details and characterization ..... S30

Scheme S1 ..... S34

Scheme S2 ..... S36

Scheme S3 ..... S37

Scheme S4 ..... S38

Kinetics experimental procedures, pyrones and BCN ..... S39

Figure S1 ..... S40

Kinetics experimental procedures, pyranthiones and BCN ..... S41

Figure S2 ..... S42

Figure S3 ..... S42

Figure S4 ..... S43

Bioorthogonality evaluation ..... S44

Figure S5 ..... S44

Figure S6 ..... S44

Protein labeling experiments ..... S45

Figure S7 ..... S45

Figure S8 ..... S46

Fluorsecent assay of $H_2S$ release ..... S47

Figure S9 ..... S47

Figure S10 ..... S47

Figure S11 ..... S48

Quantification of $H_2S$ release by methylene blue assay ..... S49

Figure S12 ..... S49

Figure S13 ..... S49

Time dependent $H_2S$ release from click reaction between pyranthiones and BCN by MB assay ..... S50

HeLa cell culture and live imaging ..... S51

NMR spectra ..... S52

References ..... S93

**List of Abbreviations**

| | |
|---|---|
| BCN | (1*R*,8*S*,9*s*)-Bicyclo[6.1.0]non-4-yn-9-ylmethanol |
| Calcd | Calculated |
| $CDCl_3$ | Deuterated chloroform |
| DCM | Dichloromethane |
| DIPEA | *N,N*-diisopropylethylamine |
| DMF | Dimethylformamide |
| DMEM | Dulbecco's modified eagle medium |
| DMSO | Dimethyl sulfoxide |
| ESI | Electrospray ionization |
| EtOAc | Ethyl acetate |
| HPLC | High performance liquid chromatography |
| HRMS | High resolution mass spectrometry |
| MeCN | Acetonitrile |
| MeOH | Methanol |
| MHz | Megahertz |
| PBS | Phosphate-buffered saline |
| THF | Tetrahydrofuran |
| TLC | Thin layer chromatography |

**General materials and methods.**

All chemical reagents were of analytical grade, obtained from commercial suppliers, and used without further purification unless otherwise specified. Reactions were monitored by thin layer chromatography on pre-coated glass TLC plates (Analtech UNIPLATE™ silica gel HLF w/ organic binder, 250 μm thickness, with UV254 indicator) or by LC/MS (Agilent 6110 Single Quad, LC-MS, direct-injection mode, ESI). TLC plates were visualized by UV illumination or developed with potassium permanganate stain. NMR spectra ($^1$H and $^{13}$C) were obtained using a 400, 500, or 700 MHz Bruker spectrometer and analyzed using TopSpin 4.2.0. $^1$H and $^{13}$C chemical shifts (δ) were referenced to residual solvent peaks. The following residual solvent peaks were chosen: (for $^1$H NMR) $CDCl_3$, 7.26 ppm; $CD_3CN$, 1.94 ppm; (for $^{13}$C NMR) $CDCl_3$, 77.16 ppm; $CD_3CN$, 118.26 ppm. The following abbreviations were used to define $^1$H NMR peaks: s, singlet; d, doublet; t, triplet; q, quartet; m, multiplet. Low-resolution electrospray ionization (ESI) and High-resolution electrospray ionization (ESI) mass spectra were obtained at CASDA Mass Spectrometry Center at Stony Brook University. The absorption spectrum and kinetics were measured on a Biotek EPOCH 2 microplate reader. Pyrones **Pyr1**[1], **Pyr2**[2], **Pyr3**[3], **Pyr4**[4] were synthesized according to previous reports. **BCN** was either synthesized according to previous report[5] or purchased from sigma-aldrich (CAS# 1263166-90-0, 742678). 35mm glass bottom microwell dish (20 mm glass diameter) were purchased from MatTek corporation (Ashland, MA). HeLa cell line was a gift from Professor Peter Tonge's lab (ATCC CCL-2TM). MitoTracker Red (Invitrogen, catlog# M7512) and Hoechst 33342 were purchased from Thermo Fisher Scientific (Boston, MA). All cell images were acquired from the confocal microscope (Zeiss Axio Examiner. D1 modified with an Andor Differential Scanning Disk confocal unit) using a Zeiss water immersion 40X NA 1 Plan-APOCHROMAT and analyzed using ImageJ to produce maximum intensity Z projections.

**Computational details.**

All DFT calculations were performed using Gaussian16 Rev.B01.[6] The Minnesota functional M06-2X14 with the 6-31G(d) basis was used for geometry optimization of minima and transition states. A frequency analysis was performed using the same level of theory to confirm the presence of a minima with no imaginary frequency or a transition state with a single imaginary frequency. Next, single point energy calculations were performed at the M06-2X[7] with the 6-311+G(d,p) basis set including the solvent with a polarizable continuum model (PCM).[8] Intrinsic reaction coordinate (IRC) calculations were performed to verify the expected connections of the first order saddle points with the local minima found on the potential energy surface.[9]

For the activation strain model (ASM),[10] Every geometry of the IRC was separated into BCN and pyrone or pyranthione. On one hand, the distortion energy was calculated as the energy difference between the energy of the distorted and the fully optimized counterpart. On the other hand, the interaction energy was calculated as the energy difference between the energy of the geometry of the reaction coordinate and the energy of the total distortion energy. The distortion, interaction, and total energies were plotted versus the C–C bond forming distance, which was used to represent the reaction coordinate. Optimized structures were illustrated using CYLview20.3.[11]

**Cartesian coordinates of optimized structures.**

**BCN**

Zero-point correction= 0.220292 (Hartree/Particle)
Thermal correction to Energy= 0.230755
Thermal correction to Enthalpy= 0.231700
Thermal correction to Gibbs Free Energy= 0.184790
Sum of electronic and zero-point Energies= -464.323839
Sum of electronic and thermal Energies= -464.313375
Sum of electronic and thermal Enthalpies= -464.312431
Sum of electronic and thermal Free Energies= -464.359341

| | | | |
|---|---|---|---|
| 6 | -2.516811000 | -0.153969000 | -0.258576000 |
| 6 | -2.241321000 | 1.018681000 | -0.202407000 |
| 6 | -2.208203000 | -1.587597000 | -0.265170000 |
| 1 | -2.566198000 | -2.068275000 | 0.649297000 |
| 1 | -2.662869000 | -2.106772000 | -1.110871000 |
| 6 | -1.336682000 | 2.165009000 | -0.069109000 |
| 1 | -1.482420000 | 2.664377000 | 0.892568000 |
| 1 | -1.489316000 | 2.908817000 | -0.853316000 |
| 6 | 0.090658000 | 1.566625000 | -0.160440000 |
| 1 | 0.220092000 | 1.150037000 | -1.162035000 |
| 1 | 0.817523000 | 2.377338000 | -0.048371000 |
| 6 | -0.662583000 | -1.660675000 | -0.351433000 |
| 1 | -0.359319000 | -1.225148000 | -1.305602000 |
| 1 | -0.359615000 | -2.712844000 | -0.359573000 |
| 6 | 0.014896000 | -0.959553000 | 0.814634000 |
| 6 | 0.352673000 | 0.518075000 | 0.906978000 |
| 6 | 1.456905000 | -0.508801000 | 0.785334000 |
| 1 | 2.034691000 | -0.701198000 | 1.683720000 |
| 1 | 0.233734000 | 0.918444000 | 1.910282000 |
| 1 | -0.286544000 | -1.383297000 | 1.768953000 |
| 6 | 2.274525000 | -0.610816000 | -0.469541000 |
| 1 | 1.664052000 | -0.474601000 | -1.364712000 |
| 1 | 2.746856000 | -1.597093000 | -0.529682000 |
| 8 | 3.286820000 | 0.403507000 | -0.419798000 |
| 1 | 3.850431000 | 0.300290000 | -1.193892000 |

**Pyr1**

Zero-point correction= 0.209970 (Hartree/Particle)
Thermal correction to Energy= 0.228650
Thermal correction to Enthalpy= 0.229594
Thermal correction to Gibbs Free Energy= 0.161523
Sum of electronic and zero-point Energies= -1026.699187
Sum of electronic and thermal Energies= -1026.680507
Sum of electronic and thermal Enthalpies= -1026.679563
Sum of electronic and thermal Free Energies= -1026.747634

| | | | |
|---|---|---|---|
| 6 | -0.071718000 | 0.168090000 | -0.080985000 |
| 6 | -1.336240000 | 0.629530000 | -0.061950000 |
| 6 | -0.665634000 | 2.920532000 | 0.096251000 |
| 6 | 0.706167000 | 2.454899000 | 0.139323000 |
| 6 | 0.989753000 | 1.141563000 | 0.043727000 |
| 8 | -1.629500000 | 1.941438000 | 0.005860000 |
| 1 | 1.479165000 | 3.206165000 | 0.235308000 |
| 8 | -1.038686000 | 4.065377000 | 0.137508000 |
| 6 | -2.591079000 | -0.192518000 | -0.109715000 |
| 8 | -3.658485000 | 0.273832000 | -0.415649000 |

| | | | |
|---|---|---|---|
| 8 | -2.365294000 | -1.443125000 | 0.233404000 |
| 6 | -3.491355000 | -2.340944000 | 0.153670000 |
| 1 | -4.269905000 | -2.010121000 | 0.839496000 |
| 1 | -3.865903000 | -2.364749000 | -0.868743000 |
| 1 | -3.104827000 | -3.311987000 | 0.447323000 |
| 6 | 0.242055000 | -1.296991000 | -0.221897000 |
| 8 | 0.108162000 | -1.914133000 | -1.247599000 |
| 8 | 0.717749000 | -1.788616000 | 0.909095000 |
| 6 | 1.197294000 | -3.146290000 | 0.847006000 |
| 1 | 0.377111000 | -3.816020000 | 0.591077000 |
| 1 | 1.992454000 | -3.215561000 | 0.104035000 |
| 1 | 1.577167000 | -3.365198000 | 1.840385000 |
| 6 | 2.437024000 | 0.729956000 | 0.070144000 |
| 8 | 3.283802000 | 1.339935000 | 0.673463000 |
| 8 | 2.648567000 | -0.355489000 | -0.649551000 |
| 6 | 3.984164000 | -0.896635000 | -0.623267000 |
| 1 | 4.688219000 | -0.155709000 | -0.999136000 |
| 1 | 4.237066000 | -1.181889000 | 0.397556000 |
| 1 | 3.956350000 | -1.765837000 | -1.273383000 |

**Pyr2**

Zero-point correction= 0.167072 (Hartree/Particle)
Thermal correction to Energy= 0.181223
Thermal correction to Enthalpy= 0.182167
Thermal correction to Gibbs Free Energy= 0.124338
Sum of electronic and zero-point Energies= -798.889873
Sum of electronic and thermal Energies= -798.875722
Sum of electronic and thermal Enthalpies= -798.874778
Sum of electronic and thermal Free Energies= -798.932607

| | | | |
|---|---|---|---|
| 6 | -0.145260000 | -0.565977000 | -0.000101000 |
| 6 | -1.212213000 | 0.246159000 | -0.000068000 |
| 6 | 0.109249000 | 2.233788000 | -0.000271000 |
| 6 | 1.286183000 | 1.383643000 | -0.000087000 |
| 6 | 1.155382000 | 0.040615000 | -0.000086000 |
| 8 | -1.105753000 | 1.595077000 | -0.000103000 |
| 1 | 2.244569000 | 1.883940000 | 0.000005000 |
| 8 | 0.104141000 | 3.441825000 | 0.000116000 |
| 6 | -2.640284000 | -0.191817000 | -0.000011000 |
| 8 | -3.563842000 | 0.584870000 | 0.000049000 |
| 8 | -2.750497000 | -1.507470000 | 0.000078000 |
| 6 | -4.094255000 | -2.027239000 | 0.000194000 |
| 1 | -4.617551000 | -1.692533000 | 0.894955000 |
| 1 | -4.617666000 | -1.692635000 | -0.894539000 |
| 1 | -3.984515000 | -3.107325000 | 0.000248000 |
| 6 | 2.347264000 | -0.871402000 | -0.000016000 |
| 8 | 2.243070000 | -2.074990000 | 0.000051000 |
| 8 | 3.494810000 | -0.217821000 | -0.000011000 |
| 6 | 4.684365000 | -1.029953000 | 0.000066000 |
| 1 | 4.704853000 | -1.651367000 | 0.894467000 |
| 1 | 4.704912000 | -1.651454000 | -0.894273000 |
| 1 | 5.512998000 | -0.328301000 | 0.000059000 |
| 1 | -0.265620000 | -1.639148000 | -0.000080000 |

**Pyr3**

Zero-point correction= 0.223005 (Hartree/Particle)
Thermal correction to Energy= 0.239973
Thermal correction to Enthalpy= 0.240917

Thermal correction to Gibbs Free Energy= 0.177853
Sum of electronic and zero-point Energies= -877.455468
Sum of electronic and thermal Energies= -877.438501
Sum of electronic and thermal Enthalpies= -877.437557
Sum of electronic and thermal Free Energies= -877.500620

| | | | |
|---|---|---|---|
| 6 | -0.657510000 | 0.576651000 | 0.122501000 |
| 6 | -1.849078000 | 1.225700000 | 0.107565000 |
| 6 | -3.052925000 | -0.830457000 | -0.301289000 |
| 6 | -1.793633000 | -1.532473000 | -0.284043000 |
| 6 | -0.651493000 | -0.848137000 | -0.067224000 |
| 8 | -2.993102000 | 0.532664000 | -0.083657000 |
| 1 | -1.817414000 | -2.606227000 | -0.412435000 |
| 8 | -4.150203000 | -1.307894000 | -0.472493000 |
| 6 | 0.621808000 | 1.319449000 | 0.264630000 |
| 8 | 0.823756000 | 2.214471000 | 1.052480000 |
| 8 | 1.527195000 | 0.860959000 | -0.593943000 |
| 6 | 2.891383000 | 1.315839000 | -0.416552000 |
| 1 | 2.906936000 | 2.402981000 | -0.498527000 |
| 1 | 3.206694000 | 1.027123000 | 0.589062000 |
| 6 | 0.632366000 | -1.620993000 | 0.058094000 |
| 8 | 0.993714000 | -2.459041000 | -0.729887000 |
| 8 | 1.281090000 | -1.282155000 | 1.161186000 |
| 6 | 2.567116000 | -1.901171000 | 1.358678000 |
| 1 | 2.446540000 | -2.979390000 | 1.456020000 |
| 1 | 3.218109000 | -1.665651000 | 0.516770000 |
| 1 | 2.955745000 | -1.473741000 | 2.278142000 |
| 6 | -2.101548000 | 2.679819000 | 0.266299000 |
| 1 | -1.207466000 | 3.262473000 | 0.060024000 |
| 1 | -2.905399000 | 2.976649000 | -0.407754000 |
| 1 | -2.423117000 | 2.877672000 | 1.292802000 |
| 6 | 3.719791000 | 0.653897000 | -1.488640000 |
| 1 | 3.386313000 | 0.964534000 | -2.480274000 |
| 1 | 4.764190000 | 0.946102000 | -1.367142000 |
| 1 | 3.651613000 | -0.433300000 | -1.416283000 |

**Pyr4**

Zero-point correction= 0.247438 (Hartree/Particle)
Thermal correction to Energy= 0.266000
Thermal correction to Enthalpy= 0.266944
Thermal correction to Gibbs Free Energy= 0.197154
Sum of electronic and zero-point Energies= -1029.828978
Sum of electronic and thermal Energies= -1029.810415
Sum of electronic and thermal Enthalpies= -1029.809471
Sum of electronic and thermal Free Energies= -1029.879261

| | | | |
|---|---|---|---|
| 6 | 0.601645000 | -0.703556000 | -0.328667000 |
| 6 | 0.528821000 | 0.621319000 | -0.123361000 |
| 6 | 2.890079000 | 0.930886000 | -0.261312000 |
| 6 | 2.983852000 | -0.507991000 | -0.465064000 |
| 6 | 1.893984000 | -1.294978000 | -0.502711000 |
| 8 | 1.627586000 | 1.414269000 | -0.084178000 |
| 8 | 3.816721000 | 1.704914000 | -0.243282000 |
| 6 | -0.719328000 | 1.410584000 | 0.084243000 |
| 8 | -0.719130000 | 2.602866000 | 0.272761000 |
| 8 | -1.797478000 | 0.646402000 | 0.035454000 |
| 6 | -3.067796000 | 1.326295000 | 0.234901000 |
| 1 | -3.045703000 | 1.796060000 | 1.218775000 |

| | | | |
|---|---|---|---|
| 1 | -3.167773000 | 2.089684000 | -0.537049000 |
| 8 | 4.248109000 | -1.001683000 | -0.695422000 |
| 1 | -0.296590000 | -1.302401000 | -0.353576000 |
| 1 | 1.999012000 | -2.360613000 | -0.666339000 |
| 6 | -4.144071000 | 0.288090000 | 0.135070000 |
| 6 | -4.776054000 | 0.045910000 | -1.084196000 |
| 6 | -4.499979000 | -0.462127000 | 1.256510000 |
| 6 | -5.754584000 | -0.939255000 | -1.183252000 |
| 6 | -5.476302000 | -1.448486000 | 1.158611000 |
| 6 | -6.104216000 | -1.687430000 | -0.062009000 |
| 1 | -4.501378000 | 0.633394000 | -1.954104000 |
| 1 | -4.011312000 | -0.268015000 | 2.205850000 |
| 1 | -6.244901000 | -1.120714000 | -2.132512000 |
| 1 | -5.750533000 | -2.026514000 | 2.033303000 |
| 1 | -6.867626000 | -2.453048000 | -0.138013000 |
| 6 | 5.128265000 | -0.958822000 | 0.360478000 |
| 8 | 4.772478000 | -0.620516000 | 1.456238000 |
| 6 | 6.490647000 | -1.377108000 | -0.063934000 |
| 1 | 6.435063000 | -2.339762000 | -0.573436000 |
| 1 | 6.875410000 | -0.640254000 | -0.772203000 |
| 1 | 7.140255000 | -1.437806000 | 0.804903000 |

**PyrS1**

Zero-point correction= 0.207587 (Hartree/Particle)
Thermal correction to Energy= 0.226680
Thermal correction to Enthalpy= 0.227624
Thermal correction to Gibbs Free Energy= 0.158348
Sum of electronic and zero-point Energies= -1349.644583
Sum of electronic and thermal Energies= -1349.625490
Sum of electronic and thermal Enthalpies= -1349.624546
Sum of electronic and thermal Free Energies= -1349.693822

| | | | |
|---|---|---|---|
| 6 | -0.074765000 | 0.071615000 | -0.085710000 |
| 6 | 1.181568000 | 0.558554000 | -0.070211000 |
| 6 | 2.194147000 | -1.593674000 | 0.068422000 |
| 6 | 0.871316000 | -2.149340000 | 0.116757000 |
| 6 | -0.223012000 | -1.358284000 | 0.032063000 |
| 8 | 2.266452000 | -0.236266000 | -0.008586000 |
| 1 | 0.776054000 | -3.223122000 | 0.205638000 |
| 16 | 3.594141000 | -2.450734000 | 0.105090000 |
| 6 | 1.590350000 | 2.001795000 | -0.118541000 |
| 8 | 2.708361000 | 2.348227000 | -0.401249000 |
| 8 | 0.590650000 | 2.798361000 | 0.194685000 |
| 6 | 0.855272000 | 4.214747000 | 0.121011000 |
| 1 | 1.637740000 | 4.475480000 | 0.832072000 |
| 1 | 1.155512000 | 4.476061000 | -0.892788000 |
| 1 | -0.082098000 | 4.695323000 | 0.383675000 |
| 6 | -1.274199000 | 0.973036000 | -0.220198000 |
| 8 | -1.602188000 | 1.493385000 | -1.255446000 |
| 8 | -1.917516000 | 1.077748000 | 0.928336000 |
| 6 | -3.156243000 | 1.813641000 | 0.885277000 |
| 1 | -2.963749000 | 2.841870000 | 0.581047000 |
| 1 | -3.840311000 | 1.330466000 | 0.187135000 |
| 1 | -3.549098000 | 1.778588000 | 1.896819000 |
| 6 | -1.569899000 | -2.025505000 | 0.053469000 |
| 8 | -1.767042000 | -3.092523000 | 0.579863000 |
| 8 | -2.480687000 | -1.312763000 | -0.581278000 |

| | | | |
|---|---|---|---|
| 6 | -3.825630000 | -1.831035000 | -0.577787000 |
| 1 | -3.841318000 | -2.811751000 | -1.051047000 |
| 1 | -4.185904000 | -1.896089000 | 0.448285000 |
| 1 | -4.410759000 | -1.117732000 | -1.150201000 |

**PyrS2**

Zero-point correction= 0.164665 (Hartree/Particle)
Thermal correction to Energy= 0.179253
Thermal correction to Enthalpy= 0.180197
Thermal correction to Gibbs Free Energy= 0.121056
Sum of electronic and zero-point Energies= -1121.835715
Sum of electronic and thermal Energies= -1121.821127
Sum of electronic and thermal Enthalpies= -1121.820183
Sum of electronic and thermal Free Energies= -1121.879323

| | | | |
|---|---|---|---|
| 6 | -0.161089000 | -0.865725000 | -0.000203000 |
| 6 | -1.218419000 | -0.038706000 | -0.000025000 |
| 6 | 0.117849000 | 1.927832000 | -0.000113000 |
| 6 | 1.276174000 | 1.076482000 | -0.000257000 |
| 6 | 1.140769000 | -0.272162000 | -0.000320000 |
| 8 | -1.090315000 | 1.307336000 | 0.000028000 |
| 1 | 2.245335000 | 1.554653000 | -0.000319000 |
| 16 | 0.150640000 | 3.574299000 | 0.000127000 |
| 6 | -2.654496000 | -0.450273000 | 0.000126000 |
| 8 | -3.563567000 | 0.342652000 | 0.000338000 |
| 8 | -2.785175000 | -1.763458000 | 0.000135000 |
| 6 | -4.137267000 | -2.261833000 | 0.000340000 |
| 1 | -4.654702000 | -1.918573000 | 0.895250000 |
| 1 | -4.654980000 | -1.918553000 | -0.894401000 |
| 1 | -4.044546000 | -3.343469000 | 0.000314000 |
| 6 | 2.326022000 | -1.189366000 | -0.000523000 |
| 8 | 2.215028000 | -2.392644000 | -0.000138000 |
| 8 | 3.477557000 | -0.542216000 | -0.000183000 |
| 6 | 4.662393000 | -1.361097000 | 0.000169000 |
| 1 | 4.679200000 | -1.982690000 | 0.894536000 |
| 1 | 4.679736000 | -1.982678000 | -0.894197000 |
| 1 | 5.495024000 | -0.664180000 | 0.000424000 |
| 1 | -0.295145000 | -1.937554000 | -0.000246000 |

**PyrS3**

Zero-point correction= 0.220619 (Hartree/Particle)
Thermal correction to Energy= 0.237999
Thermal correction to Enthalpy= 0.238944
Thermal correction to Gibbs Free Energy= 0.174530
Sum of electronic and zero-point Energies= -1200.401200
Sum of electronic and thermal Energies= -1200.383819
Sum of electronic and thermal Enthalpies= -1200.382875
Sum of electronic and thermal Free Energies= -1200.447288

| | | | |
|---|---|---|---|
| 6 | -0.271738000 | 0.713840000 | 0.152378000 |
| 6 | -1.377845000 | 1.501721000 | 0.147198000 |
| 6 | -2.830741000 | -0.390407000 | -0.172175000 |
| 6 | -1.680056000 | -1.235796000 | -0.167031000 |
| 6 | -0.446800000 | -0.699395000 | 0.001062000 |
| 8 | -2.599540000 | 0.945516000 | -0.001751000 |
| 1 | -1.824796000 | -2.302734000 | -0.269218000 |
| 16 | -4.394709000 | -0.880569000 | -0.353026000 |
| 6 | 1.092023000 | 1.300664000 | 0.252827000 |
| 8 | 1.417385000 | 2.166646000 | 1.030955000 |
| 8 | 1.910093000 | 0.735500000 | -0.627079000 |

| | | | |
|---|---|---|---|
| 6 | 3.323799000 | 1.024335000 | -0.487958000 |
| 1 | 3.466624000 | 2.101521000 | -0.577647000 |
| 1 | 3.627570000 | 0.703396000 | 0.511419000 |
| 6 | 0.730830000 | -1.626558000 | 0.104791000 |
| 8 | 0.943813000 | -2.530044000 | -0.665062000 |
| 8 | 1.469457000 | -1.337774000 | 1.164445000 |
| 6 | 2.676462000 | -2.107302000 | 1.329358000 |
| 1 | 2.428077000 | -3.159193000 | 1.464838000 |
| 1 | 3.313867000 | -1.977012000 | 0.454734000 |
| 6 | -1.454068000 | 2.977206000 | 0.281267000 |
| 1 | -0.502362000 | 3.445170000 | 0.042800000 |
| 1 | -2.234063000 | 3.356090000 | -0.379506000 |
| 1 | -1.722939000 | 3.228853000 | 1.311182000 |
| 6 | 4.038706000 | 0.263684000 | -1.575739000 |
| 1 | 5.113215000 | 0.430250000 | -1.483105000 |
| 1 | 3.843793000 | -0.807285000 | -1.492724000 |
| 1 | 3.718473000 | 0.607158000 | -2.560855000 |
| 1 | 3.154793000 | -1.707821000 | 2.218574000 |

**PyrS4**

Zero-point correction= 0.244996 (Hartree/Particle)
Thermal correction to Energy= 0.263942
Thermal correction to Enthalpy= 0.264886
Thermal correction to Gibbs Free Energy= 0.193920
Sum of electronic and zero-point Energies= -1352.775282
Sum of electronic and thermal Energies= -1352.756337
Sum of electronic and thermal Enthalpies= -1352.755393
Sum of electronic and thermal Free Energies= -1352.826359

| | | | |
|---|---|---|---|
| 6 | 0.368550000 | -0.790227000 | -0.349268000 |
| 6 | 0.325901000 | 0.530358000 | -0.095772000 |
| 6 | 2.688388000 | 0.800639000 | -0.220123000 |
| 6 | 2.754500000 | -0.615315000 | -0.484835000 |
| 6 | 1.646580000 | -1.387970000 | -0.548416000 |
| 8 | 1.439753000 | 1.290600000 | -0.033007000 |
| 16 | 3.970920000 | 1.830153000 | -0.140183000 |
| 6 | -0.901423000 | 1.343677000 | 0.150953000 |
| 8 | -0.868839000 | 2.525773000 | 0.389791000 |
| 8 | -1.996091000 | 0.607546000 | 0.074740000 |
| 6 | -3.250405000 | 1.306270000 | 0.310813000 |
| 1 | -3.217995000 | 1.720370000 | 1.319058000 |
| 1 | -3.329831000 | 2.112995000 | -0.418239000 |
| 8 | 3.996280000 | -1.151284000 | -0.738238000 |
| 1 | -0.540967000 | -1.370981000 | -0.391913000 |
| 1 | 1.738933000 | -2.447562000 | -0.753527000 |
| 6 | -4.350848000 | 0.301461000 | 0.152680000 |
| 6 | -4.968102000 | 0.125858000 | -1.085534000 |
| 6 | -4.743978000 | -0.484581000 | 1.236217000 |
| 6 | -5.969004000 | -0.828921000 | -1.240769000 |
| 6 | -5.743273000 | -1.440477000 | 1.082211000 |
| 6 | -6.356126000 | -1.613110000 | -0.156948000 |
| 1 | -4.664270000 | 0.741625000 | -1.925701000 |
| 1 | -4.266435000 | -0.342529000 | 2.200239000 |
| 1 | -6.447766000 | -0.958769000 | -2.204262000 |
| 1 | -6.046554000 | -2.046766000 | 1.927693000 |
| 1 | -7.137171000 | -2.354991000 | -0.276505000 |
| 6 | 4.814144000 | -1.361204000 | 0.347020000 |
| 8 | 4.408983000 | -1.238871000 | 1.470459000 |

| | | | |
|---|---|---|---|
| 6 | 6.184156000 | -1.742653000 | -0.086920000 |
| 1 | 6.128743000 | -2.547995000 | -0.819995000 |
| 1 | 6.643255000 | -0.875824000 | -0.568411000 |
| 1 | 6.770280000 | -2.044974000 | 0.776684000 |

**TS-Pyr1-BCN**

Frequency -337.6940
Zero-point correction= 0.431279 (Hartree/Particle)
Thermal correction to Energy= 0.461144
Thermal correction to Enthalpy= 0.462088
Thermal correction to Gibbs Free Energy= 0.369884
Sum of electronic and zero-point Energies= -1491.019961
Sum of electronic and thermal Energies= -1490.990097
Sum of electronic and thermal Enthalpies= -1490.989153
Sum of electronic and thermal Free Energies= -1491.081356

| | | | |
|---|---|---|---|
| 6 | -0.932857000 | -0.809858000 | 1.008033000 |
| 6 | -0.871828000 | 0.341156000 | 0.564657000 |
| 6 | -1.717052000 | -2.042034000 | 1.248935000 |
| 1 | -1.261890000 | -2.876747000 | 0.706717000 |
| 1 | -1.698325000 | -2.297940000 | 2.311691000 |
| 6 | -1.470766000 | 1.473808000 | -0.166520000 |
| 1 | -0.922008000 | 1.639340000 | -1.099042000 |
| 1 | -1.408169000 | 2.396346000 | 0.417917000 |
| 6 | -2.948781000 | 1.147399000 | -0.461120000 |
| 1 | -3.482904000 | 1.065916000 | 0.487694000 |
| 1 | -3.377418000 | 1.997191000 | -1.001156000 |
| 6 | -3.161655000 | -1.807781000 | 0.774269000 |
| 1 | -3.594815000 | -1.003589000 | 1.371768000 |
| 1 | -3.740782000 | -2.713203000 | 0.980639000 |
| 6 | -3.221007000 | -1.499168000 | -0.708753000 |
| 6 | -3.106909000 | -0.107652000 | -1.293475000 |
| 6 | -4.416601000 | -0.858112000 | -1.371477000 |
| 1 | -4.678065000 | -1.249331000 | -2.349217000 |
| 1 | -2.558918000 | -0.062560000 | -2.230373000 |
| 1 | -2.745390000 | -2.269924000 | -1.308559000 |
| 6 | -5.611725000 | -0.415336000 | -0.577706000 |
| 1 | -5.340836000 | -0.116258000 | 0.436895000 |
| 1 | -6.334905000 | -1.234353000 | -0.507275000 |
| 8 | -6.209308000 | 0.694148000 | -1.262388000 |
| 1 | -7.018663000 | 0.933407000 | -0.797529000 |
| 6 | 1.979226000 | 0.189804000 | 0.363917000 |
| 6 | 1.303501000 | 1.071913000 | 1.184538000 |
| 6 | 0.886555000 | -0.526974000 | 2.857454000 |
| 6 | 1.059382000 | -1.490273000 | 1.779673000 |
| 6 | 1.877273000 | -1.162517000 | 0.711978000 |
| 8 | 1.098609000 | 0.770133000 | 2.501238000 |
| 8 | 0.593513000 | -0.771049000 | 4.004257000 |
| 1 | 0.856912000 | -2.524268000 | 2.029685000 |
| 6 | 2.573907000 | 0.613749000 | -0.950813000 |
| 8 | 3.634792000 | 1.175327000 | -1.056817000 |
| 8 | 1.798960000 | 0.252741000 | -1.960865000 |
| 6 | 2.333686000 | 0.496278000 | -3.276522000 |
| 1 | 2.484238000 | 1.565629000 | -3.421870000 |
| 1 | 3.276210000 | -0.039801000 | -3.390891000 |
| 1 | 1.587763000 | 0.115734000 | -3.967913000 |
| 6 | 2.358188000 | -2.255540000 | -0.179313000 |
| 8 | 1.830806000 | -3.341279000 | -0.246360000 |

| | | | |
|---|---|---|---|
| 8 | 3.418306000 | -1.889496000 | -0.883836000 |
| 6 | 3.883875000 | -2.823033000 | -1.876262000 |
| 1 | 4.169438000 | -3.759091000 | -1.398149000 |
| 1 | 3.097039000 | -2.994473000 | -2.610719000 |
| 1 | 4.745380000 | -2.349514000 | -2.337631000 |
| 6 | 1.140917000 | 2.538671000 | 0.946796000 |
| 8 | 0.850696000 | 3.319400000 | 1.819295000 |
| 8 | 1.308050000 | 2.840331000 | -0.325462000 |
| 6 | 1.096659000 | 4.218335000 | -0.689522000 |
| 1 | 0.069336000 | 4.500505000 | -0.460036000 |
| 1 | 1.797247000 | 4.853278000 | -0.149023000 |
| 1 | 1.280210000 | 4.264649000 | -1.758716000 |

**TS-Pyr2-BCN**

Frequency -350.3912
Zero-point correction= 0.388364 (Hartree/Particle)
Thermal correction to Energy= 0.413663
Thermal correction to Enthalpy= 0.414607
Thermal correction to Gibbs Free Energy= 0.332428
Sum of electronic and zero-point Energies= -1263.206622
Sum of electronic and thermal Energies= -1263.181324
Sum of electronic and thermal Enthalpies= -1263.180379
Sum of electronic and thermal Free Energies= -1263.262559

| | | | |
|---|---|---|---|
| 6 | -0.378944000 | -0.670495000 | 0.838444000 |
| 6 | -0.464007000 | 0.425167000 | 0.272853000 |
| 6 | -1.042660000 | -1.904565000 | 1.316695000 |
| 1 | -0.615459000 | -2.771583000 | 0.802821000 |
| 1 | -0.857534000 | -2.040680000 | 2.385689000 |
| 6 | -1.238027000 | 1.436759000 | -0.471040000 |
| 1 | -0.837595000 | 1.531460000 | -1.485633000 |
| 1 | -1.156867000 | 2.418684000 | 0.004022000 |
| 6 | -2.718283000 | 1.006671000 | -0.512935000 |
| 1 | -3.109333000 | 1.006447000 | 0.506575000 |
| 1 | -3.272603000 | 1.768077000 | -1.070263000 |
| 6 | -2.551747000 | -1.797229000 | 1.040005000 |
| 1 | -2.946036000 | -0.951482000 | 1.606340000 |
| 1 | -3.037069000 | -2.699806000 | 1.424400000 |
| 6 | -2.843950000 | -1.661422000 | -0.441490000 |
| 6 | -2.911128000 | -0.340315000 | -1.177294000 |
| 6 | -4.164356000 | -1.161380000 | -0.977307000 |
| 1 | -4.538565000 | -1.672164000 | -1.858480000 |
| 1 | -2.510798000 | -0.372681000 | -2.186574000 |
| 1 | -2.409291000 | -2.469383000 | -1.023136000 |
| 6 | -5.257268000 | -0.696934000 | -0.058288000 |
| 1 | -4.860933000 | -0.279318000 | 0.869456000 |
| 1 | -5.906972000 | -1.539346000 | 0.200752000 |
| 8 | -6.020450000 | 0.302509000 | -0.747763000 |
| 1 | -6.753852000 | 0.562947000 | -0.179802000 |
| 6 | 2.324658000 | 0.420491000 | -0.317126000 |
| 6 | 1.696391000 | 1.345102000 | 0.488584000 |
| 6 | 1.620467000 | -0.063436000 | 2.363167000 |
| 6 | 1.740452000 | -1.133506000 | 1.383494000 |
| 6 | 2.391112000 | -0.876821000 | 0.183920000 |
| 8 | 1.679451000 | 1.194935000 | 1.850859000 |
| 1 | 1.657720000 | -2.140334000 | 1.773160000 |
| 8 | 1.487933000 | -0.194430000 | 3.560266000 |
| 6 | 1.434630000 | 2.759567000 | 0.100090000 |

| | | | |
|---|---|---|---|
| 8 | 1.157971000 | 3.626300000 | 0.894856000 |
| 8 | 1.506091000 | 2.925032000 | -1.209109000 |
| 6 | 1.181426000 | 4.241072000 | -1.695361000 |
| 1 | 0.156466000 | 4.489401000 | -1.420210000 |
| 1 | 1.874106000 | 4.970942000 | -1.278380000 |
| 1 | 1.287106000 | 4.184230000 | -2.774467000 |
| 6 | 2.874514000 | -2.022527000 | -0.630046000 |
| 8 | 2.673617000 | -3.182097000 | -0.342992000 |
| 8 | 3.550178000 | -1.631643000 | -1.702986000 |
| 6 | 4.048750000 | -2.678895000 | -2.554123000 |
| 1 | 4.737801000 | -3.312430000 | -1.996663000 |
| 1 | 3.218591000 | -3.270548000 | -2.938387000 |
| 1 | 4.565017000 | -2.172677000 | -3.364369000 |
| 1 | 2.585590000 | 0.671579000 | -1.335374000 |

**TS-Pyr3-BCN**

Frequency -387.1576

Zero-point correction= 0.444689 (Hartree/Particle)

Thermal correction to Energy= 0.472637

Thermal correction to Enthalpy= 0.473582

Thermal correction to Gibbs Free Energy= 0.385720

Sum of electronic and zero-point Energies= -1341.768788

Sum of electronic and thermal Energies= -1341.740840

Sum of electronic and thermal Enthalpies= -1341.739895

Sum of electronic and thermal Free Energies= -1341.827757

| | | | |
|---|---|---|---|
| 6 | 0.816943000 | 1.269750000 | -0.508328000 |
| 6 | 0.795329000 | 0.905398000 | 0.675829000 |
| 6 | 1.516707000 | 1.411336000 | -1.803052000 |
| 1 | 0.965509000 | 0.870261000 | -2.578650000 |
| 1 | 1.545052000 | 2.462727000 | -2.102480000 |
| 6 | 1.477074000 | 0.188806000 | 1.775650000 |
| 1 | 0.884505000 | -0.692050000 | 2.049018000 |
| 1 | 1.542053000 | 0.817025000 | 2.666531000 |
| 6 | 2.891088000 | -0.232159000 | 1.336344000 |
| 1 | 3.482232000 | 0.668136000 | 1.155445000 |
| 1 | 3.359678000 | -0.759618000 | 2.173238000 |
| 6 | 2.941579000 | 0.847037000 | -1.668838000 |
| 1 | 3.483487000 | 1.439413000 | -0.929446000 |
| 1 | 3.454446000 | 0.976766000 | -2.626966000 |
| 6 | 2.919264000 | -0.623735000 | -1.302039000 |
| 6 | 2.878147000 | -1.135102000 | 0.120979000 |
| 6 | 4.110143000 | -1.352854000 | -0.728061000 |
| 1 | 4.260236000 | -2.364710000 | -1.090060000 |
| 1 | 2.265376000 | -2.021404000 | 0.260361000 |
| 1 | 2.335648000 | -1.214256000 | -2.002736000 |
| 6 | 5.396024000 | -0.639462000 | -0.425574000 |
| 1 | 5.229684000 | 0.407626000 | -0.164504000 |
| 1 | 6.052388000 | -0.669165000 | -1.301451000 |
| 8 | 6.029407000 | -1.313359000 | 0.670715000 |
| 1 | 6.885053000 | -0.897821000 | 0.824257000 |
| 6 | -1.909034000 | 0.615741000 | 0.734187000 |
| 6 | -1.211184000 | 1.531387000 | 1.519032000 |
| 6 | -1.024242000 | 3.088620000 | -0.257291000 |
| 6 | -1.236019000 | 1.944864000 | -1.120715000 |
| 6 | -1.958282000 | 0.864625000 | -0.634468000 |
| 8 | -1.123393000 | 2.832759000 | 1.076821000 |
| 1 | -1.156999000 | 2.132339000 | -2.184270000 |

| | | | |
|---|---|---|---|
| 8 | -0.782420000 | 4.226154000 | -0.605231000 |
| 6 | -2.278556000 | -0.714712000 | 1.304284000 |
| 8 | -3.004231000 | -0.872841000 | 2.258365000 |
| 8 | -1.693818000 | -1.700953000 | 0.636238000 |
| 6 | -2.101228000 | -3.050213000 | 0.973868000 |
| 1 | -1.879828000 | -3.222899000 | 2.027749000 |
| 1 | -3.180470000 | -3.118927000 | 0.818828000 |
| 6 | -2.523871000 | -0.124000000 | -1.591418000 |
| 8 | -2.039355000 | -0.366948000 | -2.673114000 |
| 8 | -3.620736000 | -0.700620000 | -1.114642000 |
| 6 | -4.149500000 | -1.795669000 | -1.884057000 |
| 1 | -4.448369000 | -1.447465000 | -2.872053000 |
| 1 | -5.011612000 | -2.152087000 | -1.327804000 |
| 6 | -1.059740000 | 1.457622000 | 3.000348000 |
| 1 | -0.878730000 | 0.435501000 | 3.328842000 |
| 1 | -0.237466000 | 2.097603000 | 3.318611000 |
| 1 | -1.981623000 | 1.816596000 | 3.466388000 |
| 1 | -3.395025000 | -2.578504000 | -1.970146000 |
| 6 | -1.334150000 | -3.982435000 | 0.070510000 |
| 1 | -1.624712000 | -5.011870000 | 0.286720000 |
| 1 | -1.553194000 | -3.772346000 | -0.978187000 |
| 1 | -0.259905000 | -3.883482000 | 0.236386000 |

**TS-Pyr5-BCN**

Frequency -396.9694
Zero-point correction= 0.468882 (Hartree/Particle)
Thermal correction to Energy= 0.498386
Thermal correction to Enthalpy= 0.499330
Thermal correction to Gibbs Free Energy= 0.406083
Sum of electronic and zero-point Energies= -1494.142301
Sum of electronic and thermal Energies= -1494.112797
Sum of electronic and thermal Enthalpies= -1494.111853
Sum of electronic and thermal Free Energies= -1494.205100

| | | | |
|---|---|---|---|
| 6 | -1.715008000 | 0.101268000 | 0.198902000 |
| 6 | -0.501874000 | 0.158611000 | 0.454635000 |
| 6 | -3.019750000 | 0.768932000 | 0.016247000 |
| 1 | -3.398829000 | 0.592639000 | -0.993821000 |
| 1 | -3.752517000 | 0.359408000 | 0.718211000 |
| 6 | 0.699010000 | 1.015811000 | 0.607804000 |
| 1 | 1.379240000 | 0.823866000 | -0.228939000 |
| 1 | 1.243599000 | 0.770351000 | 1.524296000 |
| 6 | 0.301604000 | 2.500923000 | 0.650327000 |
| 1 | -0.310716000 | 2.671917000 | 1.538071000 |
| 1 | 1.218999000 | 3.084547000 | 0.778655000 |
| 6 | -2.848899000 | 2.283189000 | 0.234365000 |
| 1 | -2.498977000 | 2.454169000 | 1.253642000 |
| 1 | -3.832532000 | 2.754316000 | 0.143719000 |
| 6 | -1.906573000 | 2.877038000 | -0.792685000 |
| 6 | -0.408499000 | 2.960705000 | -0.604361000 |
| 6 | -1.242822000 | 4.221970000 | -0.627604000 |
| 1 | -1.165952000 | 4.821040000 | -1.529012000 |
| 1 | 0.168702000 | 2.785149000 | -1.507526000 |
| 1 | -2.215402000 | 2.658331000 | -1.810872000 |
| 6 | -1.438778000 | 5.046067000 | 0.611888000 |
| 1 | -1.615621000 | 4.423646000 | 1.491531000 |
| 1 | -2.303221000 | 5.706004000 | 0.485412000 |
| 8 | -0.258650000 | 5.836019000 | 0.810269000 |

| | | | |
|---|---|---|---|
| 1 | -0.416182000 | 6.424451000 | 1.556582000 |
| 6 | -0.008904000 | -2.444555000 | -0.438055000 |
| 6 | 0.106769000 | -1.906594000 | 0.839997000 |
| 6 | -2.171457000 | -2.136661000 | 1.335433000 |
| 6 | -2.325813000 | -2.143107000 | -0.115492000 |
| 6 | -1.289678000 | -2.605274000 | -0.924000000 |
| 8 | -0.888178000 | -2.153437000 | 1.761132000 |
| 8 | -3.070268000 | -2.163405000 | 2.149414000 |
| 8 | -3.649444000 | -2.152085000 | -0.473662000 |
| 6 | -4.045477000 | -1.768760000 | -1.731404000 |
| 8 | -3.268150000 | -1.404394000 | -2.569748000 |
| 6 | -5.527148000 | -1.848270000 | -1.849674000 |
| 1 | -5.969516000 | -1.133610000 | -1.151717000 |
| 1 | -5.823469000 | -1.612803000 | -2.868193000 |
| 1 | -1.480955000 | -2.943554000 | -1.931580000 |
| 1 | 0.869950000 | -2.591981000 | -1.049598000 |
| 6 | 1.416983000 | -1.720377000 | 1.529873000 |
| 8 | 1.542888000 | -1.703101000 | 2.731003000 |
| 8 | 2.394576000 | -1.534673000 | 0.658664000 |
| 6 | 3.693551000 | -1.192741000 | 1.214212000 |
| 1 | 3.544853000 | -0.414234000 | 1.963460000 |
| 1 | 4.105555000 | -2.085250000 | 1.686069000 |
| 6 | 4.544142000 | -0.707149000 | 0.079779000 |
| 6 | 5.372817000 | -1.588539000 | -0.613172000 |
| 6 | 4.486524000 | 0.633071000 | -0.306305000 |
| 6 | 6.136759000 | -1.135562000 | -1.685420000 |
| 6 | 5.247994000 | 1.086659000 | -1.378582000 |
| 6 | 6.073425000 | 0.201501000 | -2.069160000 |
| 1 | 5.420500000 | -2.628669000 | -0.308038000 |
| 1 | 3.846977000 | 1.319307000 | 0.240496000 |
| 1 | 6.781988000 | -1.824082000 | -2.218370000 |
| 1 | 5.202254000 | 2.128847000 | -1.672118000 |
| 1 | 6.669884000 | 0.555089000 | -2.902176000 |
| 1 | -5.864211000 | -2.847228000 | -1.570058000 |

**TS-PyrS1-BCN**

Frequency -325.9095
Zero-point correction= 0.428761 (Hartree/Particle)
Thermal correction to Energy= 0.459120
Thermal correction to Enthalpy= 0.460064
Thermal correction to Gibbs Free Energy= 0.366117
Sum of electronic and zero-point Energies= -1813.966491
Sum of electronic and thermal Energies= -1813.936132
Sum of electronic and thermal Enthalpies= -1813.935188
Sum of electronic and thermal Free Energies= -1814.029135

| | | | |
|---|---|---|---|
| 6 | -0.931190000 | -1.111266000 | 0.341648000 |
| 6 | -0.877144000 | 0.099587000 | 0.575227000 |
| 6 | -1.690052000 | -2.297956000 | -0.107647000 |
| 1 | -1.233196000 | -2.702262000 | -1.016385000 |
| 1 | -1.652727000 | -3.083028000 | 0.652303000 |
| 6 | -1.504922000 | 1.435616000 | 0.569463000 |
| 1 | -0.981325000 | 2.085175000 | -0.139152000 |
| 1 | -1.437122000 | 1.903968000 | 1.555695000 |
| 6 | -2.985675000 | 1.284428000 | 0.167147000 |
| 1 | -3.496309000 | 0.693649000 | 0.930462000 |
| 1 | -3.437175000 | 2.281237000 | 0.174017000 |
| 6 | -3.145505000 | -1.873884000 | -0.373700000 |

| | | | |
|---|---|---|---|
| 1 | -3.578836000 | -1.521856000 | 0.564235000 |
| 1 | -3.709357000 | -2.760584000 | -0.679482000 |
| 6 | -3.233922000 | -0.820421000 | -1.460485000 |
| 6 | -3.147817000 | 0.669577000 | -1.207423000 |
| 6 | -4.449612000 | 0.051890000 | -1.663814000 |
| 1 | -4.724546000 | 0.241814000 | -2.696162000 |
| 1 | -2.618991000 | 1.221504000 | -1.979368000 |
| 1 | -2.759739000 | -1.140088000 | -2.384002000 |
| 6 | -5.634735000 | -0.031695000 | -0.745896000 |
| 1 | -5.345710000 | -0.313830000 | 0.268512000 |
| 1 | -6.339089000 | -0.782990000 | -1.117917000 |
| 8 | -6.271602000 | 1.252596000 | -0.724633000 |
| 1 | -7.067619000 | 1.181793000 | -0.186339000 |
| 6 | 1.961286000 | 0.113039000 | 0.227812000 |
| 6 | 1.300130000 | 0.426372000 | 1.403074000 |
| 6 | 0.971425000 | -1.809619000 | 2.024012000 |
| 6 | 1.102036000 | -2.074614000 | 0.611272000 |
| 6 | 1.874810000 | -1.223047000 | -0.168779000 |
| 8 | 1.148031000 | -0.521708000 | 2.375533000 |
| 1 | 0.905318000 | -3.086418000 | 0.281377000 |
| 16 | 0.658907000 | -2.929755000 | 3.190313000 |
| 6 | 1.139374000 | 1.797380000 | 1.978276000 |
| 8 | 0.861385000 | 1.998150000 | 3.134285000 |
| 8 | 1.295462000 | 2.724439000 | 1.055193000 |
| 6 | 1.089010000 | 4.087665000 | 1.474788000 |
| 1 | 0.061972000 | 4.209196000 | 1.818426000 |
| 1 | 1.790436000 | 4.337675000 | 2.269357000 |
| 1 | 1.274865000 | 4.690922000 | 0.591304000 |
| 6 | 2.520540000 | 1.177028000 | -0.676742000 |
| 8 | 3.595267000 | 1.696003000 | -0.511933000 |
| 8 | 1.692442000 | 1.436388000 | -1.674331000 |
| 6 | 2.169646000 | 2.383183000 | -2.650232000 |
| 1 | 2.358020000 | 3.343113000 | -2.170125000 |
| 1 | 3.080855000 | 2.001436000 | -3.111334000 |
| 1 | 1.373717000 | 2.469649000 | -3.383930000 |
| 6 | 2.327372000 | -1.702446000 | -1.505558000 |
| 8 | 1.836795000 | -2.650575000 | -2.072660000 |
| 8 | 3.314568000 | -0.965425000 | -1.989765000 |
| 6 | 3.749657000 | -1.272452000 | -3.327340000 |
| 1 | 4.120994000 | -2.295560000 | -3.368525000 |
| 1 | 2.919430000 | -1.141094000 | -4.020690000 |
| 1 | 4.546558000 | -0.566490000 | -3.541393000 |

**TS-PyrS2-BCN**

Frequency -337.9739
Zero-point correction= 0.385953 (Hartree/Particle)
Thermal correction to Energy= 0.411661
Thermal correction to Enthalpy= 0.412606
Thermal correction to Gibbs Free Energy= 0.329318
Sum of electronic and zero-point Energies= -1586.153337
Sum of electronic and thermal Energies= -1586.127628
Sum of electronic and thermal Enthalpies= -1586.126684
Sum of electronic and thermal Free Energies= -1586.209971

| | | | |
|---|---|---|---|
| 6 | -0.374526000 | -0.484935000 | 0.768240000 |
| 6 | -0.505519000 | 0.429286000 | -0.052075000 |
| 6 | -0.967634000 | -1.556744000 | 1.595593000 |
| 1 | -0.558773000 | -2.524809000 | 1.289243000 |

| | | | |
|---|---|---|---|
| 1 | -0.706125000 | -1.412556000 | 2.647453000 |
| 6 | -1.350347000 | 1.201742000 | -0.983570000 |
| 1 | -1.016276000 | 1.023053000 | -2.010982000 |
| 1 | -1.263299000 | 2.275796000 | -0.795266000 |
| 6 | -2.818623000 | 0.766473000 | -0.808522000 |
| 1 | -3.141437000 | 1.032075000 | 0.200342000 |
| 1 | -3.426429000 | 1.351581000 | -1.505464000 |
| 6 | -2.495266000 | -1.532258000 | 1.410505000 |
| 1 | -2.871067000 | -0.570534000 | 1.764542000 |
| 1 | -2.930957000 | -2.306499000 | 2.049665000 |
| 6 | -2.888140000 | -1.789821000 | -0.030957000 |
| 6 | -3.025311000 | -0.708703000 | -1.081404000 |
| 6 | -4.248623000 | -1.448555000 | -0.589655000 |
| 1 | -4.669698000 | -2.173253000 | -1.278735000 |
| 1 | -2.689634000 | -1.005015000 | -2.071045000 |
| 1 | -2.478182000 | -2.722131000 | -0.408809000 |
| 6 | -5.288378000 | -0.756331000 | 0.243659000 |
| 1 | -4.840887000 | -0.114164000 | 1.005213000 |
| 1 | -5.911786000 | -1.499278000 | 0.751656000 |
| 8 | -6.102742000 | 0.033796000 | -0.633531000 |
| 1 | -6.817473000 | 0.418077000 | -0.114203000 |
| 6 | 2.205368000 | 0.211096000 | -0.877872000 |
| 6 | 1.632490000 | 1.314024000 | -0.276023000 |
| 6 | 1.788571000 | 0.443671000 | 1.889366000 |
| 6 | 1.821382000 | -0.834123000 | 1.221754000 |
| 6 | 2.336246000 | -0.910412000 | -0.071340000 |
| 8 | 1.760918000 | 1.513826000 | 1.073097000 |
| 1 | 1.779628000 | -1.714590000 | 1.849518000 |
| 16 | 1.807808000 | 0.680406000 | 3.524451000 |
| 6 | 1.329802000 | 2.555163000 | -1.044439000 |
| 8 | 1.328922000 | 2.576181000 | -2.252868000 |
| 8 | 1.032745000 | 3.579005000 | -0.266505000 |
| 6 | 0.654750000 | 4.792956000 | -0.944974000 |
| 1 | 1.480014000 | 5.141049000 | -1.564691000 |
| 1 | -0.228838000 | 4.610235000 | -1.555959000 |
| 1 | 0.437545000 | 5.507369000 | -0.156861000 |
| 6 | 2.758595000 | -2.234475000 | -0.599027000 |
| 8 | 2.633296000 | -3.270921000 | 0.014837000 |
| 8 | 3.282688000 | -2.151538000 | -1.814501000 |
| 6 | 3.706751000 | -3.392481000 | -2.406760000 |
| 1 | 4.480215000 | -3.850621000 | -1.791376000 |
| 1 | 2.855402000 | -4.064940000 | -2.505846000 |
| 1 | 4.101276000 | -3.127523000 | -3.383217000 |
| 1 | 2.375706000 | 0.203944000 | -1.945595000 |

**TS-PyrS3-BCN**

Frequency -363.9455
Zero-point correction= 0.442021 (Hartree/Particle)
Thermal correction to Energy= 0.470616
Thermal correction to Enthalpy= 0.471560
Thermal correction to Gibbs Free Energy= 0.381698
Sum of electronic and zero-point Energies= -1664.715278
Sum of electronic and thermal Energies= -1664.686683
Sum of electronic and thermal Enthalpies= -1664.685738
Sum of electronic and thermal Free Energies= -1664.775600

| | | | |
|---|---|---|---|
| 6 | 0.508928000 | -1.263207000 | 0.105577000 |
| 6 | 0.606951000 | -0.558307000 | -0.906447000 |

| | | | |
|---|---|---|---|
| 6 | 1.087604000 | -1.918279000 | 1.294973000 |
| 1 | 0.607816000 | -1.528351000 | 2.198354000 |
| 1 | 0.899320000 | -2.994971000 | 1.266690000 |
| 6 | 1.468801000 | 0.296153000 | -1.753122000 |
| 1 | 1.079223000 | 1.319974000 | -1.748295000 |
| 1 | 1.453905000 | -0.047231000 | -2.789783000 |
| 6 | 2.911934000 | 0.271578000 | -1.218098000 |
| 1 | 3.304572000 | -0.742648000 | -1.316419000 |
| 1 | 3.520206000 | 0.914380000 | -1.862379000 |
| 6 | 2.599894000 | -1.632189000 | 1.329688000 |
| 1 | 3.060008000 | -2.076974000 | 0.445589000 |
| 1 | 3.024633000 | -2.135052000 | 2.204071000 |
| 6 | 2.878973000 | -0.144632000 | 1.416973000 |
| 6 | 3.009552000 | 0.760693000 | 0.211644000 |
| 6 | 4.215662000 | 0.467625000 | 1.074982000 |
| 1 | 4.541727000 | 1.276015000 | 1.721269000 |
| 1 | 2.585702000 | 1.752004000 | 0.349163000 |
| 1 | 2.386036000 | 0.318235000 | 2.267265000 |
| 6 | 5.352024000 | -0.368183000 | 0.560427000 |
| 1 | 4.998932000 | -1.231062000 | -0.008222000 |
| 1 | 5.950914000 | -0.737684000 | 1.399282000 |
| 8 | 6.166195000 | 0.461200000 | -0.280114000 |
| 1 | 6.934084000 | -0.052513000 | -0.553450000 |
| 6 | -1.988010000 | 0.294232000 | -0.789903000 |
| 6 | -1.445737000 | -0.456358000 | -1.832274000 |
| 6 | -1.683707000 | -2.484268000 | -0.652287000 |
| 6 | -1.708215000 | -1.663408000 | 0.524164000 |
| 6 | -2.153638000 | -0.344815000 | 0.428876000 |
| 8 | -1.645561000 | -1.817067000 | -1.822491000 |
| 1 | -1.700964000 | -2.177901000 | 1.474835000 |
| 16 | -1.718941000 | -4.141013000 | -0.682071000 |
| 6 | -2.095203000 | 1.778708000 | -0.968790000 |
| 8 | -2.999932000 | 2.327460000 | -1.551639000 |
| 8 | -1.040466000 | 2.393420000 | -0.459421000 |
| 6 | -1.005340000 | 3.839620000 | -0.566719000 |
| 1 | -1.136418000 | 4.107192000 | -1.615898000 |
| 1 | -1.841734000 | 4.236216000 | 0.011908000 |
| 6 | -2.581740000 | 0.432980000 | 1.621286000 |
| 8 | -3.064975000 | 1.541905000 | 1.544458000 |
| 8 | -2.379707000 | -0.211151000 | 2.760530000 |
| 6 | -2.776208000 | 0.482186000 | 3.958234000 |
| 1 | -2.212523000 | 1.409686000 | 4.051903000 |
| 1 | -2.540796000 | -0.193609000 | 4.774965000 |
| 6 | -1.239565000 | 0.057662000 | -3.215748000 |
| 1 | -0.847509000 | 1.074911000 | -3.195958000 |
| 1 | -0.550441000 | -0.588714000 | -3.757877000 |
| 1 | -2.200636000 | 0.062318000 | -3.737658000 |
| 1 | -3.845011000 | 0.690622000 | 3.926952000 |
| 6 | 0.330386000 | 4.287350000 | -0.028958000 |
| 1 | 0.390164000 | 5.375917000 | -0.078395000 |
| 1 | 0.452778000 | 3.980127000 | 1.011130000 |
| 1 | 1.144582000 | 3.868323000 | -0.623361000 |

**TS-PyrS5-BCN**

Frequency -381.7988

Zero-point correction= 0.466078 (Hartree/Particle)

Thermal correction to Energy= 0.496212

Thermal correction to Enthalpy= 0.497156

Thermal correction to Gibbs Free Energy= 0.401741
Sum of electronic and zero-point Energies= -1817.089696
Sum of electronic and thermal Energies= -1817.059561
Sum of electronic and thermal Enthalpies= -1817.058617
Sum of electronic and thermal Free Energies= -1817.154032

| | | | |
|---|---|---|---|
| 6 | 1.594995000 | 0.288764000 | -0.119259000 |
| 6 | 0.383570000 | 0.244298000 | -0.382035000 |
| 6 | 2.860161000 | 1.026032000 | 0.048152000 |
| 1 | 3.239256000 | 0.910377000 | 1.066553000 |
| 1 | 3.618224000 | 0.632050000 | -0.635295000 |
| 6 | -0.866155000 | 1.025582000 | -0.564813000 |
| 1 | -1.533262000 | 0.825878000 | 0.280695000 |
| 1 | -1.396894000 | 0.715259000 | -1.469644000 |
| 6 | -0.553279000 | 2.527703000 | -0.664306000 |
| 1 | 0.052483000 | 2.698296000 | -1.556795000 |
| 1 | -1.501883000 | 3.052163000 | -0.817993000 |
| 6 | 2.604580000 | 2.519973000 | -0.227983000 |
| 1 | 2.252717000 | 2.633459000 | -1.254644000 |
| 1 | 3.559295000 | 3.048964000 | -0.148991000 |
| 6 | 1.621242000 | 3.094803000 | 0.771239000 |
| 6 | 0.122072000 | 3.077638000 | 0.573401000 |
| 6 | 0.876540000 | 4.388431000 | 0.550856000 |
| 1 | 0.755768000 | 5.015602000 | 1.427961000 |
| 1 | -0.448550000 | 2.902533000 | 1.480825000 |
| 1 | 1.936286000 | 2.934729000 | 1.798377000 |
| 6 | 1.028924000 | 5.174021000 | -0.719480000 |
| 1 | 1.268761000 | 4.531887000 | -1.569565000 |
| 1 | 1.837546000 | 5.903853000 | -0.608887000 |
| 8 | -0.205500000 | 5.860254000 | -0.967336000 |
| 1 | -0.087344000 | 6.417105000 | -1.744579000 |
| 6 | 0.006923000 | -2.315760000 | 0.657584000 |
| 6 | -0.081820000 | -1.846705000 | -0.654257000 |
| 6 | 2.218380000 | -2.018401000 | -1.059647000 |
| 6 | 2.326105000 | -1.946388000 | 0.378712000 |
| 6 | 1.273905000 | -2.396210000 | 1.183601000 |
| 8 | 0.955209000 | -2.109052000 | -1.513861000 |
| 16 | 3.469482000 | -2.085599000 | -2.143096000 |
| 8 | 3.613712000 | -1.857983000 | 0.833270000 |
| 6 | 3.890453000 | -1.350285000 | 2.079923000 |
| 8 | 3.039400000 | -0.918233000 | 2.807074000 |
| 6 | 5.356216000 | -1.394350000 | 2.335200000 |
| 1 | 5.718271000 | -2.415230000 | 2.205846000 |
| 1 | 5.857297000 | -0.761626000 | 1.599051000 |
| 1 | 1.451380000 | -2.684649000 | 2.209442000 |
| 1 | -0.885935000 | -2.484523000 | 1.242791000 |
| 6 | -1.371010000 | -1.776764000 | -1.408256000 |
| 8 | -1.441946000 | -1.823674000 | -2.612252000 |
| 8 | -2.394602000 | -1.616772000 | -0.587938000 |
| 6 | -3.688006000 | -1.389417000 | -1.212147000 |
| 1 | -3.557469000 | -0.635809000 | -1.989862000 |
| 1 | -4.018104000 | -2.327768000 | -1.658573000 |
| 6 | -4.618032000 | -0.915013000 | -0.137000000 |
| 6 | -5.447007000 | -1.814166000 | 0.531857000 |
| 6 | -4.635432000 | 0.434919000 | 0.218475000 |
| 6 | -6.286921000 | -1.369150000 | 1.549451000 |
| 6 | -5.471830000 | 0.880324000 | 1.236661000 |
| 6 | -6.298113000 | -0.022613000 | 1.903190000 |

| | | | |
|---|---|---|---|
| 1 | -5.435323000 | -2.862045000 | 0.250807000 |
| 1 | -3.994748000 | 1.134672000 | -0.309672000 |
| 1 | -6.932328000 | -2.071737000 | 2.063482000 |
| 1 | -5.484067000 | 1.929750000 | 1.506764000 |
| 1 | -6.952944000 | 0.324663000 | 2.693948000 |
| 1 | 5.560895000 | -1.038184000 | 3.341117000 |

**C1**

Zero-point correction= 0.436699 (Hartree/Particle)
Thermal correction to Energy= 0.465503
Thermal correction to Enthalpy= 0.466447
Thermal correction to Gibbs Free Energy= 0.376717
Sum of electronic and zero-point Energies= -1491.090681
Sum of electronic and thermal Energies= -1491.061877
Sum of electronic and thermal Enthalpies= -1491.060933
Sum of electronic and thermal Free Energies= -1491.150662

| | | | |
|---|---|---|---|
| 6 | -0.812877000 | -1.139069000 | 0.694653000 |
| 6 | -0.763959000 | 0.198877000 | 0.642235000 |
| 6 | -1.832317000 | -2.236463000 | 0.503763000 |
| 1 | -1.411915000 | -2.892529000 | -0.270273000 |
| 1 | -1.802904000 | -2.831976000 | 1.423555000 |
| 6 | -1.713293000 | 1.325491000 | 0.307622000 |
| 1 | -1.278787000 | 1.857598000 | -0.546007000 |
| 1 | -1.675792000 | 2.032639000 | 1.145419000 |
| 6 | -3.179691000 | 1.056690000 | -0.009712000 |
| 1 | -3.672906000 | 0.602399000 | 0.849349000 |
| 1 | -3.649339000 | 2.035462000 | -0.148213000 |
| 6 | -3.285012000 | -1.941848000 | 0.159759000 |
| 1 | -3.747833000 | -1.371716000 | 0.964273000 |
| 1 | -3.803562000 | -2.905339000 | 0.131461000 |
| 6 | -3.424839000 | -1.260658000 | -1.180254000 |
| 6 | -3.359810000 | 0.239819000 | -1.266237000 |
| 6 | -4.654740000 | -0.477226000 | -1.567575000 |
| 1 | -4.919531000 | -0.526800000 | -2.618646000 |
| 1 | -2.845965000 | 0.649429000 | -2.130432000 |
| 1 | -2.954648000 | -1.806666000 | -1.992228000 |
| 6 | -5.849054000 | -0.363012000 | -0.665302000 |
| 1 | -5.572842000 | -0.422079000 | 0.389454000 |
| 1 | -6.552975000 | -1.174688000 | -0.876039000 |
| 8 | -6.479900000 | 0.898186000 | -0.927126000 |
| 1 | -7.284752000 | 0.945649000 | -0.399943000 |
| 6 | 1.689680000 | 0.068748000 | 0.125749000 |
| 6 | 0.659398000 | 0.691899000 | 1.065370000 |
| 6 | 0.839217000 | -1.170062000 | 2.447363000 |
| 6 | 0.548270000 | -1.769538000 | 1.088328000 |
| 6 | 1.640668000 | -1.259383000 | 0.182079000 |
| 8 | 0.923189000 | 0.173100000 | 2.391948000 |
| 8 | 0.977588000 | -1.756125000 | 3.488337000 |
| 1 | 0.512317000 | -2.852132000 | 1.150210000 |
| 6 | 2.688701000 | 0.907419000 | -0.606458000 |
| 8 | 3.589894000 | 1.473243000 | -0.034116000 |
| 8 | 2.447894000 | 0.976943000 | -1.901406000 |
| 6 | 3.359629000 | 1.795761000 | -2.660293000 |
| 1 | 3.330567000 | 2.818689000 | -2.284930000 |
| 1 | 4.367916000 | 1.389675000 | -2.582925000 |
| 1 | 3.004235000 | 1.750919000 | -3.685249000 |
| 6 | 2.547998000 | -2.209883000 | -0.508745000 |
| 8 | 2.362631000 | -3.404725000 | -0.532776000 |

| | | | |
|---|---|---|---|
| 8 | 3.570901000 | -1.598364000 | -1.086992000 |
| 6 | 4.483893000 | -2.434617000 | -1.822059000 |
| 1 | 4.943555000 | -3.157653000 | -1.149353000 |
| 1 | 3.951363000 | -2.945787000 | -2.623112000 |
| 1 | 5.230913000 | -1.758536000 | -2.227463000 |
| 6 | 0.807545000 | 2.206063000 | 1.192427000 |
| 8 | 0.832118000 | 2.796629000 | 2.239718000 |
| 8 | 0.917643000 | 2.758437000 | -0.000183000 |
| 6 | 1.133954000 | 4.183139000 | -0.032307000 |
| 1 | 0.296558000 | 4.691005000 | 0.444534000 |
| 1 | 2.066628000 | 4.418236000 | 0.479750000 |
| 1 | 1.193868000 | 4.442737000 | -1.084860000 |

**C2**

Zero-point correction= 0.393367 (Hartree/Particle)
Thermal correction to Energy= 0.417688
Thermal correction to Enthalpy= 0.418633
Thermal correction to Gibbs Free Energy= 0.338844
Sum of electronic and zero-point Energies= -1263.276161
Sum of electronic and thermal Energies= -1263.251839
Sum of electronic and thermal Enthalpies= -1263.250895
Sum of electronic and thermal Free Energies= -1263.330684

| | | | |
|---|---|---|---|
| 6 | -0.110143000 | -0.791347000 | 0.775031000 |
| 6 | -0.272933000 | 0.477959000 | 0.377804000 |
| 6 | -0.969394000 | -2.007489000 | 1.024881000 |
| 1 | -0.534530000 | -2.805606000 | 0.408418000 |
| 1 | -0.765988000 | -2.308193000 | 2.059335000 |
| 6 | -1.407558000 | 1.351060000 | -0.104107000 |
| 1 | -1.128178000 | 1.694989000 | -1.107607000 |
| 1 | -1.409169000 | 2.250070000 | 0.523531000 |
| 6 | -2.838089000 | 0.829241000 | -0.171731000 |
| 1 | -3.191487000 | 0.577308000 | 0.827892000 |
| 1 | -3.456670000 | 1.665990000 | -0.510970000 |
| 6 | -2.475327000 | -1.989068000 | 0.804623000 |
| 1 | -2.940600000 | -1.269053000 | 1.476801000 |
| 1 | -2.851155000 | -2.972906000 | 1.102660000 |
| 6 | -2.828152000 | -1.734996000 | -0.641561000 |
| 6 | -2.992758000 | -0.323013000 | -1.134269000 |
| 6 | -4.187255000 | -1.246108000 | -1.077821000 |
| 1 | -4.523781000 | -1.622243000 | -2.038387000 |
| 1 | -2.614997000 | -0.114429000 | -2.130502000 |
| 1 | -2.350139000 | -2.427065000 | -1.328008000 |
| 6 | -5.311241000 | -1.025179000 | -0.107729000 |
| 1 | -4.948239000 | -0.750303000 | 0.884610000 |
| 1 | -5.902512000 | -1.941428000 | -0.009440000 |
| 8 | -6.139347000 | 0.025989000 | -0.624214000 |
| 1 | -6.896900000 | 0.125860000 | -0.037461000 |
| 6 | 2.088205000 | 0.513302000 | -0.450191000 |
| 6 | 1.098618000 | 1.230580000 | 0.444017000 |
| 6 | 1.717031000 | -0.155520000 | 2.208584000 |
| 6 | 1.369419000 | -1.127510000 | 1.104611000 |
| 6 | 2.251435000 | -0.747094000 | -0.061249000 |
| 8 | 1.582400000 | 1.121982000 | 1.810757000 |
| 1 | 1.505745000 | -2.149803000 | 1.441874000 |
| 8 | 2.065961000 | -0.419800000 | 3.330943000 |
| 6 | 1.003891000 | 2.726200000 | 0.177663000 |
| 8 | 0.917542000 | 3.559835000 | 1.043244000 |

| | | | |
|---|---|---|---|
| 8 | 1.012497000 | 2.977677000 | -1.118096000 |
| 6 | 0.815562000 | 4.354047000 | -1.498000000 |
| 1 | -0.150273000 | 4.698042000 | -1.128460000 |
| 1 | 1.619116000 | 4.966715000 | -1.091966000 |
| 1 | 0.837473000 | 4.360390000 | -2.583486000 |
| 6 | 3.181772000 | -1.730205000 | -0.656671000 |
| 8 | 3.320923000 | -2.858713000 | -0.237711000 |
| 8 | 3.841448000 | -1.243589000 | -1.702234000 |
| 6 | 4.759689000 | -2.145535000 | -2.344432000 |
| 1 | 5.531003000 | -2.454422000 | -1.639757000 |
| 1 | 4.222985000 | -3.015479000 | -2.721571000 |
| 1 | 5.195021000 | -1.580816000 | -3.163658000 |
| 1 | 2.585255000 | 1.009387000 | -1.272066000 |

**C3**

Zero-point correction= 0.450450 (Hartree/Particle)
Thermal correction to Energy= 0.477319
Thermal correction to Enthalpy= 0.478263
Thermal correction to Gibbs Free Energy= 0.392698
Sum of electronic and zero-point Energies= -1341.837489
Sum of electronic and thermal Energies= -1341.810620
Sum of electronic and thermal Enthalpies= -1341.809676
Sum of electronic and thermal Free Energies= -1341.895240

| | | | |
|---|---|---|---|
| 6 | 0.844945000 | 1.259188000 | -0.295239000 |
| 6 | 0.836193000 | 0.634050000 | 0.891056000 |
| 6 | 1.805471000 | 1.520942000 | -1.431201000 |
| 1 | 1.286257000 | 1.186590000 | -2.339232000 |
| 1 | 1.865809000 | 2.611739000 | -1.527596000 |
| 6 | 1.791823000 | -0.241781000 | 1.664823000 |
| 1 | 1.262402000 | -1.187916000 | 1.837150000 |
| 1 | 1.911918000 | 0.204088000 | 2.656384000 |
| 6 | 3.183178000 | -0.575785000 | 1.137490000 |
| 1 | 3.785366000 | 0.330365000 | 1.073931000 |
| 1 | 3.659058000 | -1.201182000 | 1.899557000 |
| 6 | 3.214192000 | 0.943980000 | -1.445358000 |
| 1 | 3.787827000 | 1.337103000 | -0.607043000 |
| 1 | 3.701680000 | 1.316192000 | -2.351933000 |
| 6 | 3.195014000 | -0.565950000 | -1.467823000 |
| 6 | 3.158166000 | -1.327176000 | -0.171162000 |
| 6 | 4.379248000 | -1.406635000 | -1.057973000 |
| 1 | 4.491715000 | -2.338112000 | -1.602986000 |
| 1 | 2.539348000 | -2.219189000 | -0.153725000 |
| 1 | 2.602621000 | -0.975965000 | -2.280055000 |
| 6 | 5.690583000 | -0.799902000 | -0.650965000 |
| 1 | 5.565433000 | 0.198592000 | -0.227065000 |
| 1 | 6.346635000 | -0.716063000 | -1.523451000 |
| 8 | 6.296305000 | -1.662758000 | 0.321612000 |
| 1 | 7.171240000 | -1.314577000 | 0.524966000 |
| 6 | -1.610648000 | 0.295299000 | 0.695277000 |
| 6 | -0.529220000 | 0.840024000 | 1.623892000 |
| 6 | -0.717619000 | 2.909801000 | 0.518226000 |
| 6 | -0.516475000 | 1.932953000 | -0.616469000 |
| 6 | -1.622236000 | 0.915621000 | -0.482936000 |
| 8 | -0.753147000 | 2.293334000 | 1.702983000 |
| 1 | -0.520088000 | 2.451392000 | -1.569810000 |
| 8 | -0.824843000 | 4.109040000 | 0.428673000 |
| 6 | -2.548867000 | -0.789528000 | 1.112456000 |

| | | | |
|---|---|---|---|
| 8 | -3.396770000 | -0.639666000 | 1.960939000 |
| 8 | -2.334318000 | -1.913271000 | 0.449224000 |
| 6 | -3.314887000 | -2.967762000 | 0.631366000 |
| 1 | -3.209350000 | -3.360372000 | 1.643568000 |
| 1 | -4.306144000 | -2.524749000 | 0.521067000 |
| 6 | -2.546850000 | 0.650233000 | -1.610989000 |
| 8 | -2.328135000 | 1.002073000 | -2.748374000 |
| 8 | -3.623036000 | -0.026454000 | -1.232817000 |
| 6 | -4.510440000 | -0.461215000 | -2.278265000 |
| 1 | -4.928176000 | 0.403444000 | -2.792924000 |
| 1 | -5.294141000 | -1.022887000 | -1.778086000 |
| 6 | -0.644660000 | 0.359474000 | 3.052599000 |
| 1 | -0.545017000 | -0.725536000 | 3.102783000 |
| 1 | 0.124714000 | 0.819538000 | 3.670862000 |
| 1 | -1.623726000 | 0.638265000 | 3.442469000 |
| 1 | -3.969646000 | -1.094758000 | -2.980722000 |
| 6 | -3.043728000 | -4.008673000 | -0.424792000 |
| 1 | -3.763438000 | -4.822182000 | -0.319346000 |
| 1 | -3.151645000 | -3.576019000 | -1.421514000 |
| 1 | -2.038190000 | -4.419125000 | -0.319254000 |

**C4**

Zero-point correction= 0.473485 (Hartree/Particle)
Thermal correction to Energy= 0.502148
Thermal correction to Enthalpy= 0.503092
Thermal correction to Gibbs Free Energy= 0.411562
Sum of electronic and zero-point Energies= -1494.207038
Sum of electronic and thermal Energies= -1494.178376
Sum of electronic and thermal Enthalpies= -1494.177431
Sum of electronic and thermal Free Energies= -1494.268961

| | | | |
|---|---|---|---|
| 6 | 1.821657000 | -0.225678000 | -0.132870000 |
| 6 | 0.559472000 | -0.115419000 | -0.574330000 |
| 6 | 2.986350000 | 0.711911000 | 0.085968000 |
| 1 | 3.331896000 | 0.578030000 | 1.115295000 |
| 1 | 3.796551000 | 0.336285000 | -0.548748000 |
| 6 | -0.420767000 | 0.997021000 | -0.882150000 |
| 1 | -1.253641000 | 0.852663000 | -0.182883000 |
| 1 | -0.836056000 | 0.789271000 | -1.875375000 |
| 6 | -0.048757000 | 2.475504000 | -0.855822000 |
| 1 | 0.671574000 | 2.696382000 | -1.642625000 |
| 1 | -0.960862000 | 3.018923000 | -1.122462000 |
| 6 | 2.840036000 | 2.209251000 | -0.149290000 |
| 1 | 2.553977000 | 2.400238000 | -1.182585000 |
| 1 | 3.835921000 | 2.644413000 | -0.020015000 |
| 6 | 1.886084000 | 2.834787000 | 0.840699000 |
| 6 | 0.427603000 | 2.948020000 | 0.494537000 |
| 6 | 1.267971000 | 4.195956000 | 0.635058000 |
| 1 | 1.097459000 | 4.763708000 | 1.543722000 |
| 1 | -0.276753000 | 2.766562000 | 1.300366000 |
| 1 | 2.116655000 | 2.589337000 | 1.872791000 |
| 6 | 1.607819000 | 5.057634000 | -0.546215000 |
| 1 | 1.927769000 | 4.465124000 | -1.405858000 |
| 1 | 2.424351000 | 5.738117000 | -0.284128000 |
| 8 | 0.442667000 | 5.818447000 | -0.893197000 |
| 1 | 0.681944000 | 6.426404000 | -1.601289000 |
| 6 | -0.019924000 | -2.344798000 | 0.439598000 |
| 6 | -0.044703000 | -1.531181000 | -0.834869000 |

| | | | |
|---|---|---|---|
| 6 | 2.093650000 | -2.303144000 | -1.334385000 |
| 6 | 2.232095000 | -1.724970000 | 0.075211000 |
| 6 | 1.211305000 | -2.480473000 | 0.902148000 |
| 8 | 0.844991000 | -2.186831000 | -1.790492000 |
| 8 | 2.971578000 | -2.800498000 | -1.987179000 |
| 8 | 3.571026000 | -1.937795000 | 0.465540000 |
| 6 | 3.922971000 | -1.738221000 | 1.759771000 |
| 8 | 3.141770000 | -1.349439000 | 2.591530000 |
| 6 | 5.368048000 | -2.037146000 | 1.970304000 |
| 1 | 5.957473000 | -1.318472000 | 1.396108000 |
| 1 | 5.607219000 | -1.956314000 | 3.027274000 |
| 1 | 1.524203000 | -3.086633000 | 1.740006000 |
| 1 | -0.917919000 | -2.782271000 | 0.851293000 |
| 6 | -1.404895000 | -1.526152000 | -1.518639000 |
| 8 | -1.569435000 | -1.690049000 | -2.700808000 |
| 8 | -2.366484000 | -1.305619000 | -0.640721000 |
| 6 | -3.701617000 | -1.109551000 | -1.181986000 |
| 1 | -3.631489000 | -0.360667000 | -1.972761000 |
| 1 | -4.044775000 | -2.056070000 | -1.599568000 |
| 6 | -4.569700000 | -0.643765000 | -0.052503000 |
| 6 | -5.459694000 | -1.518514000 | 0.567114000 |
| 6 | -4.469878000 | 0.673898000 | 0.399244000 |
| 6 | -6.246508000 | -1.081206000 | 1.630412000 |
| 6 | -5.250075000 | 1.109923000 | 1.464091000 |
| 6 | -6.140459000 | 0.231409000 | 2.080282000 |
| 1 | -5.537137000 | -2.541282000 | 0.213681000 |
| 1 | -3.779968000 | 1.355272000 | -0.089479000 |
| 1 | -6.939841000 | -1.764926000 | 2.106025000 |
| 1 | -5.169759000 | 2.133667000 | 1.810368000 |
| 1 | -6.752154000 | 0.572288000 | 2.907516000 |
| 1 | 5.595947000 | -3.036348000 | 1.597678000 |

**C1 (from PyrS1)**

Zero-point correction= 0.434190 (Hartree/Particle)
Thermal correction to Energy= 0.463441
Thermal correction to Enthalpy= 0.464385
Thermal correction to Gibbs Free Energy= 0.373340
Sum of electronic and zero-point Energies= -1814.035931
Sum of electronic and thermal Energies= -1814.006680
Sum of electronic and thermal Enthalpies= -1814.005736
Sum of electronic and thermal Free Energies= -1814.096780

| | | | |
|---|---|---|---|
| 6 | -0.822155000 | -1.184887000 | 0.256936000 |
| 6 | -0.797054000 | 0.120207000 | 0.558133000 |
| 6 | -1.831488000 | -2.221480000 | -0.175240000 |
| 1 | -1.426441000 | -2.656850000 | -1.098330000 |
| 1 | -1.765521000 | -3.024955000 | 0.568239000 |
| 6 | -1.773009000 | 1.273507000 | 0.559725000 |
| 1 | -1.370692000 | 2.023832000 | -0.130253000 |
| 1 | -1.723851000 | 1.732510000 | 1.554790000 |
| 6 | -3.242589000 | 1.062628000 | 0.214498000 |
| 1 | -3.703584000 | 0.388781000 | 0.936120000 |
| 1 | -3.732720000 | 2.032226000 | 0.346125000 |
| 6 | -3.297173000 | -1.878727000 | -0.399256000 |
| 1 | -3.747371000 | -1.545133000 | 0.534889000 |
| 1 | -3.802304000 | -2.812648000 | -0.665035000 |
| 6 | -3.480203000 | -0.876319000 | -1.513623000 |
| 6 | -3.440669000 | 0.596414000 | -1.207552000 |

| | | | |
|---|---|---|---|
| 6 | -4.730698000 | -0.045045000 | -1.661427000 |
| 1 | -5.017898000 | 0.174435000 | -2.684513000 |
| 1 | -2.953517000 | 1.226947000 | -1.944970000 |
| 1 | -3.020495000 | -1.182723000 | -2.448137000 |
| 6 | -5.906580000 | -0.195773000 | -0.740698000 |
| 1 | -5.605435000 | -0.516624000 | 0.258611000 |
| 1 | -6.597872000 | -0.944338000 | -1.141526000 |
| 8 | -6.570026000 | 1.072912000 | -0.658794000 |
| 1 | -7.361189000 | 0.962250000 | -0.120202000 |
| 6 | 1.643760000 | 0.188593000 | -0.028964000 |
| 6 | 0.627259000 | 0.515138000 | 1.057502000 |
| 6 | 0.878805000 | -1.643227000 | 1.898177000 |
| 6 | 0.558801000 | -1.865350000 | 0.435285000 |
| 6 | 1.617980000 | -1.108101000 | -0.325142000 |
| 8 | 0.937373000 | -0.347996000 | 2.192547000 |
| 1 | 0.539110000 | -2.921734000 | 0.192743000 |
| 16 | 1.116579000 | -2.794286000 | 3.029945000 |
| 6 | 0.758154000 | 1.936226000 | 1.598662000 |
| 8 | 0.799486000 | 2.212067000 | 2.768320000 |
| 8 | 0.832941000 | 2.799452000 | 0.604513000 |
| 6 | 1.031917000 | 4.181230000 | 0.964371000 |
| 1 | 0.202437000 | 4.521531000 | 1.583009000 |
| 1 | 1.975864000 | 4.282415000 | 1.499151000 |
| 1 | 1.059438000 | 4.723443000 | 0.024176000 |
| 6 | 2.615174000 | 1.214383000 | -0.522038000 |
| 8 | 3.521776000 | 1.620553000 | 0.165378000 |
| 8 | 2.341025000 | 1.629704000 | -1.743444000 |
| 6 | 3.221342000 | 2.643174000 | -2.268509000 |
| 1 | 3.189715000 | 3.523227000 | -1.626070000 |
| 1 | 4.236390000 | 2.251269000 | -2.328699000 |
| 1 | 2.838799000 | 2.874151000 | -3.258080000 |
| 6 | 2.540160000 | -1.828567000 | -1.238935000 |
| 8 | 2.408086000 | -2.994762000 | -1.530256000 |
| 8 | 3.512893000 | -1.049708000 | -1.688281000 |
| 6 | 4.438081000 | -1.650832000 | -2.613506000 |
| 1 | 4.950827000 | -2.483008000 | -2.132824000 |
| 1 | 3.903039000 | -1.993787000 | -3.498260000 |
| 1 | 5.140673000 | -0.863558000 | -2.870342000 |

**C2 (from PyrS2)**

Zero-point correction= 0.390865 (Hartree/Particle)
Thermal correction to Energy= 0.415622
Thermal correction to Enthalpy= 0.416566
Thermal correction to Gibbs Free Energy= 0.335188
Sum of electronic and zero-point Energies= -1586.221623
Sum of electronic and thermal Energies= -1586.196866
Sum of electronic and thermal Enthalpies= -1586.195922
Sum of electronic and thermal Free Energies= -1586.277300

| | | | |
|---|---|---|---|
| 6 | -0.156885000 | -0.657337000 | 0.671082000 |
| 6 | -0.365620000 | 0.487324000 | 0.007476000 |
| 6 | -0.982527000 | -1.768680000 | 1.271619000 |
| 1 | -0.586803000 | -2.697866000 | 0.841113000 |
| 1 | -0.704281000 | -1.806870000 | 2.331776000 |
| 6 | -1.537755000 | 1.230880000 | -0.589648000 |
| 1 | -1.320540000 | 1.331426000 | -1.660724000 |
| 1 | -1.511492000 | 2.251121000 | -0.188620000 |
| 6 | -2.965650000 | 0.714294000 | -0.450282000 |

| | | | |
|---|---|---|---|
| 1 | -3.264627000 | 0.717632000 | 0.597587000 |
| 1 | -3.607511000 | 1.446059000 | -0.950604000 |
| 6 | -2.499619000 | -1.785718000 | 1.155600000 |
| 1 | -2.919677000 | -0.916332000 | 1.659833000 |
| 1 | -2.851703000 | -2.661436000 | 1.709826000 |
| 6 | -2.948507000 | -1.888169000 | -0.282967000 |
| 6 | -3.164595000 | -0.637446000 | -1.091031000 |
| 6 | -4.339945000 | -1.516458000 | -0.732899000 |
| 1 | -4.733819000 | -2.112983000 | -1.549179000 |
| 1 | -2.855593000 | -0.678621000 | -2.130962000 |
| 1 | -2.506212000 | -2.726927000 | -0.811656000 |
| 6 | -5.397932000 | -1.067150000 | 0.232447000 |
| 1 | -4.970059000 | -0.578110000 | 1.110030000 |
| 1 | -5.975656000 | -1.931280000 | 0.576407000 |
| 8 | -6.265660000 | -0.153448000 | -0.452136000 |
| 1 | -6.983331000 | 0.080456000 | 0.146524000 |
| 6 | 1.925655000 | 0.323133000 | -0.982317000 |
| 6 | 0.995572000 | 1.221923000 | -0.205814000 |
| 6 | 1.765166000 | 0.272147000 | 1.781315000 |
| 6 | 1.346029000 | -0.920924000 | 0.952791000 |
| 6 | 2.134637000 | -0.816458000 | -0.331128000 |
| 8 | 1.588454000 | 1.410487000 | 1.122017000 |
| 1 | 1.513060000 | -1.849163000 | 1.487092000 |
| 16 | 2.352396000 | 0.244391000 | 3.306920000 |
| 6 | 0.867971000 | 2.598056000 | -0.840285000 |
| 8 | 0.857690000 | 2.743955000 | -2.037951000 |
| 8 | 0.744495000 | 3.567974000 | 0.042288000 |
| 6 | 0.552764000 | 4.890371000 | -0.499767000 |
| 1 | 1.407028000 | 5.161708000 | -1.118346000 |
| 1 | -0.365181000 | 4.913843000 | -1.086290000 |
| 1 | 0.477406000 | 5.546725000 | 0.361715000 |
| 6 | 3.060423000 | -1.896511000 | -0.734874000 |
| 8 | 3.285240000 | -2.874587000 | -0.055739000 |
| 8 | 3.614837000 | -1.677375000 | -1.922294000 |
| 6 | 4.528887000 | -2.685482000 | -2.388682000 |
| 1 | 5.363173000 | -2.779931000 | -1.694399000 |
| 1 | 4.011011000 | -3.638876000 | -2.488145000 |
| 1 | 4.875434000 | -2.336451000 | -3.356975000 |
| 1 | 2.362178000 | 0.632790000 | -1.922035000 |

**C3 (from PyrS3)**

Zero-point correction= 0.447507 (Hartree/Particle)
Thermal correction to Energy= 0.475103
Thermal correction to Enthalpy= 0.476047
Thermal correction to Gibbs Free Energy= 0.387802
Sum of electronic and zero-point Energies= -1664.783545
Sum of electronic and thermal Energies= -1664.755950
Sum of electronic and thermal Enthalpies= -1664.755005
Sum of electronic and thermal Free Energies= -1664.843250

| | | | |
|---|---|---|---|
| 6 | 0.655455000 | -1.150856000 | 0.010161000 |
| 6 | 0.730044000 | -0.218551000 | -0.950816000 |
| 6 | 1.581183000 | -1.856387000 | 0.972277000 |
| 1 | 1.127214000 | -1.735394000 | 1.964540000 |
| 1 | 1.487103000 | -2.925995000 | 0.747418000 |
| 6 | 1.789480000 | 0.711553000 | -1.492717000 |
| 1 | 1.389379000 | 1.726622000 | -1.370619000 |
| 1 | 1.839916000 | 0.552225000 | -2.573531000 |

| | | | |
|---|---|---|---|
| 6 | 3.217633000 | 0.706215000 | -0.957694000 |
| 1 | 3.700941000 | -0.242819000 | -1.189356000 |
| 1 | 3.761358000 | 1.466928000 | -1.527037000 |
| 6 | 3.056836000 | -1.497588000 | 1.073083000 |
| 1 | 3.555040000 | -1.701796000 | 0.126352000 |
| 1 | 3.501377000 | -2.177630000 | 1.806536000 |
| 6 | 3.247871000 | -0.070315000 | 1.529039000 |
| 6 | 3.303622000 | 1.036895000 | 0.512160000 |
| 6 | 4.531545000 | 0.695491000 | 1.323996000 |
| 1 | 4.774436000 | 1.401658000 | 2.111243000 |
| 1 | 2.810248000 | 1.965266000 | 0.783550000 |
| 1 | 2.724939000 | 0.157306000 | 2.452938000 |
| 6 | 5.745960000 | 0.075673000 | 0.696178000 |
| 1 | 5.485271000 | -0.715072000 | -0.010040000 |
| 1 | 6.382142000 | -0.362025000 | 1.472427000 |
| 8 | 6.467936000 | 1.109904000 | 0.013155000 |
| 1 | 7.277682000 | 0.725292000 | -0.339830000 |
| 6 | -1.653979000 | 0.350458000 | -0.577940000 |
| 6 | -0.648027000 | -0.032434000 | -1.651765000 |
| 6 | -1.112388000 | -2.294101000 | -1.167664000 |
| 6 | -0.781673000 | -1.709677000 | 0.186746000 |
| 6 | -1.746878000 | -0.564328000 | 0.382319000 |
| 8 | -1.065397000 | -1.378827000 | -2.117284000 |
| 1 | -0.842678000 | -2.463471000 | 0.962522000 |
| 16 | -1.473773000 | -3.860798000 | -1.482772000 |
| 6 | -2.458828000 | 1.605030000 | -0.684044000 |
| 8 | -3.479709000 | 1.685738000 | -1.325393000 |
| 8 | -1.887955000 | 2.607256000 | -0.038236000 |
| 6 | -2.586266000 | 3.880767000 | -0.073833000 |
| 1 | -2.686349000 | 4.181828000 | -1.117397000 |
| 1 | -3.579160000 | 3.727593000 | 0.351871000 |
| 6 | -2.715888000 | -0.463045000 | 1.497304000 |
| 8 | -3.432653000 | 0.500636000 | 1.663799000 |
| 8 | -2.710257000 | -1.531090000 | 2.280934000 |
| 6 | -3.633928000 | -1.511058000 | 3.384812000 |
| 1 | -3.403587000 | -0.674490000 | 4.043432000 |
| 1 | -3.489431000 | -2.455926000 | 3.900145000 |
| 6 | -0.717338000 | 0.833373000 | -2.887885000 |
| 1 | -0.468905000 | 1.867125000 | -2.643610000 |
| 1 | -0.031210000 | 0.469805000 | -3.651362000 |
| 1 | -1.733287000 | 0.802693000 | -3.284279000 |
| 1 | -4.653841000 | -1.428557000 | 3.011044000 |
| 6 | -1.767137000 | 4.860268000 | 0.727109000 |
| 1 | -2.266976000 | 5.830322000 | 0.722250000 |
| 1 | -1.667526000 | 4.527244000 | 1.761432000 |
| 1 | -0.773250000 | 4.980805000 | 0.292967000 |

**C4 (from PyrS4)**

Zero-point correction= 0.471115 (Hartree/Particle)
Thermal correction to Energy= 0.499124
Thermal correction to Enthalpy= 0.500069
Thermal correction to Gibbs Free Energy= 0.411502
Sum of electronic and zero-point Energies= -1817.151643
Sum of electronic and thermal Energies= -1817.123633
Sum of electronic and thermal Enthalpies= -1817.122689
Sum of electronic and thermal Free Energies= -1817.211256

| | | | |
|---|---|---|---|
| 6 | 1.722091000 | -0.047733000 | 0.008733000 |

| | | | |
|---|---|---|---|
| 6 | 0.468046000 | 0.007328000 | -0.464632000 |
| 6 | 2.862994000 | 0.929819000 | 0.173959000 |
| 1 | 3.212939000 | 0.871301000 | 1.208230000 |
| 1 | 3.682511000 | 0.536852000 | -0.438708000 |
| 6 | -0.538604000 | 1.072872000 | -0.847676000 |
| 1 | -1.380860000 | 0.935196000 | -0.157939000 |
| 1 | -0.926801000 | 0.804428000 | -1.836988000 |
| 6 | -0.214020000 | 2.562586000 | -0.888557000 |
| 1 | 0.502997000 | 2.768553000 | -1.682199000 |
| 1 | -1.141378000 | 3.062846000 | -1.185214000 |
| 6 | 2.673486000 | 2.404846000 | -0.152217000 |
| 1 | 2.382773000 | 2.523635000 | -1.194952000 |
| 1 | 3.655243000 | 2.877482000 | -0.049268000 |
| 6 | 1.697878000 | 3.056651000 | 0.799176000 |
| 6 | 0.239558000 | 3.115110000 | 0.438847000 |
| 6 | 1.048017000 | 4.389411000 | 0.520745000 |
| 1 | 0.856180000 | 4.998165000 | 1.398058000 |
| 1 | -0.465926000 | 2.957482000 | 1.248621000 |
| 1 | 1.927051000 | 2.869071000 | 1.843672000 |
| 6 | 1.378789000 | 5.198474000 | -0.699556000 |
| 1 | 1.720805000 | 4.572096000 | -1.526167000 |
| 1 | 2.177227000 | 5.909682000 | -0.464725000 |
| 8 | 0.200436000 | 5.914602000 | -1.093896000 |
| 1 | 0.435996000 | 6.500020000 | -1.822010000 |
| 6 | -0.079223000 | -2.194575000 | 0.628835000 |
| 6 | -0.085826000 | -1.433088000 | -0.672671000 |
| 6 | 2.099527000 | -2.163549000 | -1.082716000 |
| 6 | 2.168876000 | -1.521716000 | 0.306814000 |
| 6 | 1.141884000 | -2.273629000 | 1.128051000 |
| 8 | 0.868899000 | -2.103733000 | -1.565284000 |
| 16 | 3.327628000 | -2.830497000 | -1.922225000 |
| 8 | 3.482453000 | -1.652853000 | 0.802046000 |
| 6 | 3.728833000 | -1.339621000 | 2.096842000 |
| 8 | 2.877157000 | -0.914812000 | 2.837811000 |
| 6 | 5.163127000 | -1.566971000 | 2.434792000 |
| 1 | 5.448965000 | -2.584083000 | 2.164391000 |
| 1 | 5.771410000 | -0.876197000 | 1.846136000 |
| 1 | 1.451118000 | -2.844090000 | 1.992343000 |
| 1 | -0.975023000 | -2.650344000 | 1.025161000 |
| 6 | -1.413233000 | -1.507416000 | -1.413053000 |
| 8 | -1.518674000 | -1.719262000 | -2.594290000 |
| 8 | -2.420592000 | -1.297218000 | -0.585620000 |
| 6 | -3.737900000 | -1.187868000 | -1.191530000 |
| 1 | -3.675132000 | -0.441398000 | -1.985318000 |
| 1 | -4.001856000 | -2.156482000 | -1.615903000 |
| 6 | -4.686644000 | -0.770738000 | -0.108947000 |
| 6 | -5.546868000 | -1.697772000 | 0.475912000 |
| 6 | -4.693772000 | 0.553227000 | 0.334911000 |
| 6 | -6.410468000 | -1.306211000 | 1.496456000 |
| 6 | -5.551065000 | 0.944071000 | 1.357216000 |
| 6 | -6.411296000 | 0.013382000 | 1.938443000 |
| 1 | -5.540525000 | -2.725503000 | 0.128551000 |
| 1 | -4.026177000 | 1.274659000 | -0.126422000 |
| 1 | -7.080122000 | -2.030512000 | 1.945180000 |
| 1 | -5.553915000 | 1.972855000 | 1.697683000 |
| 1 | -7.082688000 | 0.318889000 | 2.732518000 |
| 1 | 5.320583000 | -1.391782000 | 3.495671000 |

**Synthetic details and compound characterization**

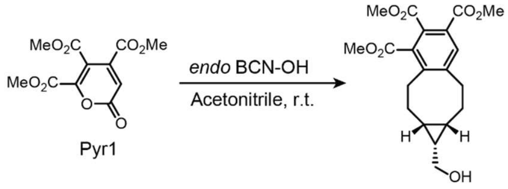


C1

**Trimethyl (1*S*,1a*R*,9a*S*)-1-(hydroxymethyl)-1a,2,3,8,9,9a-hexahydro-1*H*-benzo[*a*]cyclopropa[*e*][8]annulene-4,5,6-tricarboxylate (C1)**: To a mixture of trimethyl 2-oxo-2*H*-pyran-4,5,6-tricarboxylate (**Pyr1**) (8.3 mg, 0.031 mmol, 1 eq) and (1*R*,8*S*,9*s*)-Bicyclo[6.1.0]non-4-yn-9-ylmethanol (**BCN**) (5 mg, 0.033 mmol, 1.1 eq), acetonitrile (0.60 mL) was added, and the reaction mixture was stirred at room temperature overnight. After concentration *in vacuo*, the reaction crude mixture was purified by preparative TLC to obtain (60% EtOAc/hexanes (v/v)) to obtain **C1** (9.7 mg, 84%). $^{1}$H NMR (700 MHz, $CDCl_3$): δ = 7.70 (s, 1H), 3.88 (s, 3H), 3.87 (s, 3H), 3.87 (s, 3H), 3.70 (br, 2H), 3.06 (br, 1H), 2.96–2.88 (m, 2H), 2.80 (br, 1H), 2.35–2.30 (m, 1H), 2.26 (br, 1H), 1.61–0.86 (m, 6H).$^{13}$C NMR (176 MHz, $CDCl_3$): δ = 168.55, 168.35, 166.55, 145.67, 144.46, 133.45, 132.78, 131.01, 127.53, 59.72, 52.94, 52.76, 33.94, 29.82, 22.82, 18.67. HRMS (ESI): Calcd for $C_{20}H_{25}O_7$ $[M+H]^+$: 377.1595, found: 377.1592.

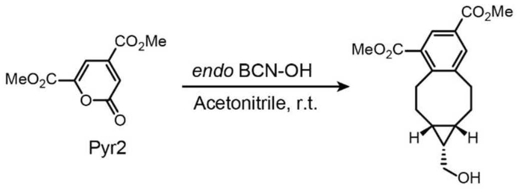


C2

**Dimethyl (1*S*,1a*R*,9a*S*)-1-(hydroxymethyl)-1a,2,3,8,9,9a-hexahydro-1*H*-benzo[*a*]cyclopropa[*e*][8]annulene-4,6-dicarboxylate (C2)**: To a mixture of dimethyl 2-oxo-2*H*-pyran-4,6-dicarboxylate (**Pyr2**) (12.1 mg, 0.057 mmol, 1 eq) and (1*R*,8*S*,9*s*)-Bicyclo[6.1.0]non-4-yn-9-ylmethanol (**BCN**) (9.4 mg, 0.063 mmol, 1.1 eq), acetonitrile (1.14 mL) was added, and the reaction mixture was stirred at room temperature overnight. After concentration *in vacuo*, the reaction crude mixture was purified by preparative TLC to obtain (40% EtOAc/hexanes (v/v)) to obtain **C2** (13.9 mg, 77%). $^{1}$H NMR (700 MHz, $CDCl_3$): δ = 8.27 (s, 1H), 7.90 (s, 1H), 3.91 (s, 6H), 3.71 (br, 2H), 3.10–2.90 (m, 4H), 2.50–2.45 (m, 1H), 2.26 (br, 1H), 1.75–0.60 (m, 6H).$^{13}$C NMR (176 MHz, $CDCl_3$): δ = 168.35, 166.66, 148.42, 143.88, 134.34, 131.46, 129.24, 127.42, 59.81, 52.35, 52.30, 34.23, 29.81, 23.03, 22.81, 17.58. HRMS (ESI): Calcd for $C_{18}H_{24}O_5$ $[M+H]^+$: 319.1540, found: 319.1540.

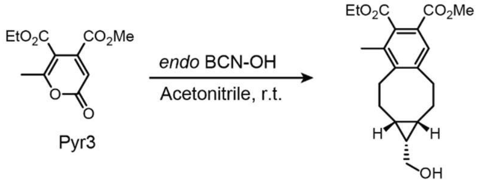


C3

**5-Ethyl 6-methyl (1*S*,1a*R*,9a*S*)-1-(hydroxymethyl)-4-methyl-1a,2,3,8,9,9a-hexahydro-1*H*-benzo[*a*]cyclopropa[*e*][8]annulene-5,6-dicarboxylate (C3)**: To a mixture of 5-ethyl 4-methyl 6-methyl-2-oxo-2*H*-pyran-4,5-dicarboxylate (**Pyr3**) (11.8 mg, 0.049 mmol, 1 eq) and (1*R*,8*S*,9*s*)-Bicyclo[6.1.0]non-4-yn-9-ylmethanol (**BCN**) (8.1 mg, 0.054 mmol, 1.1 eq), acetonitrile (0.98 mL) and dichloromethane (0.20 mL) were added, and the reaction mixture was stirred at room temperature overnight. After concentration *in vacuo*, the reaction crude mixture was purified by preparative TLC to obtain (40% EtOAc/hexanes (v/v)) to obtain **C3** (14.0 mg, 82%). $^{1}$H NMR (700 MHz, $CDCl_3$): δ = 7.62 (s, 1H), 4.44–4.40 (m, 2H), 3.85 (s, 3H), 3.75–3.68 (m, 2H), 3.05–2.84 (m, 4H), 2.29 (s, 4H), 2.24 (br, 1H), 1.47–0.87 (m, 6H), 1.38 (t, 3H).$^{13}$C NMR (176 MHz, $CDCl_3$): δ = 170.14, 166.56, 146.48, 142.81, 134.34, 133.12, 129.85, 124.61, 61.56, 59.77, 52.33, 34.36, 29.82, 27.40, 22.81, 16.37, 14.26. HRMS (ESI): Calcd for $C_{20}H_{27}O_5$ $[M+H]^+$: 347.1853, found: 347.1850.

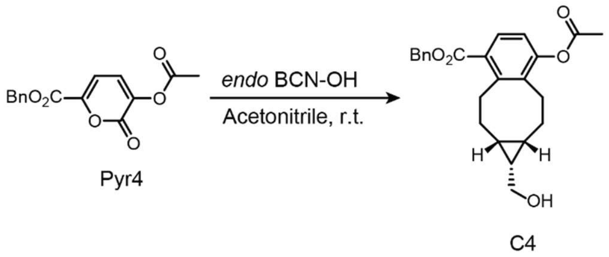

**Benzyl (1*R*,1a*R*,9a*S*)-7-acetoxy-1-(hydroxymethyl)-1a,2,3,8,9,9a-hexahydro-1*H*-benzo[*a*]cyclopropa[*e*][8]annulene-4-carboxylate (C4)**: To a mixture of benzyl 3-acetoxy-2-oxo-2*H*-pyran-6-carboxylate (**Pyr4**) (9.5 mg, 0.033 mmol, 1 eq) and (1*R*,8*S*,9*s*)-Bicyclo[6.1.0]non-4-yn-9-ylmethanol (**BCN**) (5.4 mg, 0.036 mmol, 1.1 eq), acetonitrile (0.66 mL) was added, and the reaction mixture was stirred at room temperature overnight. After concentration *in vacuo*, the reaction crude mixture was purified by preparative TLC to obtain (40% EtOAc/hexanes (v/v)) to obtain **C4** (11.4 mg, 88%). $^{1}$H NMR (700 MHz, $CDCl_3$): δ = 7.68 (br, 1H), 7.44–7.43 (m, 2H), 7.41–7.38 (m, 2H), 7.35–7.33 (m, 1H), 5.33 (s, 2H), 3.71 and 3.46 (br, 3H), 2.97–2.76 (m, 2H), 2.43–2.35 (m, 1H), 2.34 (s, 3H), 2.18 (br, 1H), 1.72–0.61 (m, 6H). $^{13}$C NMR (176 MHz, $CDCl_3$): δ = 169.44, 167.87, 151.59, 136.09. 129.16, 128.75, 128.42, 128.37, 119.79, 66.90, 59.89, 31.02, 29.83, 27.80, 25.89, 24.29, 22.82, 21.89, 21.03, 16.18. HRMS (ESI): Calcd for $C_{24}H_{27}O_5$ $[M+H]^+$: 395.1853, found: 395.1851.

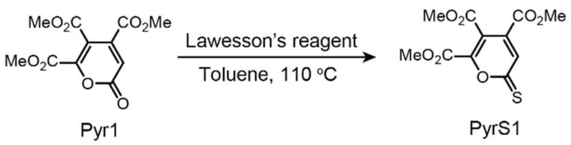

**Trimethyl 2-thioxo-2*H*-pyran-4,5,6-tricarboxylate (PyrS1)**: To a mixture of trimethyl 2-oxo-2*H*-pyran-4,5,6-tricarboxylate (**Pyr1**) (65 mg, 0.24 mmol, 1 eq) and Lawesson's reagent (97 mg, 0.24 mmol, 1.0 eq), anhydrous toluene (4.0 mL) was added, and the reaction mixture was stirred at 110 °C for 2 days under nitrogen. After cooling to room temperature, the reaction crude mixture was directly loaded on silica, and initially purified by flash column (5% EtOAc/hexanes (v/v)) then by preparative TLC (20% EtOAc/hexanes (v/v)) to obtain **PyrS1** (30 mg, 44%). $^{1}$H NMR (500 MHz, $CDCl_3$): δ = 7.75 (s, 1H), 3.96 (d, *J* = 2.8 Hz, 6H) and 3.91 (s, 3H). $^{13}$C NMR (126 MHz, $CDCl_3$): δ = 193.26, 163.98, 162.73, 158.50, 150.69, 136.46, 130.92, 119.95, 53.95, 53.88, 53.68. HRMS (ESI): Calcd for $C_{11}H_{11}O_7S$ $[M+H]^+$: 287.0220, found: 287.0218.

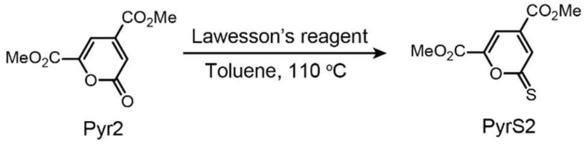

**Dimethyl 2-thioxo-2*H*-pyran-4,6-dicarboxylate (PyrS2)**: To a mixture of dimethyl 2-oxo-2*H*-pyran-4,6-dicarboxylate (**Pyr2**) (58 mg, 0.27 mmol, 1 eq) and Lawesson's reagent (111 mg, 0.27 mmol, 1.0 eq), anhydrous toluene (5.4 mL) was added, and the reaction mixture was stirred at 110 °C for 2 days under nitrogen. After cooling to room temperature, the reaction crude mixture was then directly loaded on silica, and initially purified by flash column (5% EtOAc/hexanes (v/v)) to obtain **PyrS2** (44 mg, 71%). $^{1}$H NMR (500 MHz, $CDCl_3$): δ = 7.78 (d, *J* = 1.4 Hz, 1H), 7.62 (d, *J* = 1.4 Hz, 1H), 3.97 (s, 3H), 3.95 (s, 3H). $^{13}$C NMR (126 MHz, $CDCl_3$): δ = 195.34, 163.64, 159.02, 153.17, 136.30, 132.98, 111.73, 53.70, 53.60. HRMS (ESI): Calcd for $C_9H_9O_5S$ $[M+H]^+$: 229.0165, found: 229.0162.

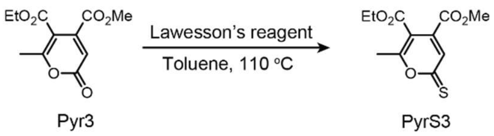

**5-Ethyl 4-methyl 6-methyl-2-thioxo-2*H*-pyran-4,5-dicarboxylate (PyrS3)**: To a mixture of 5-ethyl 4-methyl 6-methyl-2-oxo-2*H*-pyran-4,5-dicarboxylate (**Pyr3**) (48.2 mg, 0.20 mmol, 1 eq) and Lawesson's reagent (81.0 mg, 0.20 mmol, 1.0 eq), anhydrous toluene (2.0 mL) was added, and the reaction mixture was stirred at 110 °C for overnight. After cooling to room temperature, the reaction crude mixture was directly loaded on silica, and initially purified by preparative TLC (15% EtOAc/hexanes (v/v)) to obtain **PyrS3** (30.0 mg, 59%). $^{1}$H NMR (500 MHz, $CDCl_3$): δ = 7.29 (s, 1H), 4.31 (q, 2H), 3.86 (s, 3H), 2.51 (s, 3H), 1.32 (t, 3H). $^{13}$C NMR (100 MHz, $CDCl_3$): δ = 195.79, 169.39, 164.85, 164.26, 135.65, 129.04, 112.79, 62.42, 53.35, 19.47, 14.05. HRMS (ESI): Calcd for $C_{11}H_{13}O_5S$ $[M+H]^+$: 257.0478, found: 257.0472.

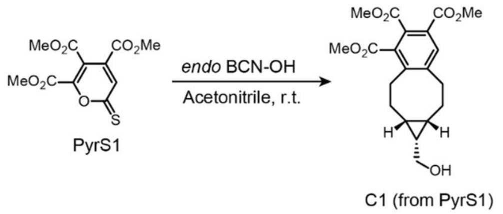

**Trimethyl (1*S*,1a*R*,9a*S*)-1-(hydroxymethyl)-1a,2,3,8,9,9a-hexahydro-1*H*-benzo[*a*]cyclopropa[*e*][8]annulene-4,5,6-tricarboxylate (C1)**: To a mixture of trimethyl 2-thioxo-2*H*-pyran-4,5,6-tricarboxylate (**PyrS1**) (7.0 mg, 0.024 mmol, 1 eq) and (1*R*,8*S*,9*s*)-Bicyclo[6.1.0]non-4-yn-9-ylmethanol (**BCN**) (8.2 mg, 0.054 mmol, 2.2 eq), acetonitrile (0.68 mL) was added, and the reaction mixture was stirred at room temperature overnight. After concentration *in vacuo*, the reaction crude mixture was purified by preparative TLC to obtain (60% EtOAc/hexanes (v/v)) to obtain **C1** (7.7 mg, 85%). $^{1}$H NMR (700 MHz, $CDCl_3$): δ = 7.70 (s, 1H), 3.88 (s, 3H), 3.87 (s, 3H), 3.87 (s, 3H), 3.70 (br, 2H), 3.06 (br, 1H), 2.96–2.88 (m, 2H), 2.80 (br, 1H), 2.35–2.30 (m, 1H), 2.26 (br, 1H), 1.61–0.86 (m, 6H). $^{13}$C NMR (176 MHz, $CDCl_3$): δ = 168.55, 168.35, 166.55, 145.67, 144.46, 133.45, 132.78, 131.01, 127.53, 59.73, 52.94, 52.76, 33.95, 29.83, 22.82, 22.29, 18.67. HRMS (ESI): Calcd for $C_{20}H_{25}O_7$ $[M+H]^+$: 377.1595, found: 377.1592.

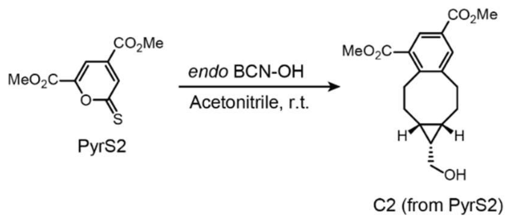

**Dimethyl (1*S*,1a*R*,9a*S*)-1-(hydroxymethyl)-1a,2,3,8,9,9a-hexahydro-1*H*-benzo[*a*]cyclopropa[*e*][8]annulene-4,6-dicarboxylate (C2)**: To a mixture of dimethyl 2-thioxo-2*H*-pyran-4,6-dicarboxylate (**PyrS2**) (4.7 mg, 0.021 mmol, 1 eq) and (1*R*,8*S*,9*s*)-Bicyclo[6.1.0]non-4-yn-9-ylmethanol (**BCN**) (3.4 mg, 0.023 mmol, 1.1 eq), acetonitrile (0.40 mL) was added, and the reaction mixture was stirred at room temperature overnight. After concentration *in vacuo*, the reaction crude mixture was purified by preparative TLC to obtain (40% EtOAc/hexanes (v/v)) to obtain **C2** (6.8 mg, quantitative). $^{1}$H NMR (700 MHz, $CDCl_3$): δ = 8.27 (s, 1H), 7.90 (s, 1H), 3.91 (s, 6H), 3.71 (br, 2H), 3.10–2.90 (m, 4H), 2.50–2.45 (m, 1H), 2.26 (br, 1H), 1.75–0.60 (m, 6H). $^{13}$C NMR (176 MHz, $CDCl_3$): δ = 168.37, 166.67, 148.45, 144.06, 134.36, 131.48, 129.27, 127.45, 59.86, 52.37, 52.31, 34.28, 29.83, 22.83, 17.47. HRMS (ESI): Calcd for $C_{18}H_{26}NO_5$ $[M+NH_4]^+$: 336.1805, found: 336.1810.

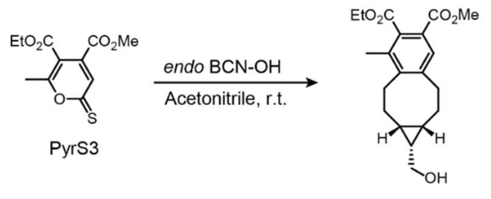


**5-Ethyl 6-methyl (1*S*,1a*R*,9a*S*)-1-(hydroxymethyl)-4-methyl-1a,2,3,8,9,9a-hexahydro-1*H*-benzo[*a*]cyclopropa[*e*][8]annulene-5,6-dicarboxylate (C3)**: To a mixture of 5-ethyl 4-methyl 6-methyl-2-thioxo-2*H*-pyran-4,5-dicarboxylate (**PyrS3**) (8.7 mg, 0.034 mmol, 1 eq) and (1*R*,8*S*,9*s*)-Bicyclo[6.1.0]non-4-yn-9-ylmethanol (**BCN**) (8.1 mg, 0.054 mmol, 1.6 eq), acetonitrile (0.88 mL) was added, and the reaction mixture was stirred at room temperature overnight. After concentration *in vacuo*, the reaction crude mixture was purified by preparative TLC to obtain (40% EtOAc/hexanes (v/v)) to obtain **C3** (9.3 mg, 79%). $^{1}$H NMR (700 MHz, $CDCl_3$): δ = 7.62 (s, 1H), 4.44–4.40 (m, 2H), 3.85 (s, 3H), 3.75–3.68 (m, 2H), 3.05–2.84 (m, 4H), 2.29 (s, 4H), 2.24 (br, 1H), 1.47–0.87 (m, 6H), 1.38 (t, 3H).$^{13}$C NMR (176 MHz, $CDCl_3$): δ = 170.42, 166.56, 146.45, 143.17, 134.35, 133.13, 129.87, 124.62, 61.56, 59.88, 52.33, 34.37, 29.83, 27.47, 22.82, 16.33, 14.27. HRMS (ESI): Calcd for $C_{20}H_{27}O_5$ $[M+NH_4]^+$: 364.2118, found: 364.2114.

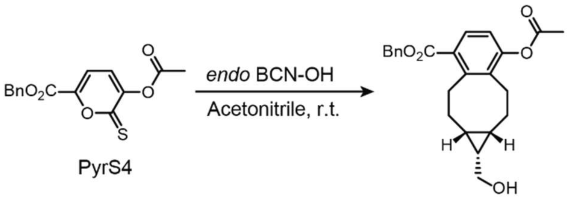


**Benzyl (1*R*,1a*R*,9a*S*)-7-acetoxy-1-(hydroxymethyl)-1a,2,3,8,9,9a-hexahydro-1*H*-benzo[*a*]cyclopropa[*e*][8]annulene-4-carboxylate (C4)**: To a mixture of benzyl 3-acetoxy-2-thioxo-2*H*-pyran-6-carboxylate (**PyrS4**) (6.7 mg, 0.022 mmol, 1 eq) and (1*R*,8*S*,9*s*)-Bicyclo[6.1.0]non-4-yn-9-ylmethanol (**BCN**) (8.0 mg, 0.053 mmol, 2.4 eq), acetonitrile (0.44 mL) was added, and the reaction mixture was stirred at room temperature for 2 days. After concentration *in vacuo*, the reaction crude mixture was purified by preparative TLC to obtain (40% EtOAc/hexanes (v/v)) to obtain **C4** (7.5 mg, 86%). $^{1}$H NMR (700 MHz, $CDCl_3$): δ = 7.68 (br, 1H), 7.44–7.43 (m, 2H), 7.41–7.38 (m, 2H), 7.35–7.33 (m, 1H), 5.33 (s, 2H), 3.71 and 3.46 (br, 3H), 2.97–2.76 (m, 2H), 2.43–2.35 (m, 1H), 2.34 (s, 3H), 2.18 (br, 1H), 1.72–0.61 (m, 6H). $^{13}$C NMR (176 MHz, $CDCl_3$): δ = 169.45, 167.88, 151.60, 136.10. 129.16, 128.75, 128.43, 128.38, 119.79, 66.91, 59.89, 29.83, 27.80, 25.94, 24.30, 22.83, 21.97, 21.03, 16.56. HRMS (ESI): Calcd for $C_{24}H_{30}NO_5$ $[M+NH_4]^+$: 412.2118, found: 412.2120.

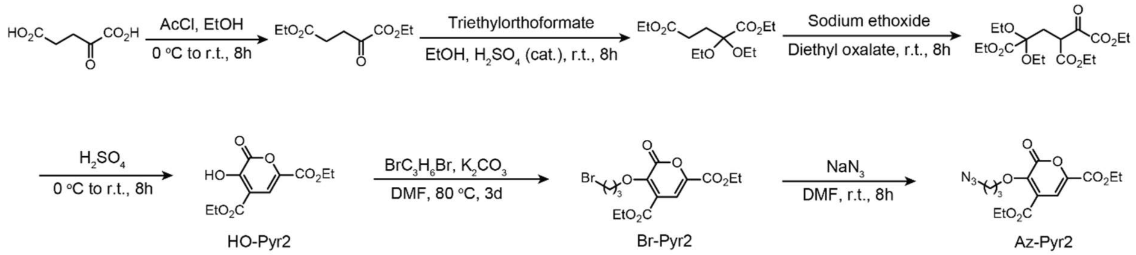

**Scheme S1** Synthetic scheme for clickable pyrone2 for further bioconjugation.

**Diethyl 3-hydroxy-2-oxo-2*H*-pyran-4,6-dicarboxylate (HO-Pyr2)** was synthesized according to the previous report starting from α-ketoglutaric acid[12].

**Diethyl 3-(3-bromopropoxy)-2-oxo-2*H*-pyran-4,6-dicarboxylate (Br-Pyr2)**: To a solution of diethyl 3-hydroxy-2-oxo-2*H*-pyran-4,6-dicarboxylate (**HO-Pyr2**) (1.46 g, 5.69 mmol, 1 eq) in 1,3-dibromopropane (12 mL) and anhydrous DMF (1.2 mL), potassium carbonate (1.57 g, 11.4 mmol, 2 eq) was added in one portion. The mixture was stirred at 80 °C for 4 days under nitrogen and then concentrated *in vacuo*. The resulting crude was purified by flash column (5% EtOAc/hexanes (v/v)) to obtain **Br-Pyr2** (1.12 g, 52%). $^{1}$H NMR (500 MHz, $CDCl_3$): δ = 7.36 (s, 1H), 4.55 (t, 2H), 4.41–4.36 (m, 4H), 3.62 (q, 2H), 2.32–2.28 (m, 2H), 1.39 (q, 6H). $^{13}$C NMR (126 MHz, $CDCl_3$): δ = 162.77, 159.01, 157.48, 148.02, 142.61, 126.42, 110.94, 71.71, 62.62, 62.60, 33.07, 29.63, 14.31, 14.30. HRMS (ESI): Calcd for $C_{14}H_{18}BrO_7$ $[M+H]^+$: 377.0230, found: 377.0221.

**Diethyl 3-(3-azidopropoxy)-2-oxo-2*H*-pyran-4,6-dicarboxylate (Az-Pyr2)**: To an ice-cold solution of diethyl 3-(3-bromopropoxy)-2-oxo-2*H*-pyran-4,6-dicarboxylate (**Br-Pyr2**) (143 mg, 0.38 mmol, 1 eq) in DMF (1.9 mL), sodium azide (25 mg, 0.38 mmol, 1 eq) was added in one portion. The mixture was warmed to room temperate and stirred overnight. After concentrated *in vacuo*, the resulting crude was purified by flash column (5% to 10% EtOAc/hexanes (v/v)) to obtain **Az-Pyr2** (77 mg, 60%). $^{1}$H NMR (700 MHz, $CDCl_3$): δ = 7.32 (s, 1H), 4.55 (t, 2H), 4.37–4.33 (m, 4H), 3.52 (q, 2H), 2.02–1.98 (m, 2H), 1.36 (q, 6H). $^{13}$C NMR (176 MHz, $CDCl_3$): δ = 162.57, 158.88, 157.39, 147.92, 142.55, 126.54, 110.77, 70.81, 62.51, 62.44, 47.84, 29.48, 14.18. HRMS (ESI): Calcd for $C_{14}H_{18}N_3O_7$ $[M+H]^+$: 340.1139, found: 340.1128.

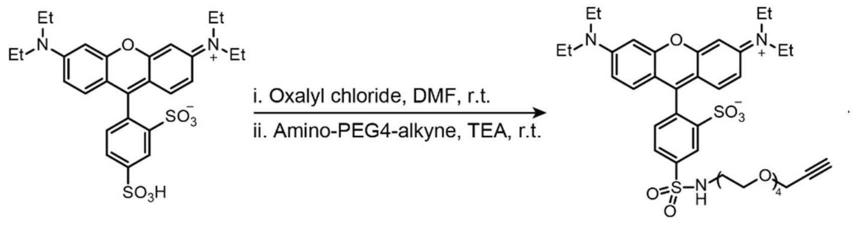

Sulforhodamine B-PEG4-alkyne

**Sulforhodamine B-PEG4-alkyne**: **Sulforhodamine B sulfonyl chloride** was prepared according to literature. To an ice-cold solution of sulforhodamine B sulfonyl chloride (25 mg, 0.043 mmol, 1 eq) and amino-PEG4-alkyne (Sigma-Aldrich #764248) (10 mg, 0.043 mmol, 1 eq) in anhydrous DMF (0.40 mL), DIPEA (14.7 µL, 0.086 mmol, 2 eq) was added. The mixture was warmed to room temperate and stirred overnight. After concentration *in vacuo*, the resulting crude was purified by flash column (1% to 4% MeOH/DCM (v/v)) to obtain **Sulforhodamine B-PEG4-alkyne** (5.9 mg, 18%). $^1$H NMR (500 MHz, $CDCl_3$): δ = 8.84 (s, 1H), 7.98 (d, *J* = 7.7 Hz, 1H), 7.28 (d, *J* = 10.3 Hz, 2H), 7.20 (d, *J* = 7.8 Hz, 1H), 6.80 (d, *J* = 9.0 Hz, 2H), 6.66 (s, 2H), 5.67 (br, 1H), 4.20 (s, 2H), 3.71–3.53 (m, 22H), 3.28–3.27 (m, 2H), 2.43 (s, 1H), 1.29 (t, 12H). $^{13}$C NMR (126 MHz, $CDCl_3$): δ = 159.36, 158.04, 155.64, 148.58, 141.96, 133.79, 133.66, 129.81, 127.61, 127.02, 114.52, 113.60, 95.75, 79.88, 74.67, 70.68, 70.65, 70.62, 70.48, 69.68, 69.27, 58.51, 45.97, 43.39, 29.81, 12.72. HRMS (ESI): Calcd for $C_{38}H_{50}N_3O_{10}S_2$ $[M+H]^+$: 772.2932, found: 772.2928.

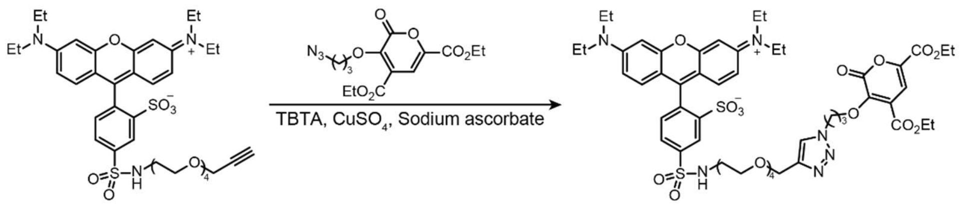

Sulforhodamine B-PEG4-alkyne | Sulforhodamine B-PEG4-pyrone

**Sulforhodamine B-PEG4-pyrone**: To a solution of **sulforhodamine B-PEG4-alkyne** (5.9 mg, 7.6 µmol, 1 eq) and **Az-Pyr2** (3.8 mg, 11.2 µmol, 1.5 eq) in DMSO (0.42 mL), TBTA (20 mM, 60 µL, 0.16 eq) and $CuSO_4$ (20 mM, 60 µL, 0.16 eq) were added. Finally, sodium ascorbate (0.5 M, 100 µL, 6.6 eq) was added and the mixture was stirred overnight at room temperature. The crude was purified by reversed-phase HPLC to obtain **Sulforhodamine B-PEG4-pyrone** (6.2 mg, 73%). $^1$H NMR (700 MHz, $CDCl_3$): δ = 8.62 (s, 1H), 8.01–8.00 (m, 1H), 7.90 (s, 1H), 7.38 (s, 1H), 7.21–7.17 (m, 3H), 6.82 (d, *J* = 8.3 Hz, 2H), 6.66 (d, *J* = 2.2 Hz, 2H), 4.67 (br, 2H), 4.42–4.36 (m, 6H), 3.68–3.52 (m, 18H), 3.29 (br, 2H), 2.38 (t, 2H), 1.40–1.37 (m, 6H), 1.29 (t, 12H). $^{13}$C NMR (176 MHz, $CDCl_3$): δ = 162.65, 159.03, 157.95, 157.75, 155.71, 148.09, 144.77, 142.71, 142.42, 133.60, 133.29, 130.11, 127.77, 127.16, 127.11, 123.91, 114.34, 113.82, 111.01, 95.82, 70.59, 70.04, 69.92, 69.82, 69.68, 69.59, 68.94, 64.20, 62.77, 62.64, 46.94, 46.01, 42.85, 30.74, 29.85, 14.31, 12.69. HRMS (ESI): Calcd for $C_{52}H_{67}N_6O_{17}S_2$ $[M+H]^+$: 1111.3999, found: 1111.3997.

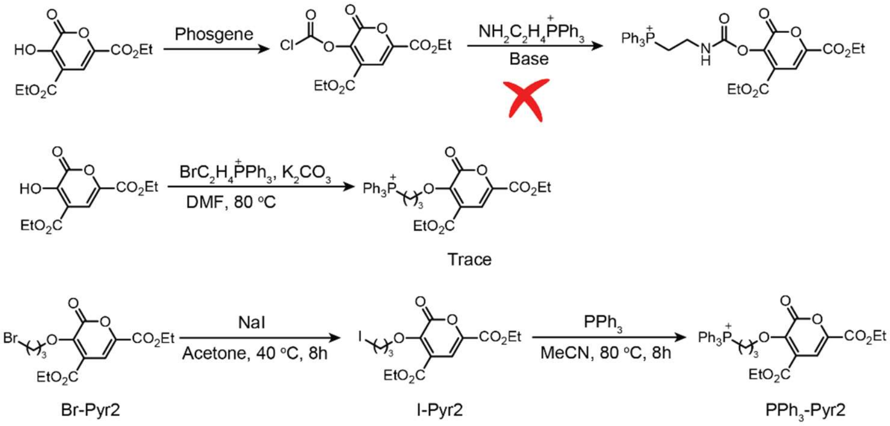

**Scheme S2** Synthetic trails for the preparation of mitochondrial targeting Pyr2 analogue.

**Diethyl 3-(3-iodopropoxy)-2-oxo-2*H*-pyran-4,6-dicarboxylate (I-Pyr2)**: To a solution of diethyl 3-(3-bromopropoxy)-2-oxo-2*H*-pyran-4,6-dicarboxylate (**Br-Pyr2**) (19 mg, 0.050 mmol, 1 eq) in acetone (0.40 mL), sodium iodine (15.1 mg, 0.10 mmol, 2 eq) was added and the mixture was stirred at 40 °C for overnight. The resulting mixture was poured into water and extracted three times with DCM. The combined organic layers were dried over $Na_2SO_4$, concentrated *in vacuo*, and used for next step without further purification. The formation of **I-Pyr2** (22.5 mg) was confirmed by $^1$H NMR (400 MHz, $CDCl_3$): δ = 7.33 (s, 1H), 4.46 (t, 2H), 4.39–4.32 (m, 4H), 3.36 (t, 2H), 2.26–2.20 (m, 2H), 1.37 (q, 6H).

**(3-((4,6-Bis(ethoxycarbonyl)-2-oxo-2*H*-pyran-3-yl)oxy)propyl)triphenylphosphonium (PPh$_3$-Pyr2)**: To a solution of diethyl 3-(3-iodopropoxy)-2-oxo-2*H*-pyran-4,6-dicarboxylate (**I-Pyr2**) (crude from last step, 85 mg, 0.20 mmol, 1 eq) in anhydrous acetonitrile (2.0 mL), triphenylphosphine (53 mg, 0.20 mmol, 1 eq) was added and the mixture was stirred at 80 °C for overnight under nitrogen. After cooled to room temperature and concentration *in vacuo*, the resulting crude was purified by flash column (1% to 4% MeOH/DCM (v/v)) to obtain **PPh$_3$-Pyr2** (80 mg, 62% over 2 steps). $^1$H NMR (400 MHz, $CDCl_3$): δ = 7.82–7.76 (m, 9H), 7.69–7.67 (m, 1H), 7.31 (s, 1H), 4.65 (br, 2H), 4.34–4.25 (m, 4H), 3.99 (t, 2H), 2.13 (br, 2H), 1.34–1.27 (m, 6H). $^{13}$C NMR (100 MHz, $CDCl_3$): δ = 162.04, 158.78, 157.53, 148.34, 142.22, 135.21, 135.18, 133.75, 133.65, 130.63, 130.51, 125.90, 118.29, 117.43, 110.69, 73.28, 73.12, 62.56, 62.47, 23.52, 19.55, 19.02, 14.20, 14.14. HRMS (ESI): Calcd for $C_{32}H_{32}O_7P$ $[M]^+$: 559.1880, found: 559.1869. Note: **PPh$_3$-Pyr2** was further purified by HPLC (50% to 70% acetonitrile/$H_2O$ w 0.1% TFA) and lyophilized before used in cell imaging experiments.

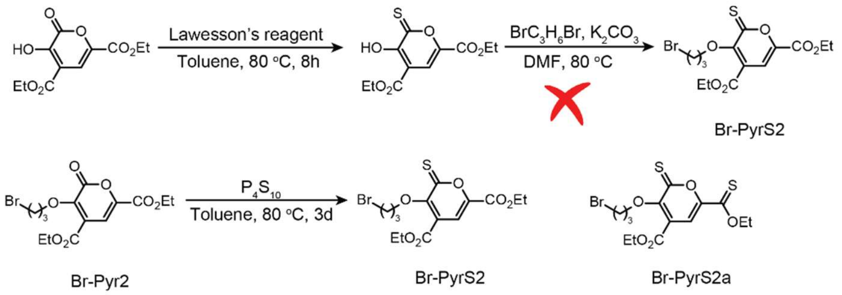


**Scheme S3** Synthetic trails for the preparation of functionalized PyrS2 analogue.

**Diethyl 3-(3-bromopropoxy)-2-thioxo-2*H*-pyran-4,6-dicarboxylate (Br-PyrS2)** and **ethyl 3-(3-bromopropoxy)-6-(ethoxycarbonothioyl)-2-thioxo-2*H*-pyran-4-carboxylate (Br-PyrS2a)**: To a solution of diethyl 3-(3-bromopropoxy)-2-oxo-2*H*-pyran-4,6-dicarboxylate (**Br-Pyr2**) (738 mg, 1.96 mmol, 1 eq) in anhydrous toluene (10 mL), $P_4S_{10}$ (870 mg, 1.96 mmol, 1 eq) was added and the mixture was stirred at 80 °C for 3 days under nitrogen. After cooling to room temperature and concentration *in vacuo*, the resulting crude was purified by flash column (5% EtOAc/hexanes (v/v)) to obtain **Br-PyrS2** (315 mg, 41%) and **Br-PyrS2a** (62 mg, 7.7%). **Br-PyrS2** $^1H$ NMR (700 MHz, $CDCl_3$): δ = 7.46 (s, 1H), 4.40 (q, 4H), 4.32 (t, 2H), 3.64 (t, 2H), 2.37–2.34 (m, 2H), 1.39 (q, 6H). $^{13}C$ NMR (176 MHz, $CDCl_3$): δ = 191.14, 163.27, 158.35, 156.31, 147.96, 122.99, 113.51, 72.01, 62.87, 33.21, 29.92, 14.26. HRMS (ESI): Calcd for $C_{14}H_{18}BrO_6S$ $[M+H]^+$: 393.0002, found: 392.9993. **Br-PyrS2a** $^1H$ NMR (700 MHz, $CDCl_3$): δ = 7.60 (s, 1H), 4.71 (q, 2H), 4.43 (q, 2H), 4.35 (t, 2H), 3.66 (t, 2H), 2.39–2.35 (m, 2H), 1.51 (t, 3H), 1.42 (t, 3H). $^{13}C$ NMR (176 MHz, $CDCl_3$): δ = 196.76, 190.84, 163.62, 155.90, 153.22, 124.21, 112.07, 71.94, 69.47, 62.94, 33.31, 29.98, 14.34, 13.77. HRMS (ESI): Calcd for $C_{14}H_{18}BrO_5S_2$ $[M+H]^+$: 408.9774, found: 408.9758. Note: we also tried Lawesson's reagent to perform the thiolation but did not observe any conversion of **Br-Pyr2**.

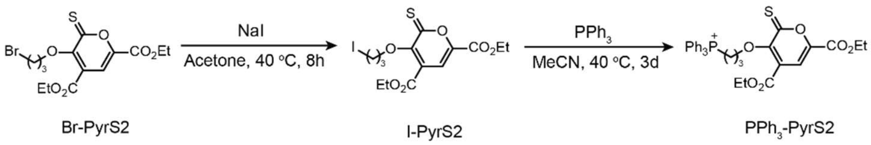

**Scheme S4** Synthetic scheme for the preparation of mitochondrial targeting PyrS2 analogues, $PPh_3$-PyS2.

**Diethyl 3-(3-iodopropoxy)-2-thioxo-2*H*-pyran-4,6-dicarboxylate (I-PyrS2)**: To a solution of diethyl 3-(3-bromopropoxy)-2-thioxo-2*H*-pyran-4,6-dicarboxylate (**Br-PyrS2**) (38 mg, 0.097 mmol, 1 eq) in acetone (0.40 mL), sodium iodine (30 mg, 0.20 mmol, 2 eq) was added and the mixture was stirred at 40 °C for overnight. The resulting mixture was poured into water and extracted three times with DCM. The combined organic layers were dried over $Na_2SO_4$, concentrated *in vacuo*, and used for next step without further purification. The formation of **I-PyrS2** (40 mg) was confirmed by NMRs. $^1$H NMR (700 MHz, $CDCl_3$): δ = 7.46 (s, 1H), 4.42–4.38 (m, 4H), 4.26 (t, 2H), 3.41 (t, 2H), 2.35–2.31 (m, 2H), 1.41–1.24 (m, 6H). $^{13}$C NMR (176 MHz, $CDCl_3$): δ = 191.11, 163.27, 158.37, 156.32, 147.96, 122.96, 113.52, 73.91, 62.89, 33.94, 14.31, 14.27, 2.33.

**(3-((4,6-bis(ethoxycarbonyl)-2-thioxo-2*H*-pyran-3-yl)oxy)propyl)triphenylphosphonium ($PPh_3$-PyrS2)**: To a solution of diethyl 3-(3-iodopropoxy)-2-thioxo-2*H*-pyran-4,6-dicarboxylate (**I-PyrS2**) (crude from last step, 39 mg, 0.089 mmol, 1 eq) in anhydrous acetonitrile (2.0 mL), triphenylphosphine (23 mg, 0.089 mmol, 1 eq) was added and the mixture was stirred at 40 °C for 3 days under nitrogen. After cooled to room temperature and concentration *in vacuo*, the resulting crude was purified by preparative TLC (4% MeOH/DCM (v/v)) to obtain **$PPh_3$-PyrS2** (11 mg, 16% over 2 steps). $^1$H NMR (700 MHz, $CD_3CN$): δ = 7.89–7.72 (m, 15H), 7.46 (s, 1H), 4.37 (q, 2H), 4.32–4.29 (m, 4H), 3.58–3.53 (m, 2H), 2.12–2.10 (m, 2H), 1.35 (t, 3H), 1.29 (t, 3H). $^{13}$C NMR (176 MHz, $CD_3CN$): δ = 193.41, 164.05, 159.16, 156.76, 149.03, 136.23, 136.21, 134.72, 134.66, 131.36, 131.28, 124.58, 119.26, 118.76, 114.45, 74.13, 74.03, 63.77, 63.70, 55.32, 24.01, 23.99, 20.07, 19.76, 14.32. HRMS (ESI): Calcd for $C_{32}H_{32}O_6PS$ $[M]^+$: 575.1652, found: 575.1643. Note: we also tried the reactions under 60 °C and 75 °C to improve reaction conversions but found that side reactions happened severely at higher reaction temperatures and **$PPh_3$-PyrS2** was further purified by HPLC (50% to 70% acetonitrile/$H_2O$ w 0.1% TFA) and lyophilized before used in cell imaging experiments.

**$2^{nd}$ order rate constant determination between pyrones and BCN**.

Reaction mixtures consisting of pyrones (0.1 mM for **Pyr1** and 1 mM for **Pyr2** and **Pyr3**) and BCN (1, 1.5, 2 mM for **Pyr1**, and 10, 15, 20 mM for **Pyr2** and **Pyr3**) in 50% acetonitrile/water were prepared in separate tubes at room temperature. Reaction aliquots (20 µL) were taken with different reaction time (2, 4, 8, 10, and 16 min for **Pyr1**, 1, 2, 4, 6, 10 min for **Pyr2**, and 1, 41, 81, and 161 min for **Pyr3**), diluted with 50% acetonitrile/water (180 or 200 µL), and injected into HPLC (time program, acetonitrile 50% to 100% in 25 minutes). The concentrations of pyrones were determined from the integration of the peaks of pyrones relative to pyrones control. The pseudo $1^{st}$ order rate constants were derived from the linear fitting of ln[pyrone] against the reaction time. The $2^{nd}$ rate constant of **Pyr4** will be reported later by our lab[13] using $^{1}H$ NMR in 50% $CD_3CN/D_2O$.

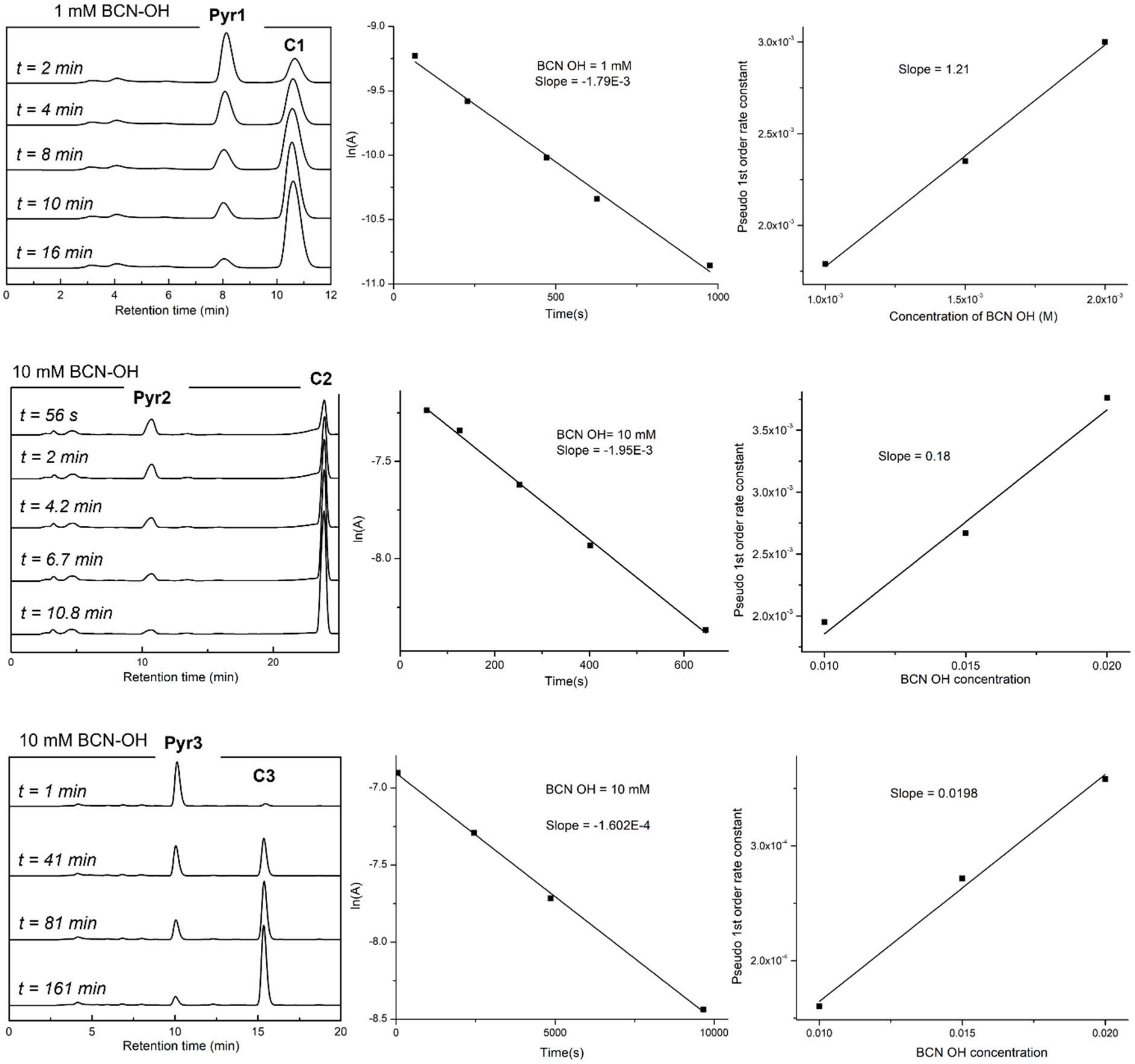


**FigureS1** Representative HPLC traces and linear fittings for 2$^{nd}$ order rate constant determination between BCN and Pyr1, Pyr2, and Pyr3, respectively.

**2$^{nd}$ order rate constant determination between pyranthiones and *endo* BCN**: Reaction mixtures consisting of pyranthiones (0.1 mM for **PyrS1**, 1 mM for **PyrS2** and **PyrS3**) and BCN (1, 1.5, 2 mM for **PyrS1**, 10, 15, and 20 mM for **PyrS2** and **PyrS3**) in 50% acetonitrile/water were prepared in separate wells of a 96-well plate at room temperature. The reaction was monitored by recording the disappearance of the characteristic absorption of pyranthiones at 412 nm every 2 or 4 minutes. The pseudo 1$^{st}$ order rate constants were derived from an exponential fitting by plotting the absorbance against the reaction time. The 2$^{nd}$ order rate constants were derived from linear fitting of the pseudo 1$^{st}$ order rates with the concentration of BCN. The 2$^{nd}$ rate constant of **PyrS4** has been reported by our recent work[14].

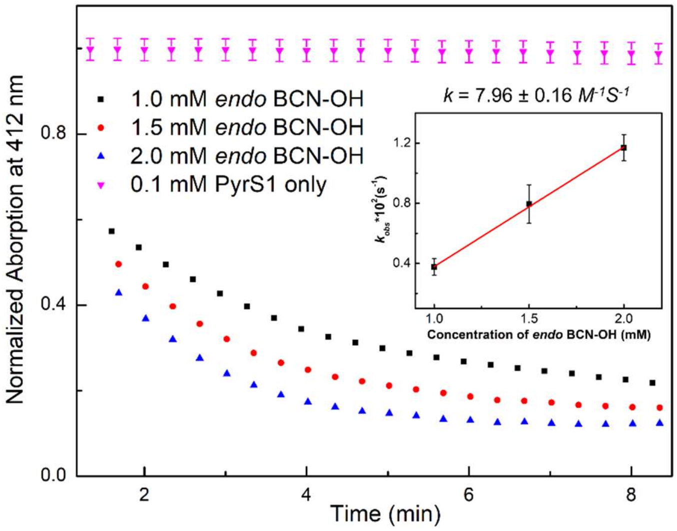


**Figure S2** Kinetics evaluation of ligation between PyrS1 and BCN (1, 1.5, and 2 mM) with measured absorption at 412 nm by plate reader.

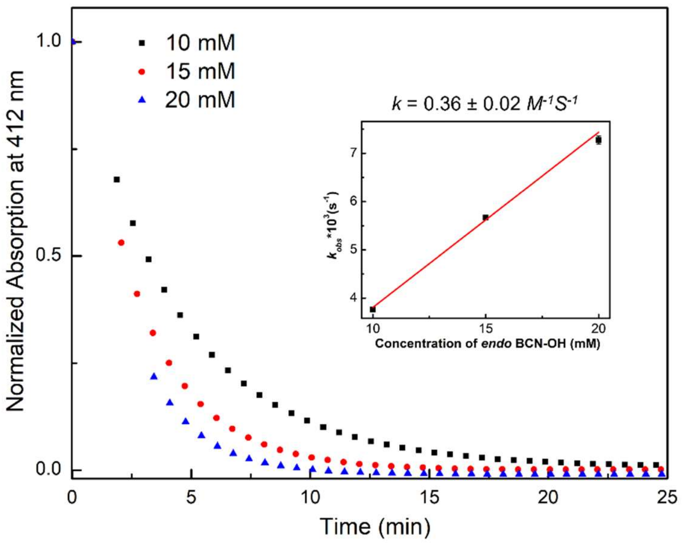


**Figure S3** Kinetics evaluation of ligation between PyrS2 and BCN (10, 15, and 20 mM) with measured absorption at 412 nm by plate reader.

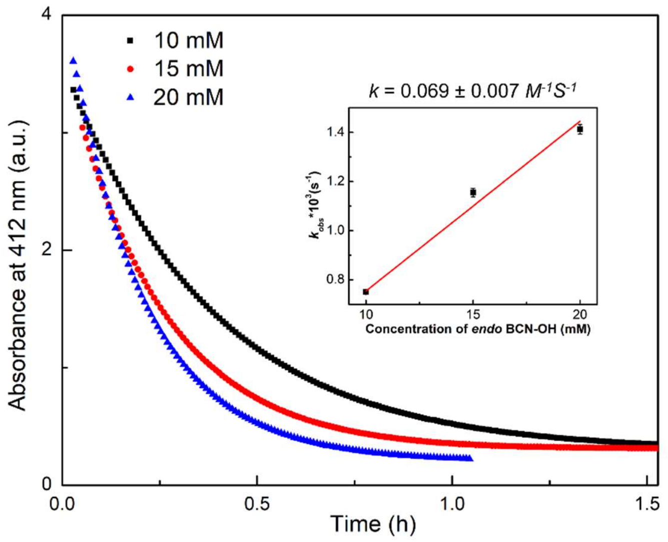


**Figure S4** Kinetics evaluation of ligation between PyrS3 and BCN (10, 15, and 20 mM) with measured absorption at 412 nm by plate reader.

**Bioorthogonality tests between pyrones and amino acids.**

Reaction mixtures consisting of pyrones (0.1 mM for Pyr1 and 1 mM for Pyr2) and amino acids (glutathione, L-lysine, L-serine, each with final concentration of 1, 1.5, 2 mM for **Pyr1**, and 10, 15, 20 mM for **Pyr2**) in 50% acetonitrile/PBS (1X) were prepared in separate tubes at room temperature. Reaction aliquots (20 μL) were taken with different reaction time (1 and 30, 35 or 38 min), diluted with 50% acetonitrile/water (180 or 200 μL), and injected into HPLC (time program, acetonitrile 50% to 100% in 25 minutes). The concentrations of pyrones were determined from the integration of the peaks of pyrones relative to pyrones control.

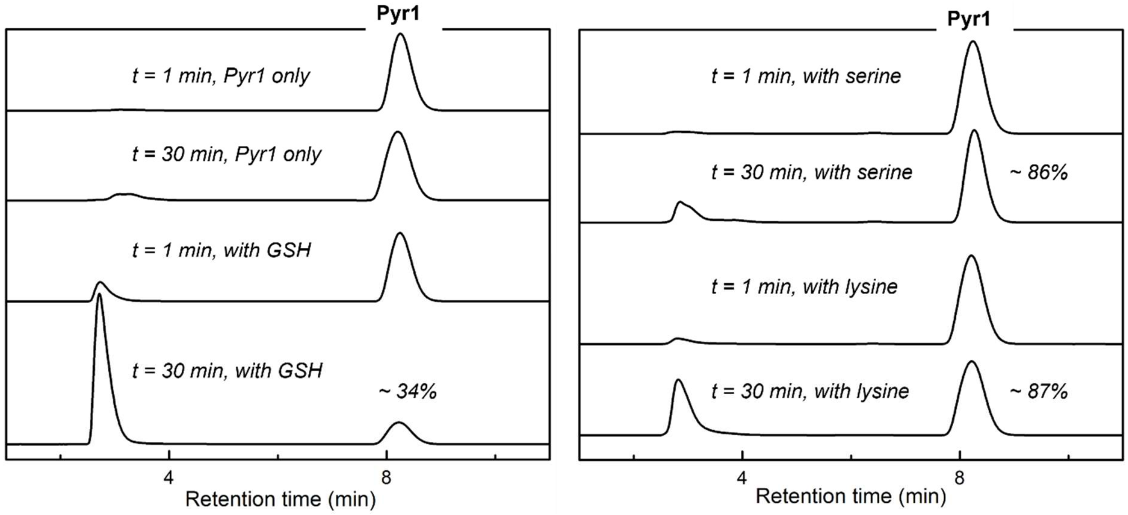


**Figure S5** Bioorthogonality tests between Pyr1 (0.1 mM) and amino acids (GSH, serine, lysine, 1 mM each) determined by HPLC.

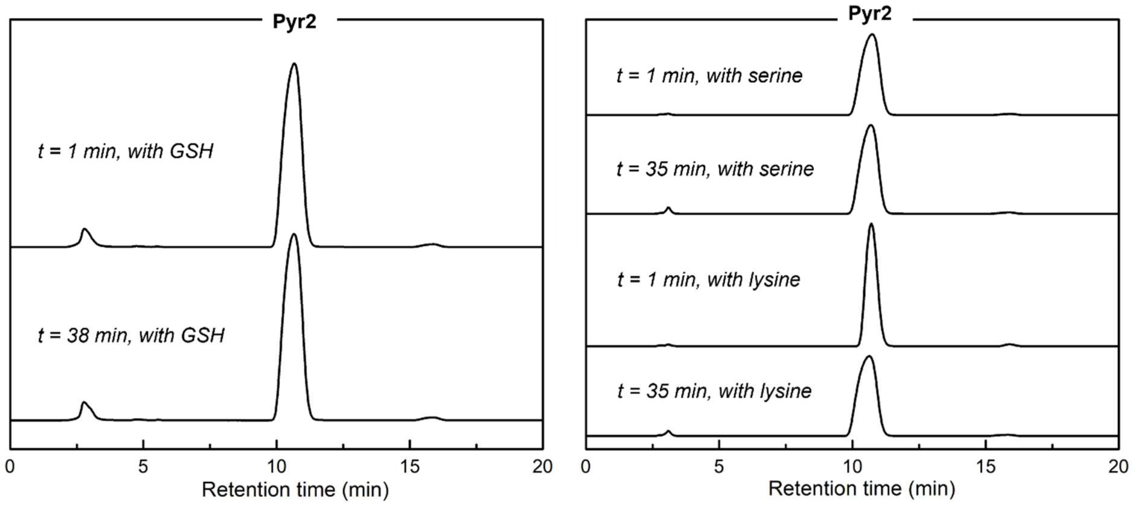


**Figure S6** Bioorthogonality tests between Pyr2 (1 mM) and amino acids (GSH, serine, lysine, 10 mM each) determined by HPLC.

### Protein Labeling Experiments

**Synthesis of lysozyme-BCN**: BCN-NHS was synthesized according to literature[5]. Lysozyme (CAS# 12650-88-3, egg white, 50 mg/mL in deionized water) was diluted with PBS (1X, pH 7.4) to obtain a final concentration of 10 mg/mL. Into the lysozyme solution (250 µL, 10 mg/mL), a solution of BCN-NHS (55 µL, 20 mM in DMSO, 1.0 eq) and additional DMSO (20 µL) were added and then was incubated at room temperature for 1h. The reaction mixture was purified by spin filtration (10 kDa MWCO, 5X 1:5 dilution with PBS (1X, pH 7.4)) to remove unreacted BCN-NHS. The concentration of lysozyme solution was determined using Nanodrop by measuring $A_{280}$ in PBS (1X, pH 7.4). A final concentration of 1.31 mg/mL was obtained by diluting the lysozyme solution with PBS (1X, pH 7.4), ready for protein labeling experiments. Aliquots of lysozyme solution were kept at -20 °C for future experiments. The presence of lysozyme-BCN was confirmed by MALDI-TOF.

**Concentration-dependent protein labeling experiments**.

Reaction mixtures consisting of protein solution (unmodified lysozyme or lysozyme-BCN, 5 µL, 1.31 mg/mL) and sulforhodamine B-pyrone (0.26 µL in DMSO, final concentration, 400, 200, 100, and 50 µM) were incubated for 4h at room temperature. The reactions were quenched with sample loading buffer (1.75 µL, 4x Laemmli sample buffer, Cat. #1610747, BIO-RAD). Each solution was loaded into a 10 well 10% SDS-PAGE gel (Cat. #456-1033, BIO-RAD) and run at 120 V for 1h at room temperature. In-gel fluorescent imaging was obtained with a Typhoon 9400 at 580 nm (580 BP 30) with a photomultiplier tube (PMT) setting of 400V.

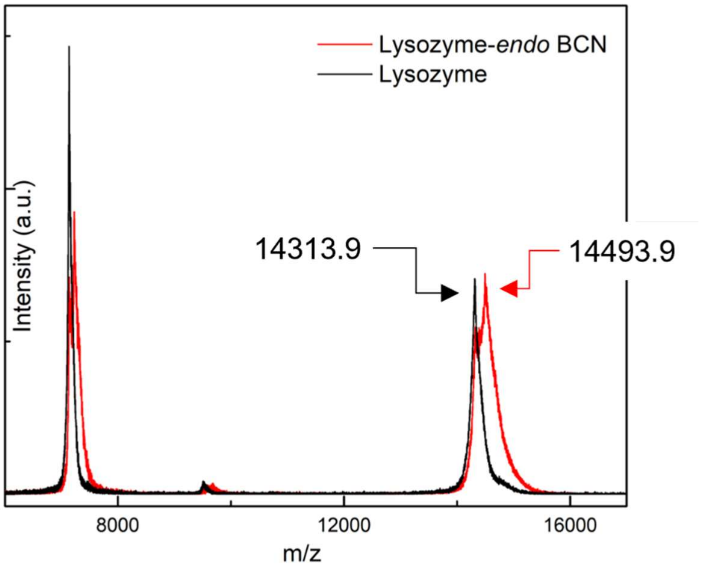


**Figure S7** MALDI-TOF spectrum of BCN functionalized lysozyme and unfunctionalized lysozyme.

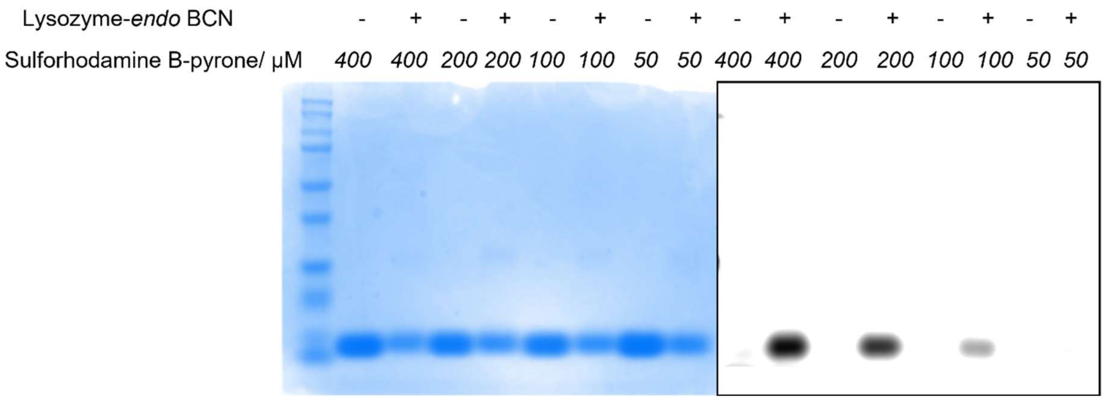


**Figure S8** In-gel fluorescent results for concentration dependent labeling of lysozyme using Sulforhodamine B-pyrone conjugate. Left, Coomassie staining; right, fluorescence at 580 nm.

**Fluorescent Assay of $H_2S$ release**.

Reaction mixtures (200 µL each, 1%DMSO in PBS (1X, pH 7.4)) consisting of pyranthiones (Pyr1, Pyr2, Pyr3, Pyr4, final concentration 100 µM), BCN (final concentration, 100 µM), fluorescent sensor NDI (final concentration, 10 µM), with or without carbonic anhydrase (Isozyme II from bovine erythrocytes, C2522, sigma, final concentration, 50 µg/mL) in were prepared in separate wells of a 96-well plate at room temperature. For PyrS2, control experiments with amino acids were conducted by replacing BCN by serine, lysine or GSH with a final concentration 100 µM in the presence of carbonic anhydrase. Positive control was done by using $Na_2S$ (final concentration, 100 µM) as the $H_2S$ donor. The reaction was monitored by measuring the fluorescence emission generated by released NDI piperazine intermediate using plate reader (excitation filter: 340/30 nm, emission filter: 528/20 nm).

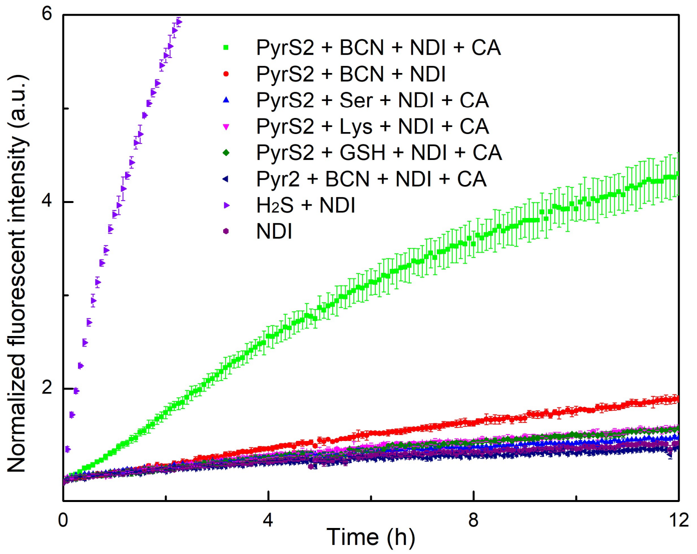


**Figure S9** Bioorthogonal release of $H_2S$ through the reaction of PyrS2 with BCN as determined by fluorescent assay using NDI.

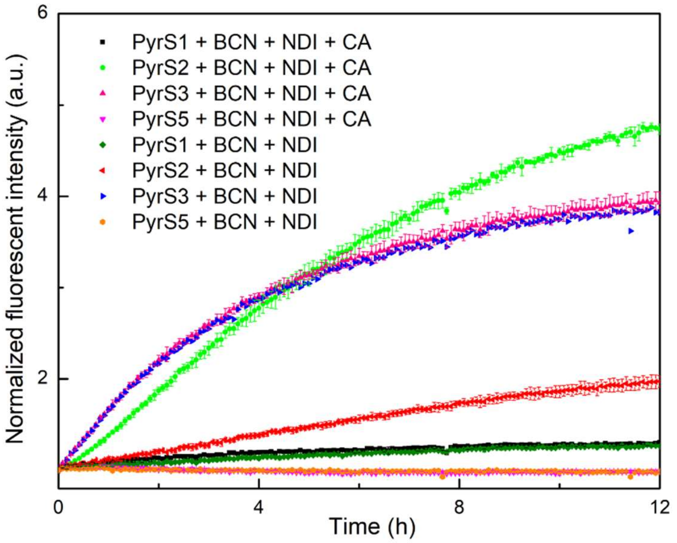


**Figure S10** $H_2S$ release evaluation of pyranthiones with different reaction rates with BCN as determined by fluorescent assay using NDI.

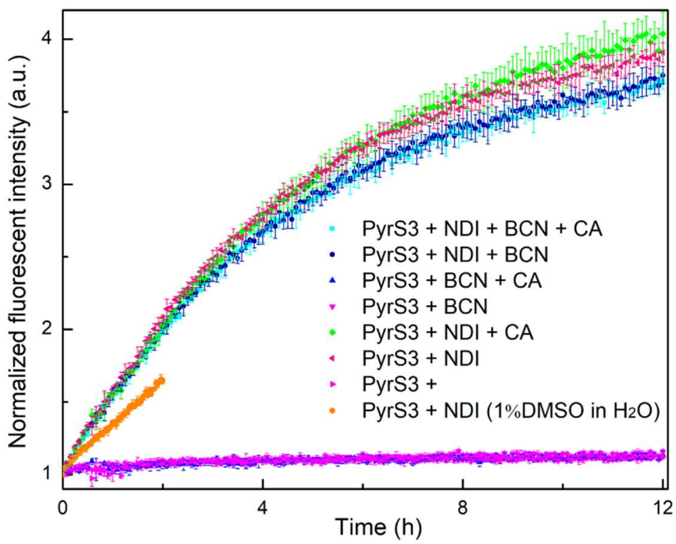


**Figure S11** $H_2S$ release evaluation of PyrS3 as determined by fluorescent assay using NDI showed the possible degradation of PyrS3 in DMSO.

**Quantification of $H_2S$ release by methylene blue (MB) assay.**

**$H_2S$ calibration curve generation**. PBS (1X, pH 7.4, ~20 mL) was degassed for 40 minutes using nitrogen before use. 8.0 mL of MB cocktail [1.6 mL $Zn(OAc)_2$ (1% w/v, 120 mg $Zn(OAc)_2$ dihydrate in 10 mL deionized water), 3.2 mL $FeCl_3$ (30 mM in 1.2 M HCl), and 3.2 mL *N*,*N*-dimethyl-p-phenylene diamine (20 mM in 7.2 M HCl) was mixed with 1.6 mL DMSO or acetonitrile and then was aliquoted each with 500 μL. To each aliquot, NaHS stock solutions (100-fold to final concentrations in PBS) was added to prepare the final NaHS concentration of 1.56, 2.34, 3.13, 4.69, 6.25, 9.38, 12.5, 18.8, 25, 37.5, 50, 75, 100, 150, and 200 μM. Then these aliquot mixtures were incubated for 1h at 37 °C in the dark. After that, 200 μL from each aliquot mixture was transferred to separate wells of a 96-well plate and the absorbance was determined at 670 nm by plate reader. The $H_2S$ calibration curve was generated by linear fitting of the absorbance value versus concentration of NaHS.

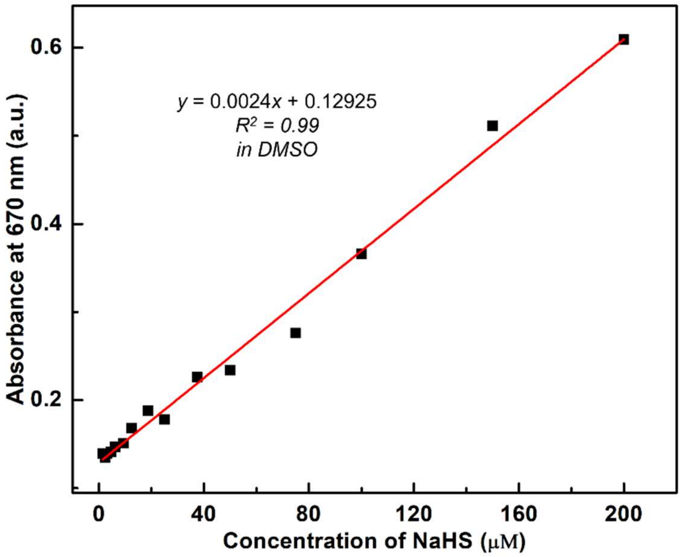


**Figure S12** Calibration curve between $H_2S$ concentration and absorbance at 670 nm determined by Methylene Blue assay in DMSO.

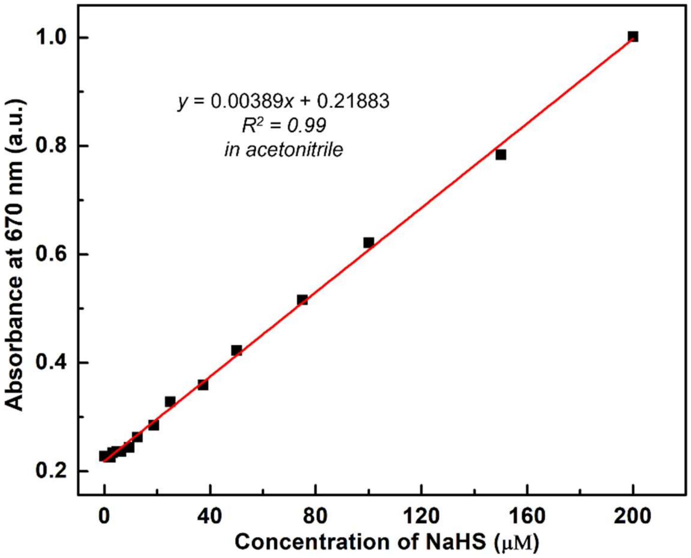


**Figure S13** Calibration curve between $H_2S$ concentration and absorbance at 670 nm determined by Methylene Blue assay in acetonitrile.

**Time dependent $H_2S$ release from click reaction between pyranthiones and BCN by MB assay**. 20 mL PBS (1X, pH 7.4) was degassed for 40 minutes using nitrogen before use. MB cocktail was mixed with DMSO or acetonitrile with a volume ratio of 3 to 1 before use. Into 4 mL PBS containing carbonic anhydrase (final concentration 50 μg/mL) in 20 mL plastic vial, pyranthiones (10 mM in DMSO, 40 μL) and BCN (100 mM in DMSO, 40 μL) were sequentially added to make final concentrations of 100 μM and 1 mM for pyranthiones and BCN, respectively. Aliquots (120 μL, 3 times) of reaction mixture were taken out with certain time intervals, diluted with MB cocktail (120 μL), and incubated for 1h at 37 $^oC$ in the dark. After that, 200 μL of reaction mixture was transferred into separate wells of a 96-well plate and the absorbance was determined at 670 nm by plate reader. The concentration of generated $H_2S$ was determined using the $H_2S$ calibration curve. Note, we replaced DMSO by acetonitrile for $H_2S$ calibration curve determination, the MB cocktail, and the stock solution preparation of PyrS3 since significant degradation of PyrS3 in DMSO was observed. And we used a final concentration of 200 μM for PyrS4 due to its relatively slower kinetics.

**HeLa cell culture.** HeLa cells were maintained in high-glucose Dulbecco's Modified Eagle Medium (DMEM) supplemented with 10% FBS and 1% Gibco$^{TM}$ Antibiotic-Antimycotic at 37 °C in air supplemented with 5% $CO_2$. Cells were passaged on alternating days when they reached approximately 90% confluency.

**HeLa cell imaging.** Cells were seeded at 100,000 cells per 35 mm glass-bottom culture dish (MatTek) and grown for 48 h to 60–80% confluency. To begin labeling, the cells were washed once with 37 $^oC$ PBS (1 mL) and then incubated with a mixed solution containing Mito-NDI (10 µM, 100 µL from 100 µM stock solution in DMSO) and $PPh_3$-PyrS2 (40 µM, 10 µL from 4 mM stock solution in DMSO) in DMEM (890 µL) at 37 $^oC$ for 30 minutes. After that, the media was aspirated and the cells were incubated with BCN (200 µM, 10 µL from 20 mM stock solution in DMSO) in DMEM (990 µL) at 37 $^oC$ for 2.5h. After removal of the medium and washing with 37 $^oC$ PBS (1 mL), the cells were stained with MitoTracker Red (200 nM, 1 µL from 200 µM stock solution in DMSO) in DMEM (1 mL) at 37 $^oC$ for 21 minutes. The medium was removed, the cells were washed with 37 $^oC$ PBS (1 mL) and then stained by Hoechst (1 µg/mL, 1 µL from 1 mg/mL stock in water) in DMEM (1 mL) at 37 $^oC$ for 15 minutes. Finally, the medium was removed and the cells were washed with 37 $^oC$ PBS (1 mL) followed by adding 37 $^oC$ FluoroBrite DMEM (2 mL). The cells were then imaged by confocal microscopy. For control experiments, BCN was replaced by DMSO vehicle or $PPh_3$-PyrS2 was replaced by the pyrone counterpart, $PPh_3$-Pyr2.

**NMR spectra**

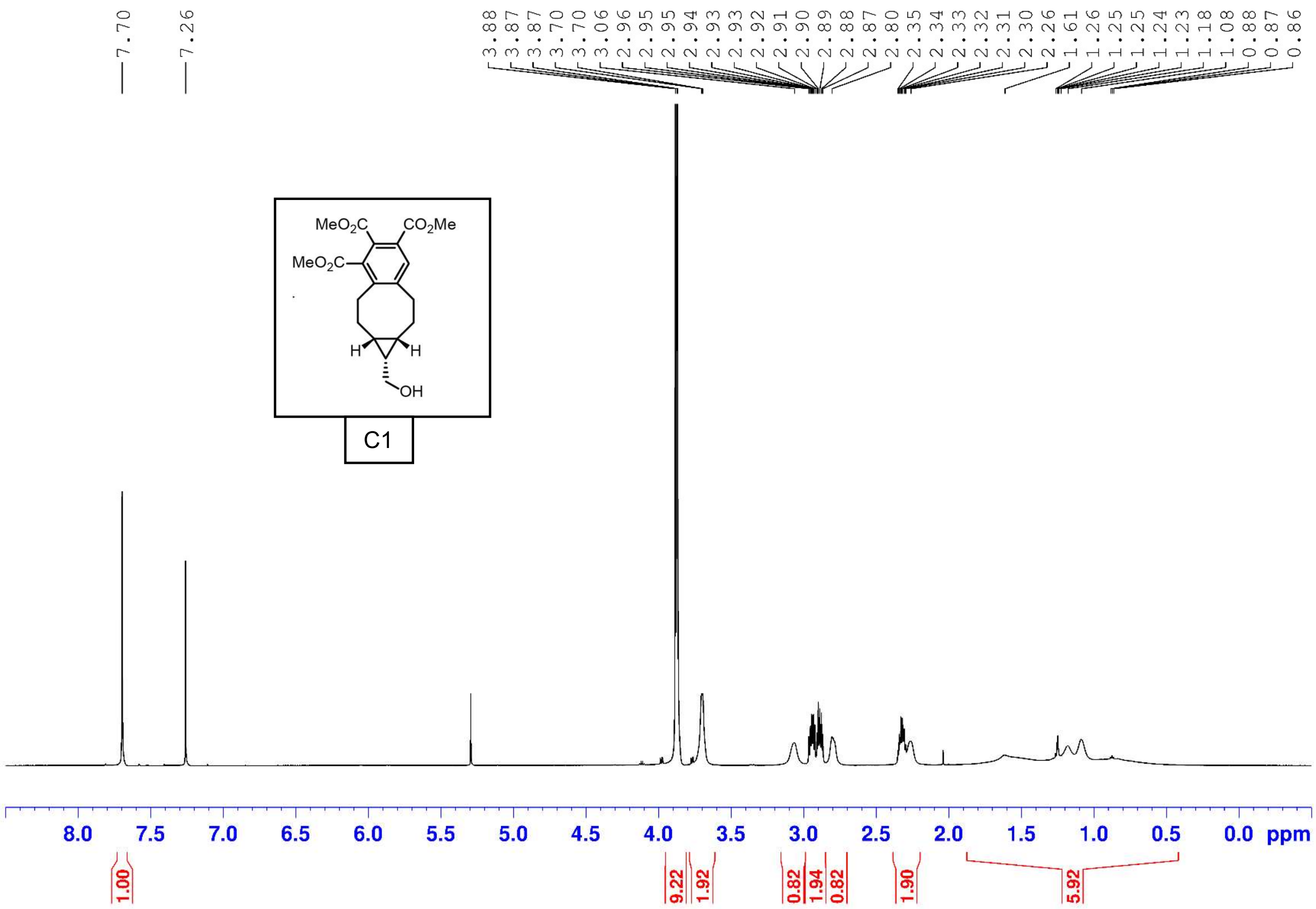

7.70
7.26
3.88 3.87 3.87 3.70 3.70 3.06 2.96 2.95 2.95 2.94 2.93 2.93 2.92 2.91 2.90 2.89 2.88 2.87 2.80 2.35 2.34 2.33 2.32 2.31 2.30 2.26 1.61 1.26 1.25 1.25 1.24 1.23 1.18 1.08 0.88 0.87 0.86
MeO2C
CO2Me
MeO2C
H
H
OH
C1
8.0 7.5 7.0 6.5 6.0 5.5 5.0 4.5 4.0 3.5 3.0 2.5 2.0 1.5 1.0 0.5 0.0 ppm
1.00
9.22
1.92
0.82
1.94
0.82
1.90
5.92

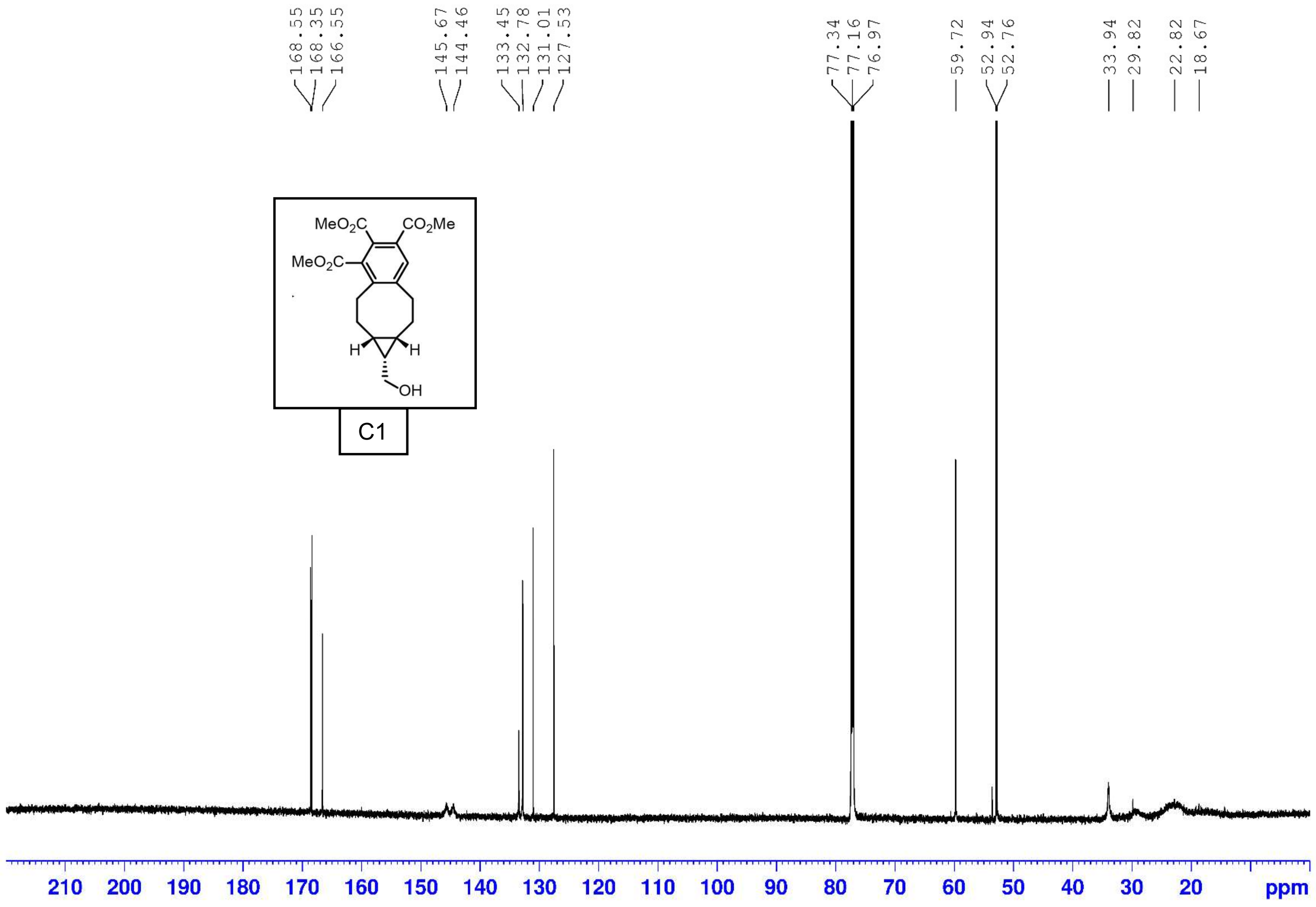
168.55
168.35
166.55
145.67
144.46
133.45
132.78
131.01
127.53
77.34
77.16
76.97
59.72
52.94
52.76
33.94
29.82
22.82
18.67
MeO2C
CO2Me
MeO2C
H
H
OH
C1
210 200 190 180 170 160 150 140 130 120 110 100 90 80 70 60 50 40 30 20 ppm

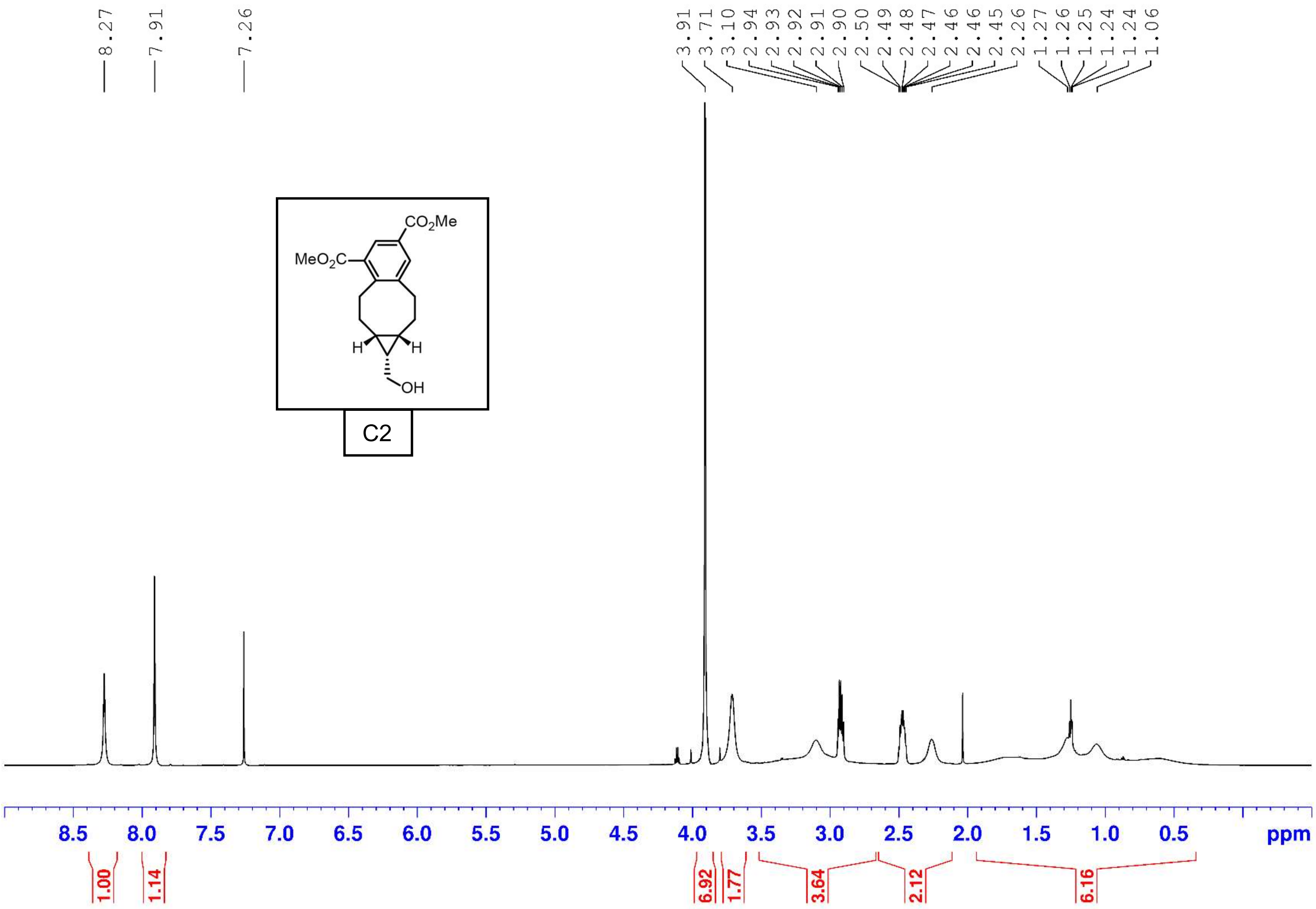
8.27
7.91
7.26
3.91
3.71
3.10
2.94
2.93
2.92
2.91
2.90
2.50
2.49
2.48
2.47
2.46
2.46
2.45
2.26
1.27
1.26
1.25
1.24
1.24
1.06
$CO_2Me$
$MeO_2C$
H
H
OH
C2
8.5
8.0
7.5
7.0
6.5
6.0
5.5
5.0
4.5
4.0
3.5
3.0
2.5
2.0
1.5
1.0
0.5
ppm
1.00
1.14
6.92
1.77
3.64
2.12
6.16

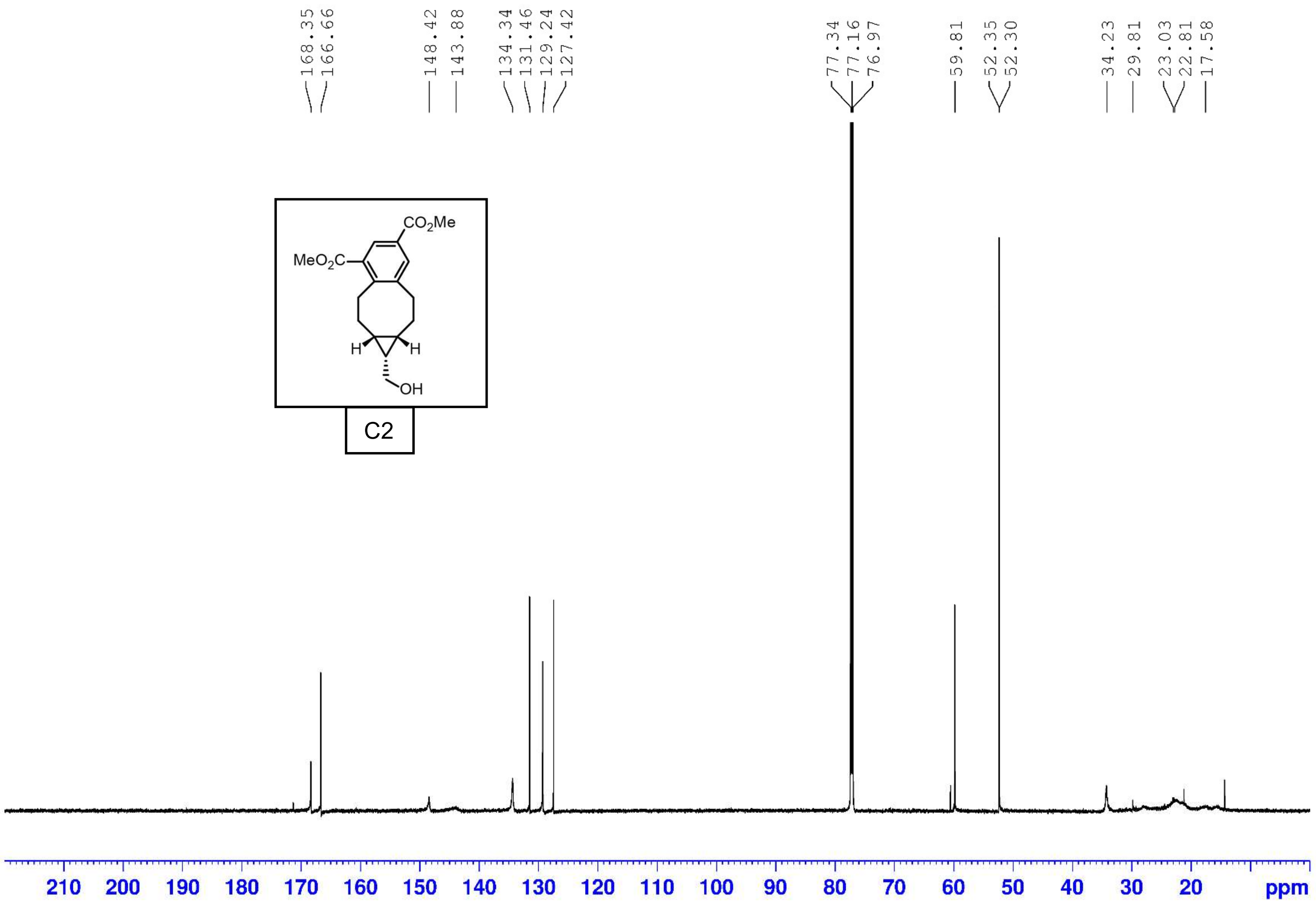
168.35
166.66
148.42
143.88
134.34
131.46
129.24
127.42
77.34
77.16
76.97
59.81
52.35
52.30
34.23
29.81
23.03
22.81
17.58
$CO_2Me$
$MeO_2C$
H
H
OH
C2
210 200 190 180 170 160 150 140 130 120 110 100 90 80 70 60 50 40 30 20 ppm

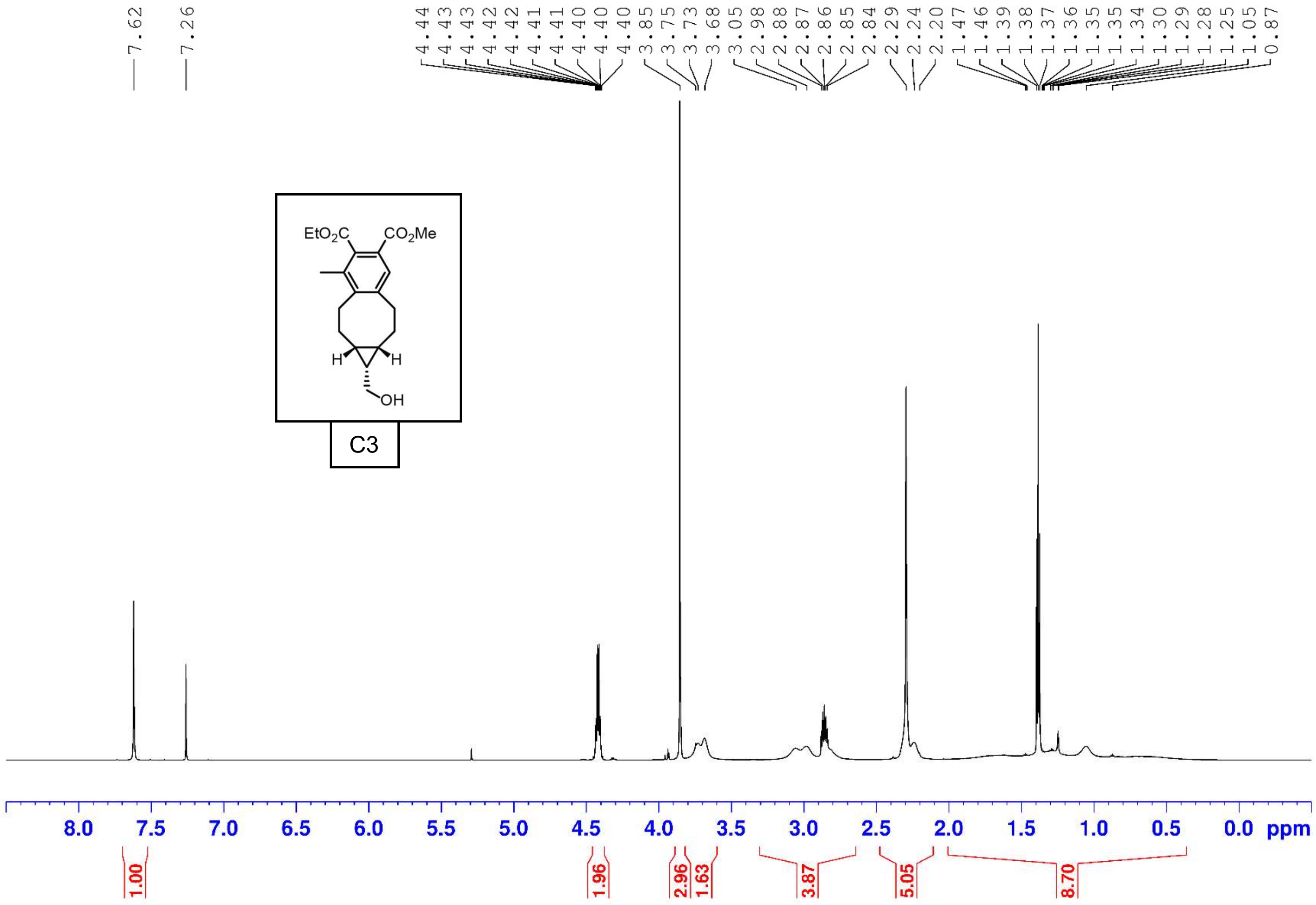
7.62
7.26
4.44 4.43 4.43 4.42 4.42 4.41 4.41 4.40 4.40 4.40 3.85 3.75 3.73 3.68 3.05 2.98 2.88 2.87 2.86 2.85 2.84 2.29 2.24 2.20 1.47 1.46 1.39 1.38 1.37 1.36 1.35 1.35 1.34 1.30 1.29 1.28 1.25 1.05 0.87
EtO2C
CO2Me
H
H
OH
C3
8.0 7.5 7.0 6.5 6.0 5.5 5.0 4.5 4.0 3.5 3.0 2.5 2.0 1.5 1.0 0.5 0.0 ppm
1.00
1.96
2.96
1.63
3.87
5.05
8.70

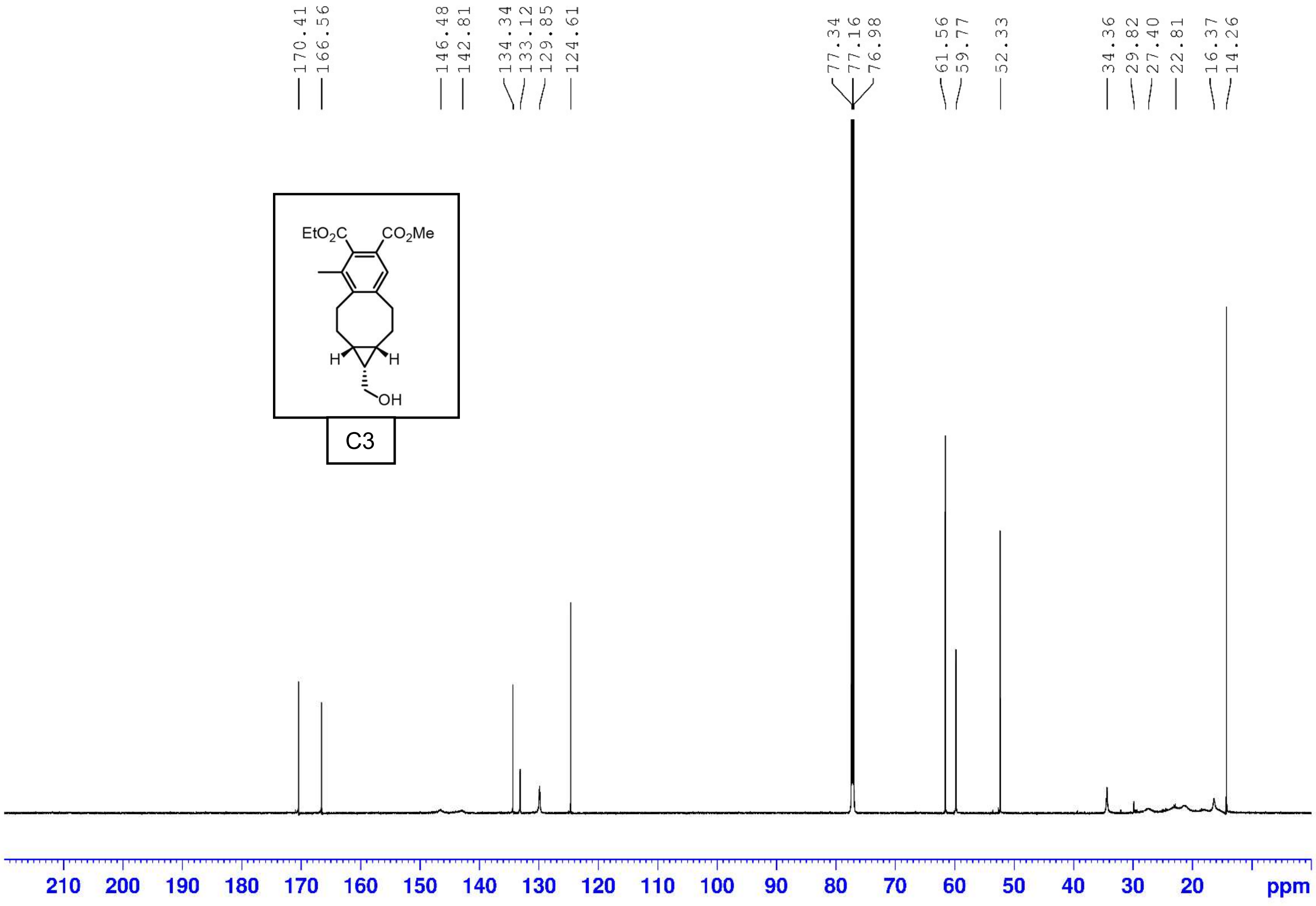
170.41
166.56
146.48
142.81
134.34
133.12
129.85
124.61
77.34
77.16
76.98
61.56
59.77
52.33
34.36
29.82
27.40
22.81
16.37
14.26
EtO2C
CO2Me
H
H
OH
C3
210 200 190 180 170 160 150 140 130 120 110 100 90 80 70 60 50 40 30 20 ppm

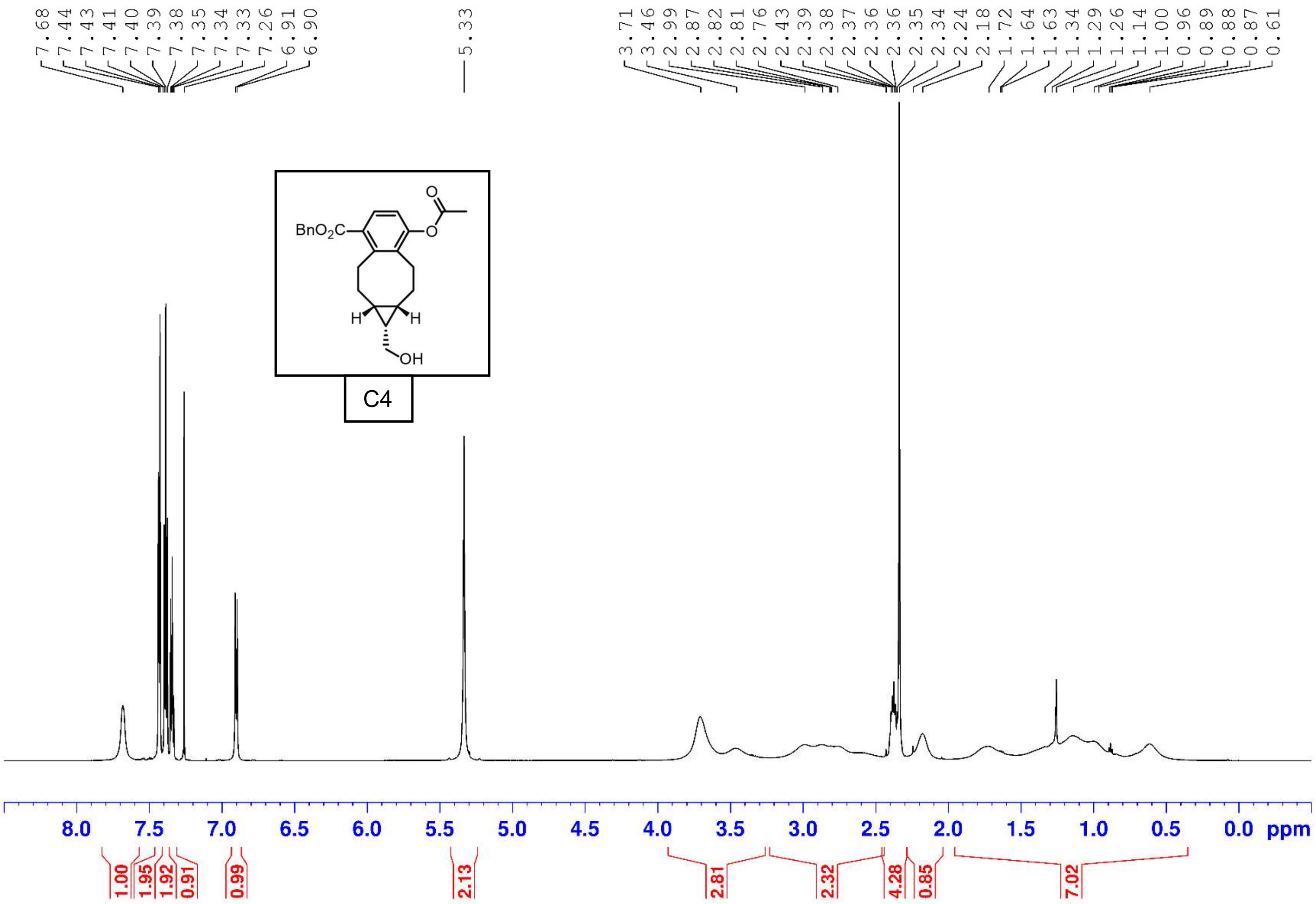
7.68 7.44 7.43 7.41 7.40 7.39 7.38 7.35 7.34 7.33 7.26 6.91 6.90
5.33
3.71 3.46 2.99 2.87 2.82 2.81 2.76 2.43 2.39 2.38 2.37 2.36 2.36 2.35 2.34 2.24 2.18 1.72 1.64 1.63 1.34 1.29 1.26 1.14 1.00 0.96 0.89 0.88 0.87 0.61
BnO2C
O
O
H
H
OH
C4
8.0 7.5 7.0 6.5 6.0 5.5 5.0 4.5 4.0 3.5 3.0 2.5 2.0 1.5 1.0 0.5 0.0 ppm
1.00 1.95 1.92 0.91 0.99 2.13 2.81 2.32 4.28 0.85 7.02

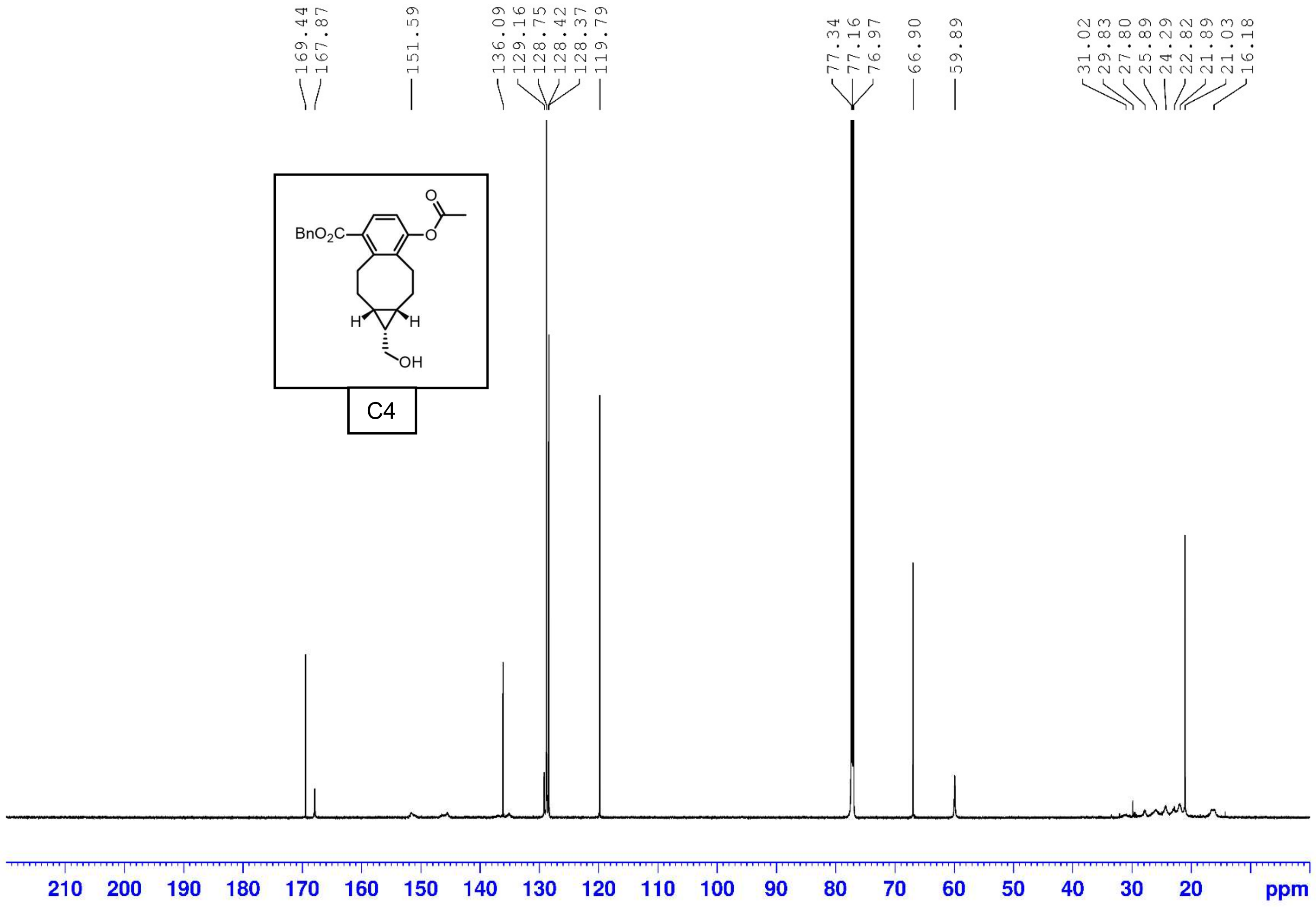
169.44
167.87
151.59
136.09
129.16
128.75
128.42
128.37
119.79
77.34
77.16
76.97
66.90
59.89
31.02
29.83
27.80
25.89
24.29
22.82
21.89
21.03
16.18
BnO2C
O
O
H
H
OH
C4
210
200
190
180
170
160
150
140
130
120
110
100
90
80
70
60
50
40
30
20
ppm

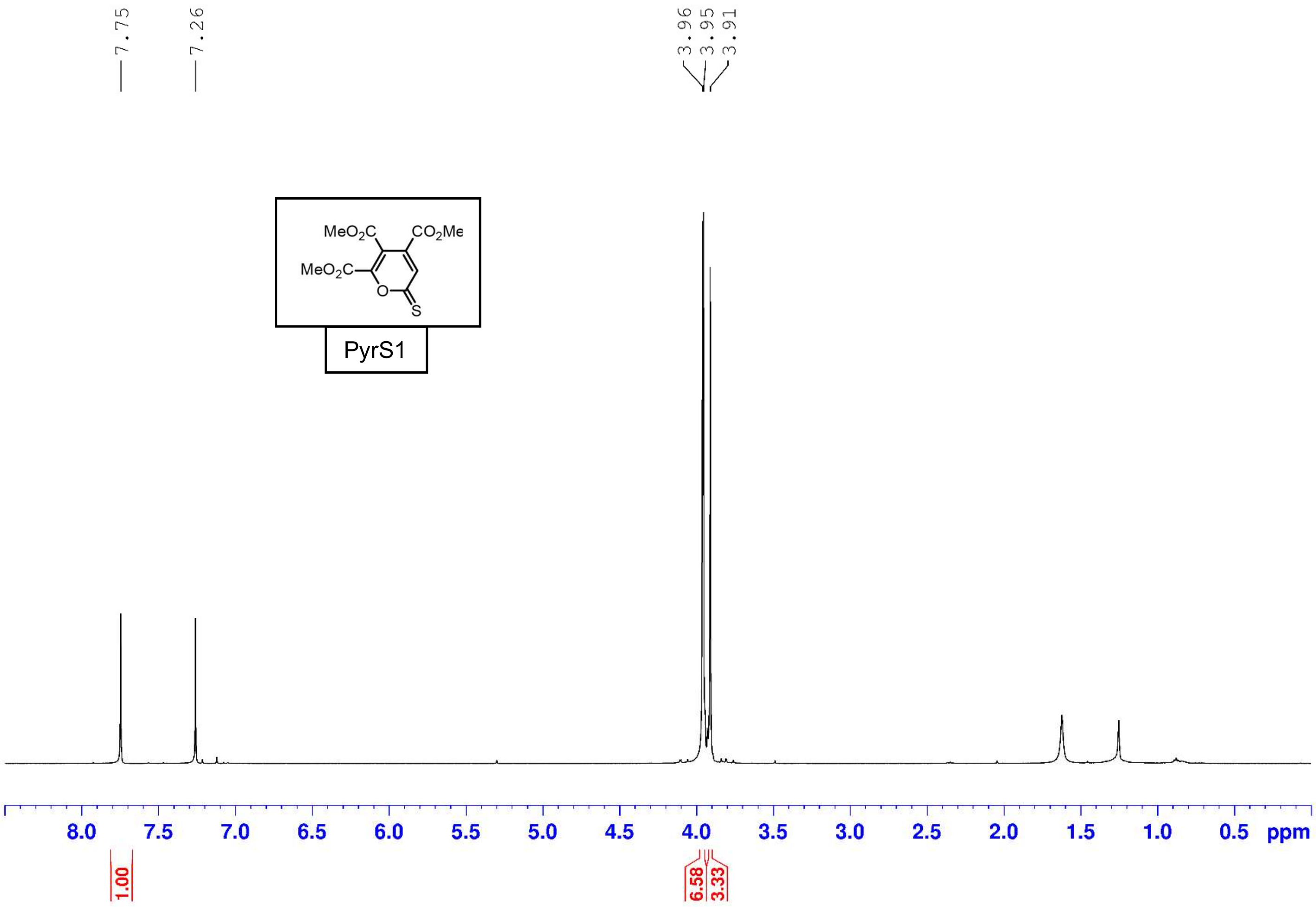

7.75
7.26
3.96
3.95
3.91
MeO2C
CO2Me
MeO2C
O
S
PyrS1
8.0
7.5
7.0
6.5
6.0
5.5
5.0
4.5
4.0
3.5
3.0
2.5
2.0
1.5
1.0
0.5
ppm
1.00
6.58
3.33

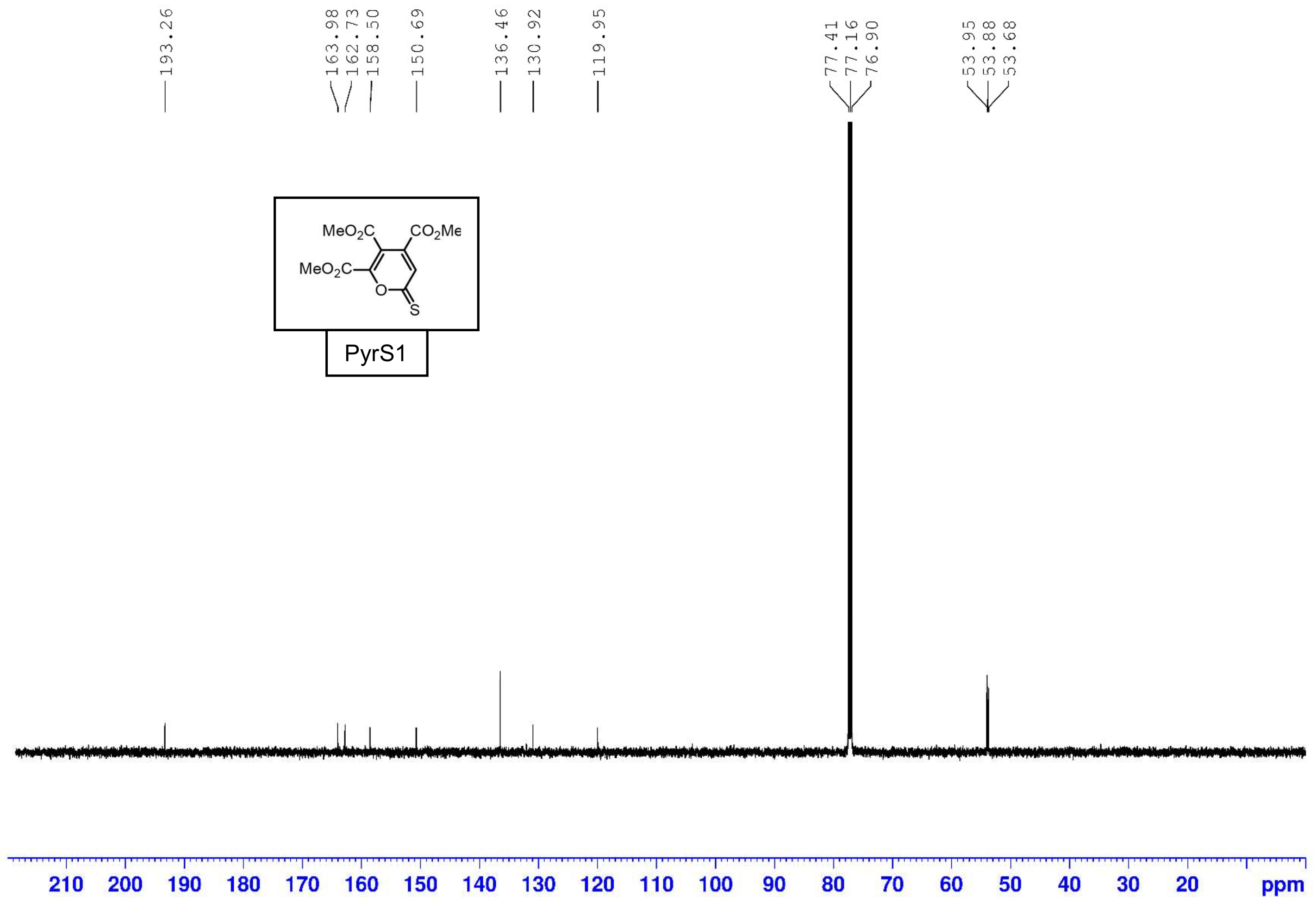
193.26
163.98
162.73
158.50
150.69
136.46
130.92
119.95
77.41
77.16
76.90
53.95
53.88
53.68
MeO2C
CO2Me
MeO2C
O
S
PyrS1
210 200 190 180 170 160 150 140 130 120 110 100 90 80 70 60 50 40 30 20
ppm

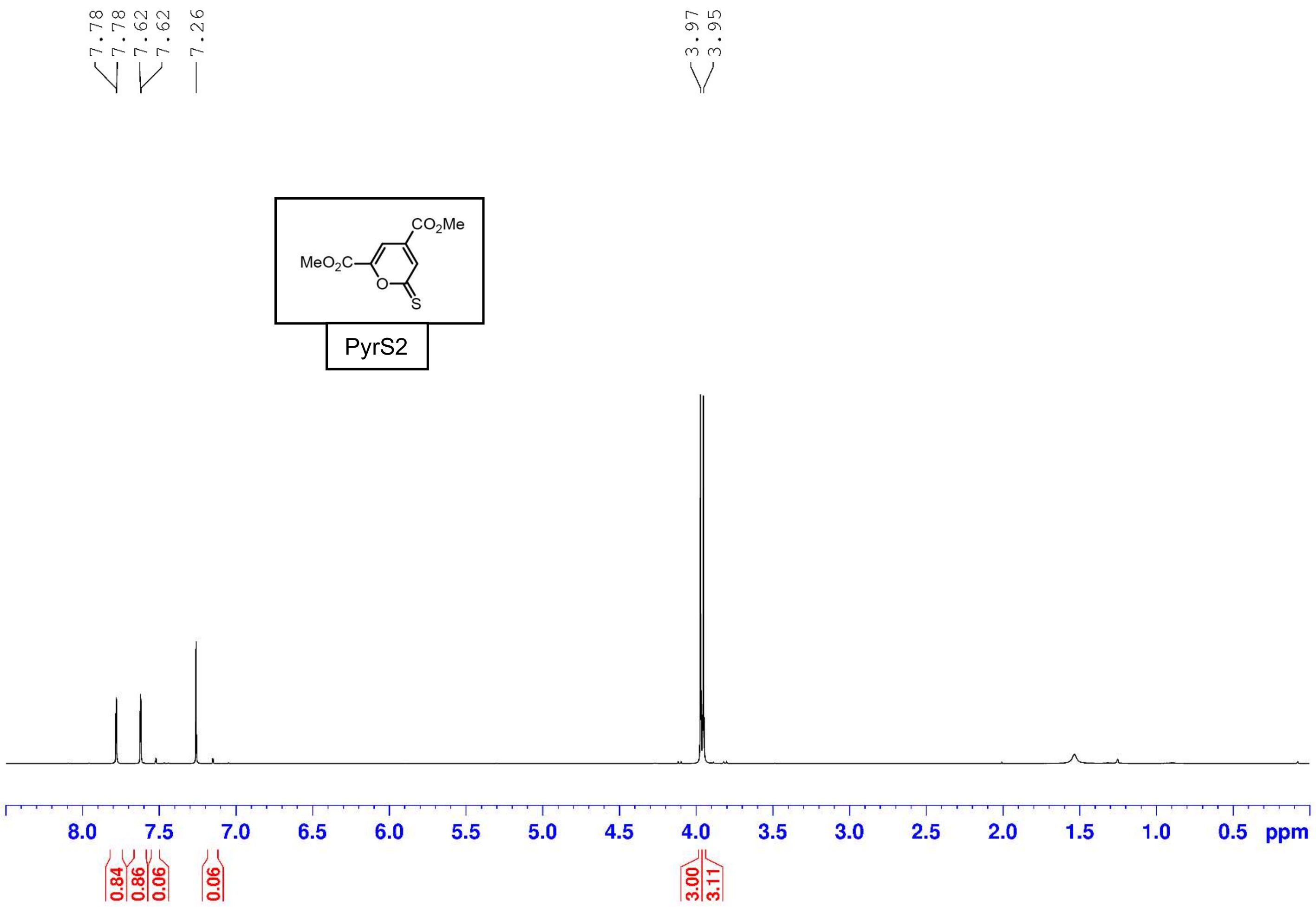
7.78
7.78
7.62
7.62
7.26
3.97
3.95
CO2Me
MeO2C
O
S
PyrS2
8.0
7.5
7.0
6.5
6.0
5.5
5.0
4.5
4.0
3.5
3.0
2.5
2.0
1.5
1.0
0.5
ppm
0.84
0.86
0.06
0.06
3.00
3.11

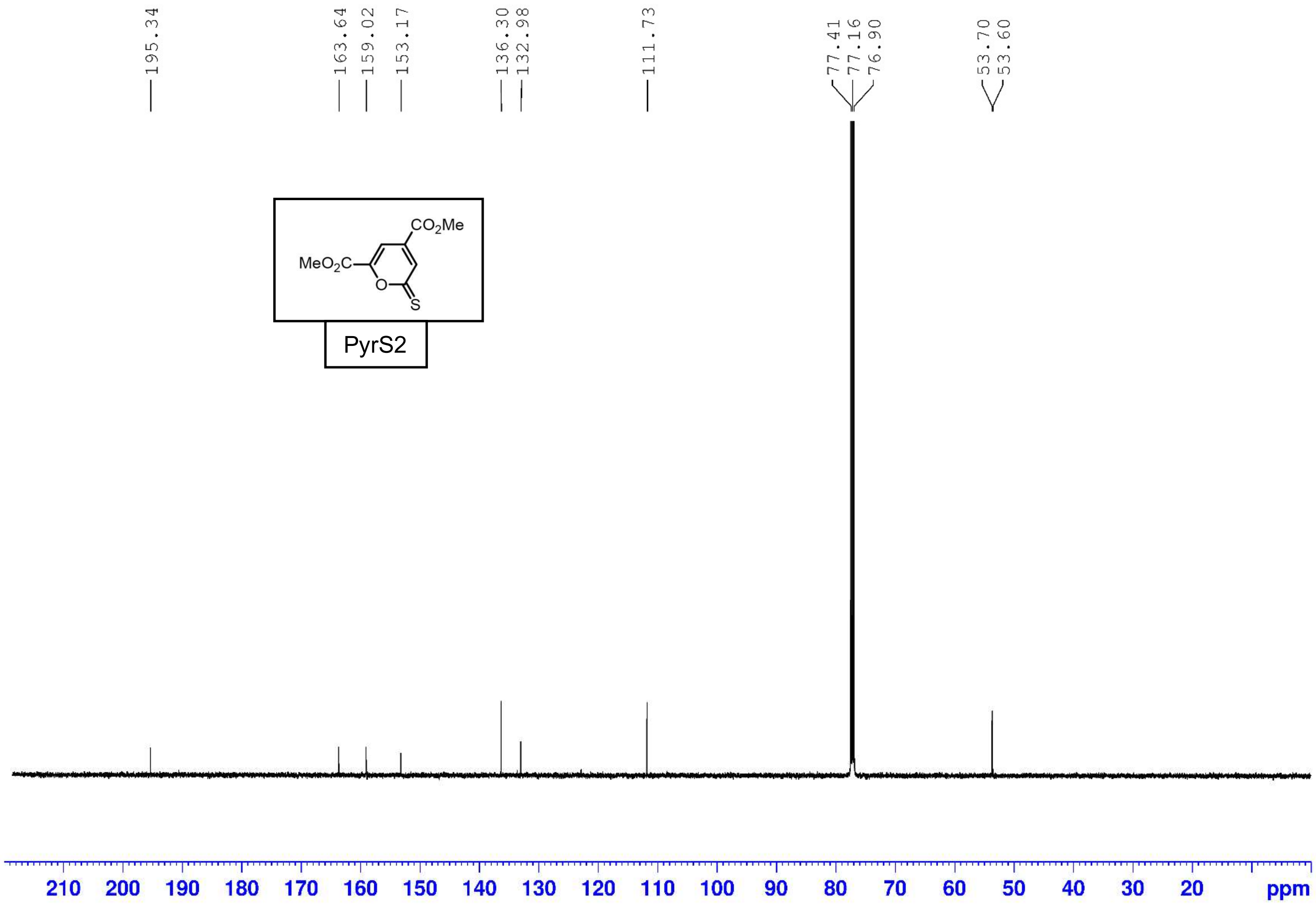
195.34
163.64
159.02
153.17
136.30
132.98
111.73
77.41
77.16
76.90
53.70
53.60
CO2Me
MeO2C
O
S
PyrS2
210
200
190
180
170
160
150
140
130
120
110
100
90
80
70
60
50
40
30
20
ppm

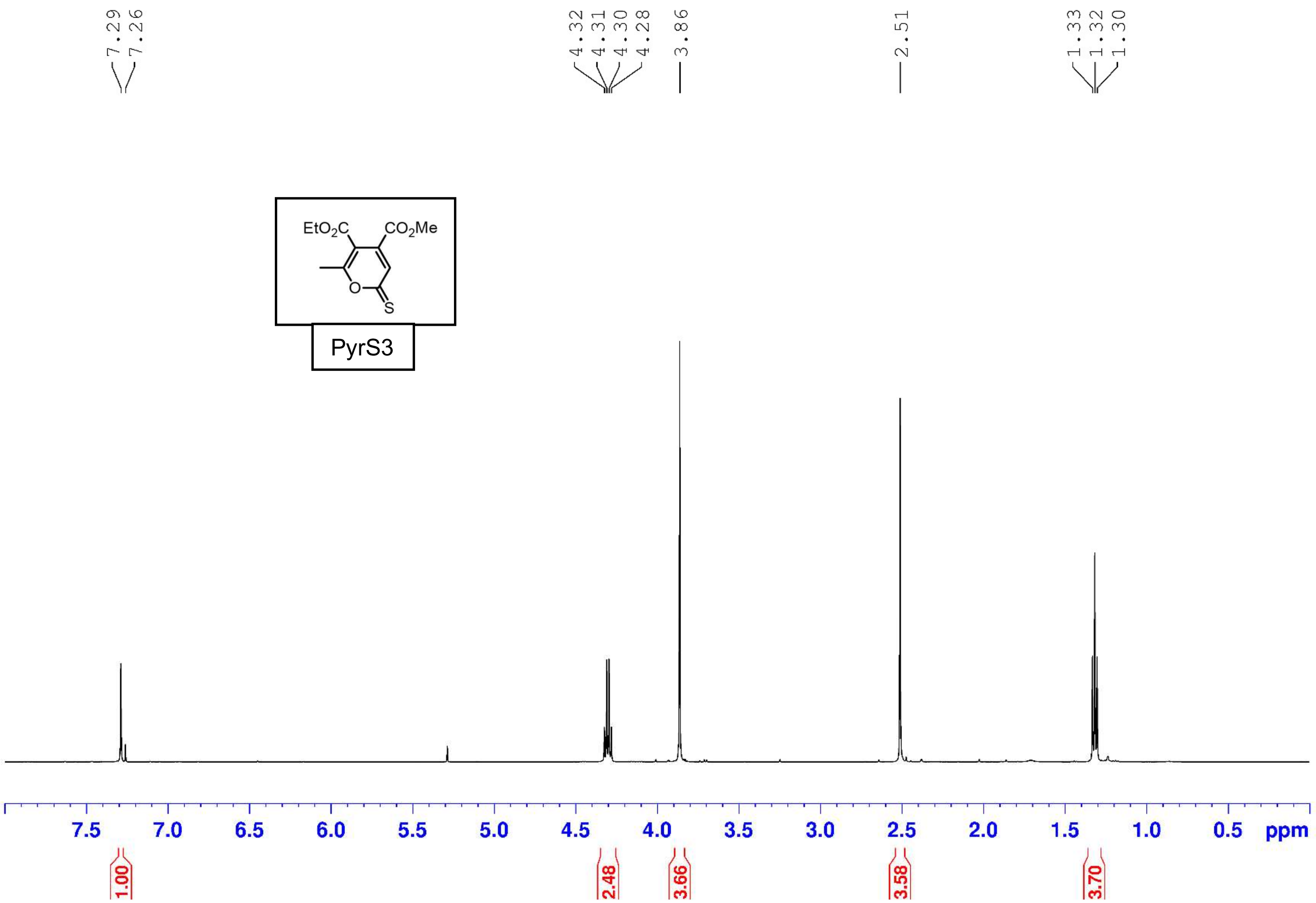

7.29
7.26
4.32
4.31
4.30
4.28
3.86
2.51
1.33
1.32
1.30
EtO2C
CO2Me
O
S
PyrS3
7.5
7.0
6.5
6.0
5.5
5.0
4.5
4.0
3.5
3.0
2.5
2.0
1.5
1.0
0.5
ppm
1.00
2.48
3.66
3.58
3.70

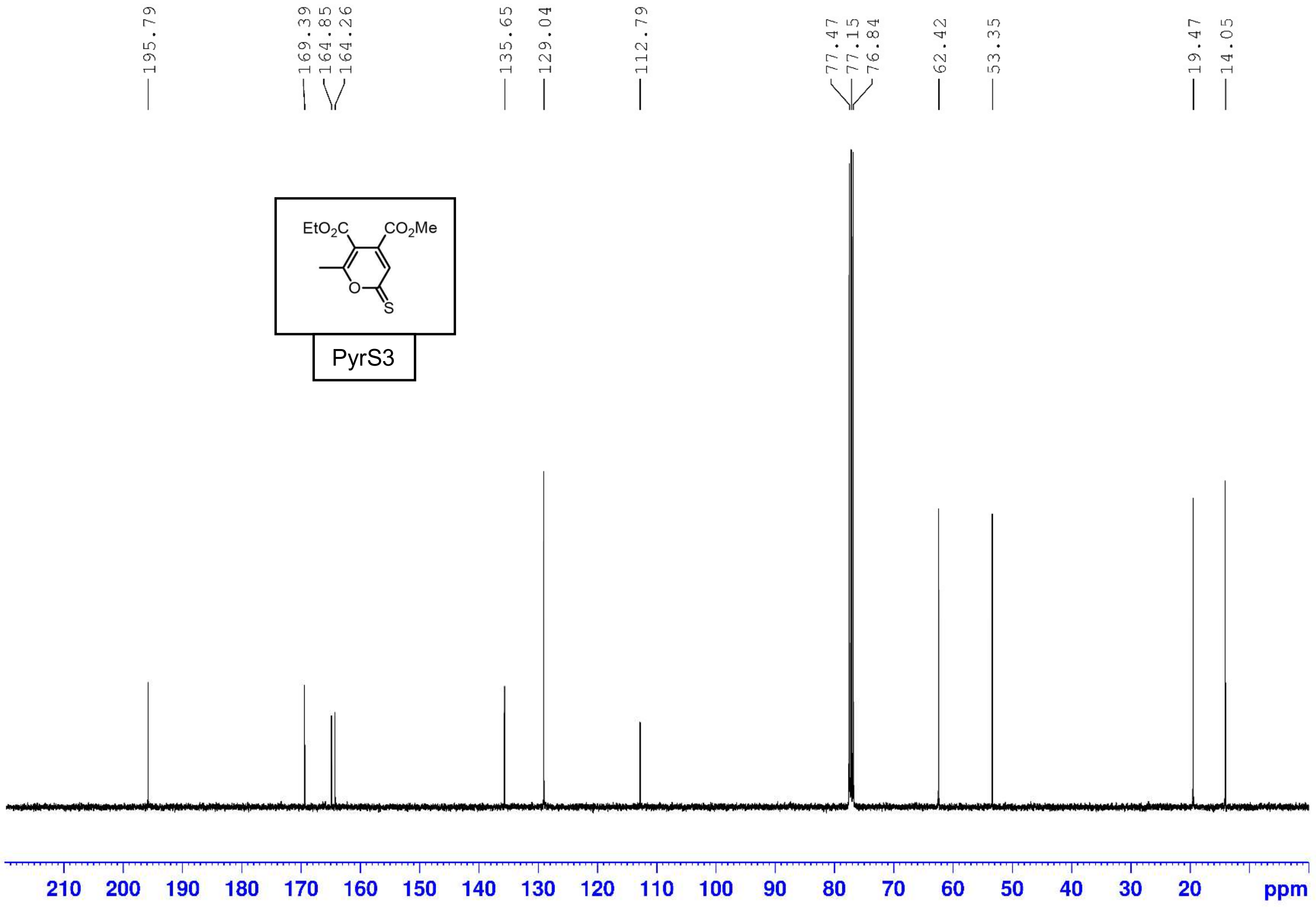
195.79
169.39
164.85
164.26
135.65
129.04
112.79
77.47
77.15
76.84
62.42
53.35
19.47
14.05
EtO2C
CO2Me
O
S
PyrS3
210 200 190 180 170 160 150 140 130 120 110 100 90 80 70 60 50 40 30 20
ppm

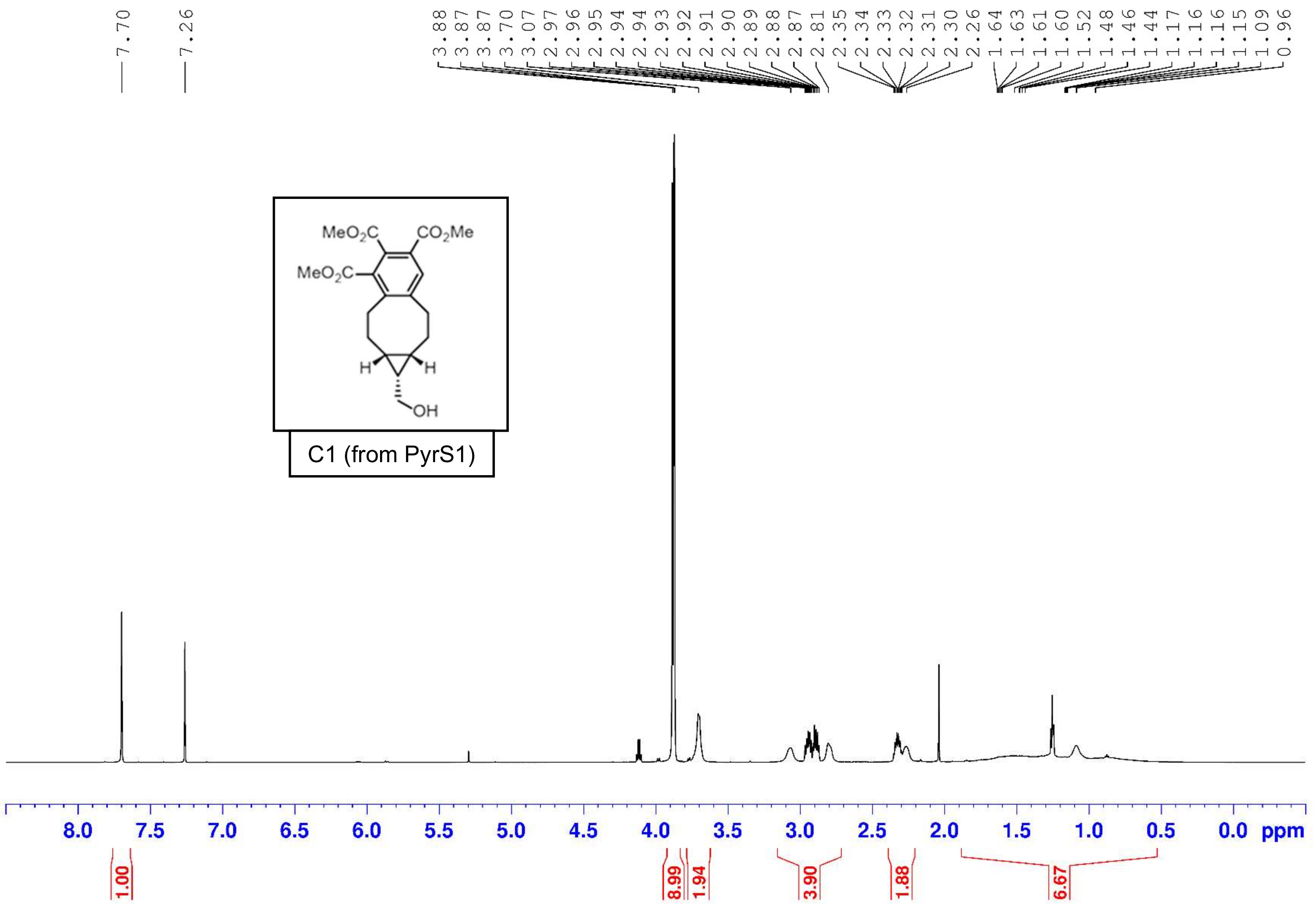

7.70
7.26
3.88 3.87 3.87 3.70 3.07 2.97 2.96 2.95 2.94 2.94 2.93 2.92 2.91 2.90 2.89 2.88 2.87 2.81 2.35 2.34 2.33 2.32 2.31 2.30 2.26 1.64 1.63 1.61 1.60 1.52 1.48 1.46 1.44 1.17 1.16 1.16 1.15 1.09 0.96
MeO2C
CO2Me
MeO2C
H
H
OH
C1 (from PyrS1)
8.0 7.5 7.0 6.5 6.0 5.5 5.0 4.5 4.0 3.5 3.0 2.5 2.0 1.5 1.0 0.5 0.0 ppm
1.00
8.99
1.94
3.90
1.88
6.67

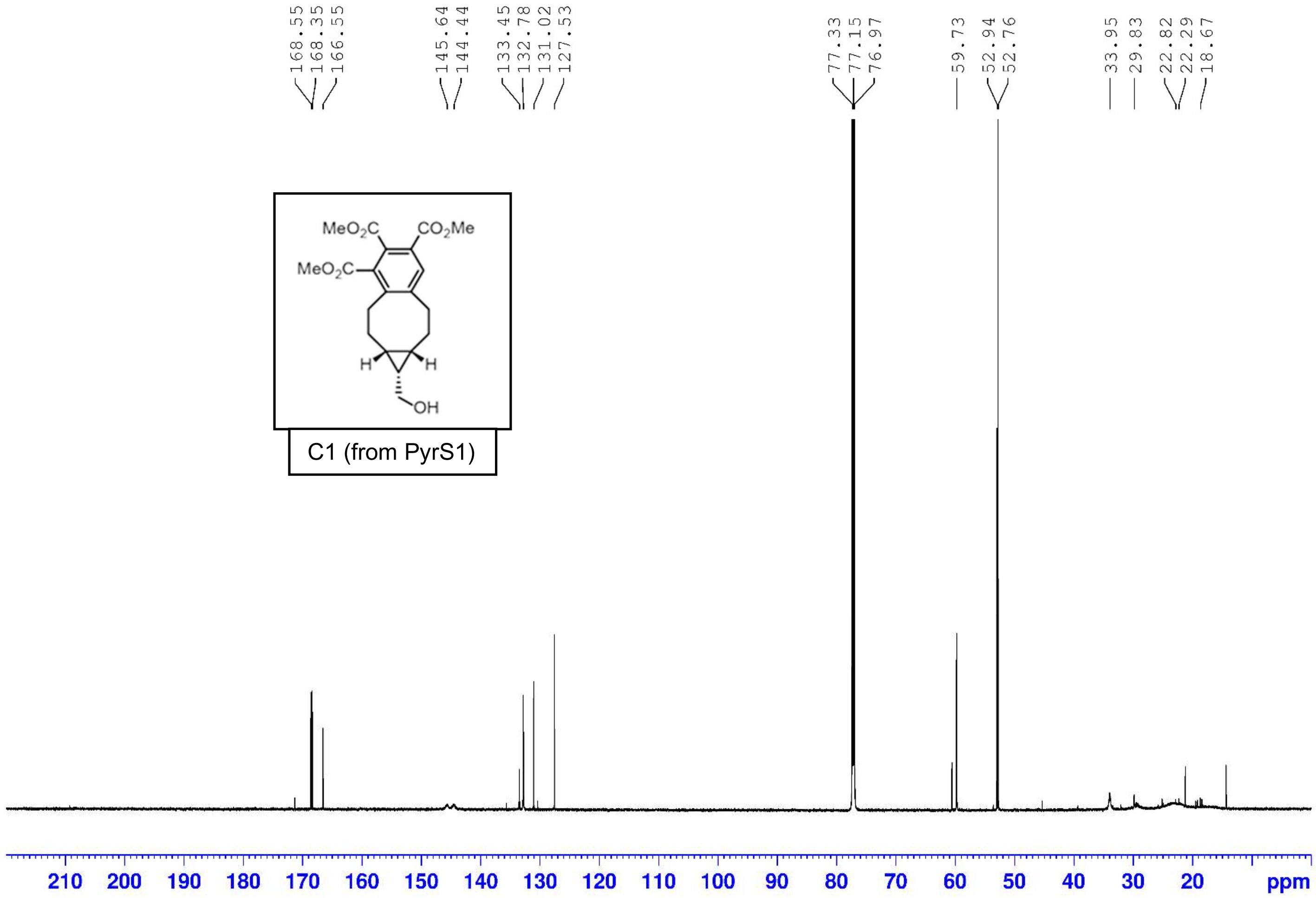
168.55
168.35
166.55
145.64
144.44
133.45
132.78
131.02
127.53
77.33
77.15
76.97
59.73
52.94
52.76
33.95
29.83
22.82
22.29
18.67
MeO2C
CO2Me
MeO2C
H
H
OH
C1 (from PyrS1)
210 200 190 180 170 160 150 140 130 120 110 100 90 80 70 60 50 40 30 20 ppm

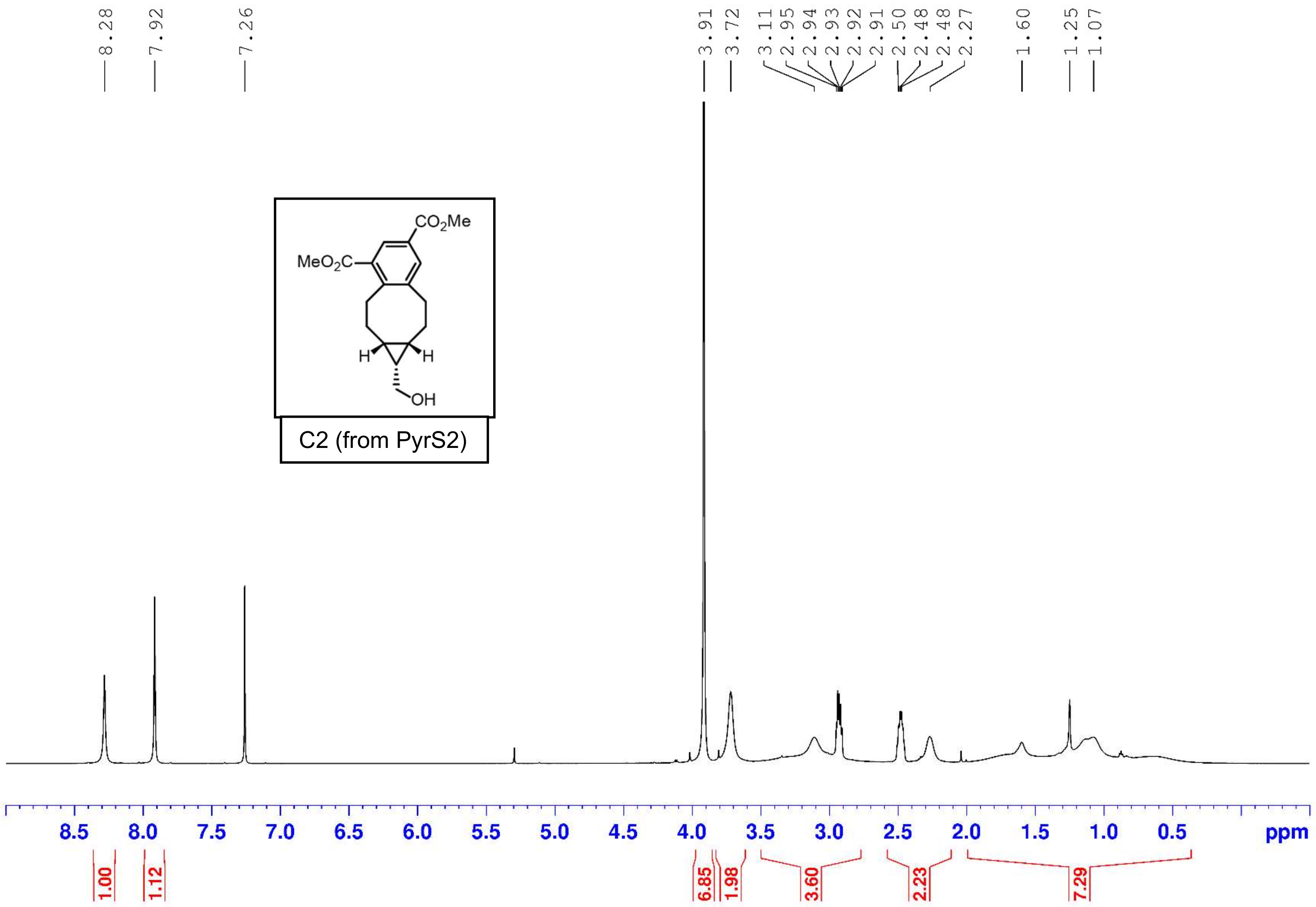

8.28
7.92
7.26
3.91
3.72
3.11
2.95
2.94
2.93
2.92
2.91
2.50
2.48
2.48
2.27
1.60
1.25
1.07
CO2Me
MeO2C
H
H
OH
C2 (from PyrS2)
8.5
8.0
7.5
7.0
6.5
6.0
5.5
5.0
4.5
4.0
3.5
3.0
2.5
2.0
1.5
1.0
0.5
ppm
1.00
1.12
6.85
1.98
3.60
2.23
7.29

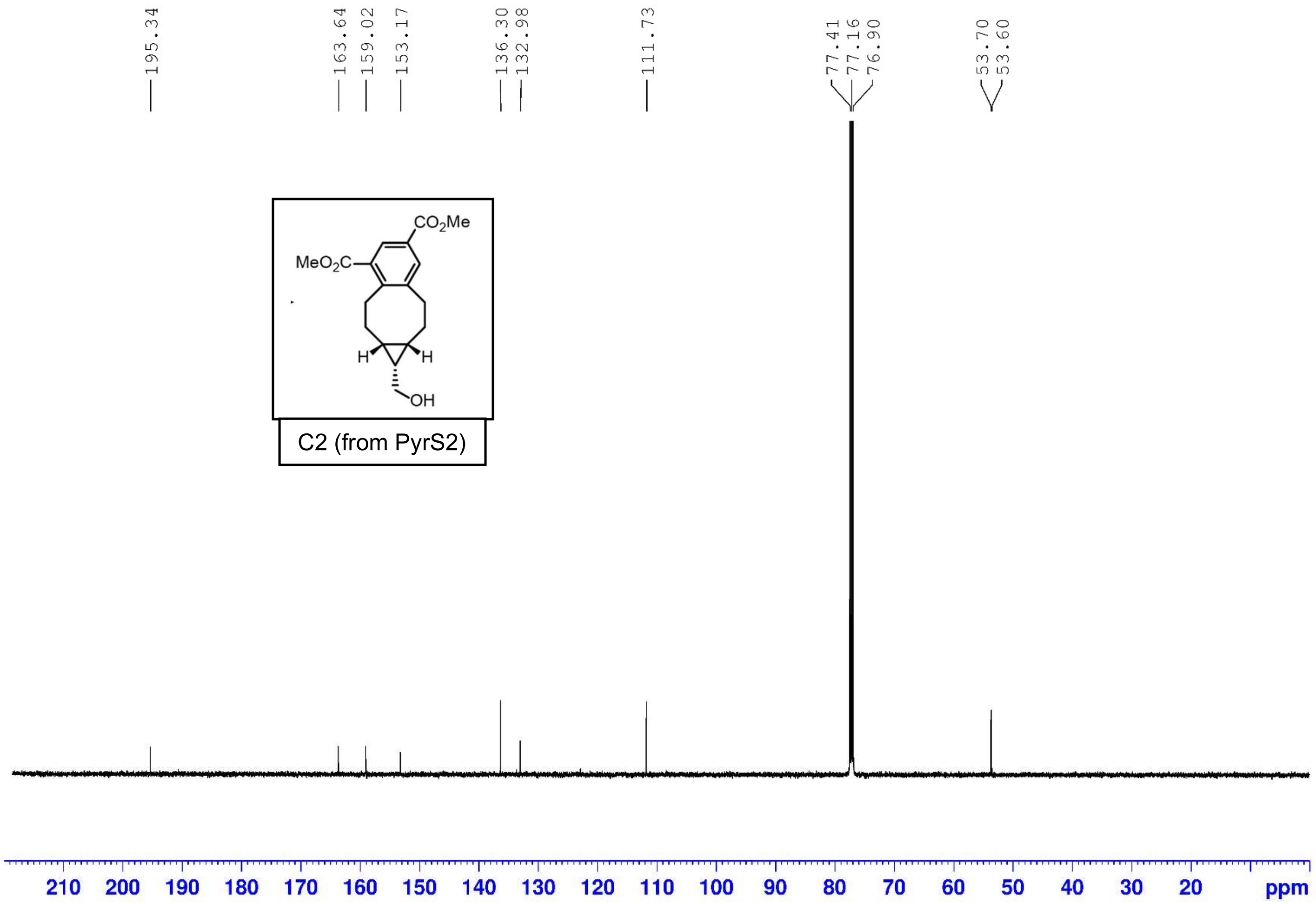
195.34
163.64
159.02
153.17
136.30
132.98
111.73
77.41
77.16
76.90
53.70
53.60
CO2Me
MeO2C
H
H
OH
C2 (from PyrS2)
210 200 190 180 170 160 150 140 130 120 110 100 90 80 70 60 50 40 30 20 ppm

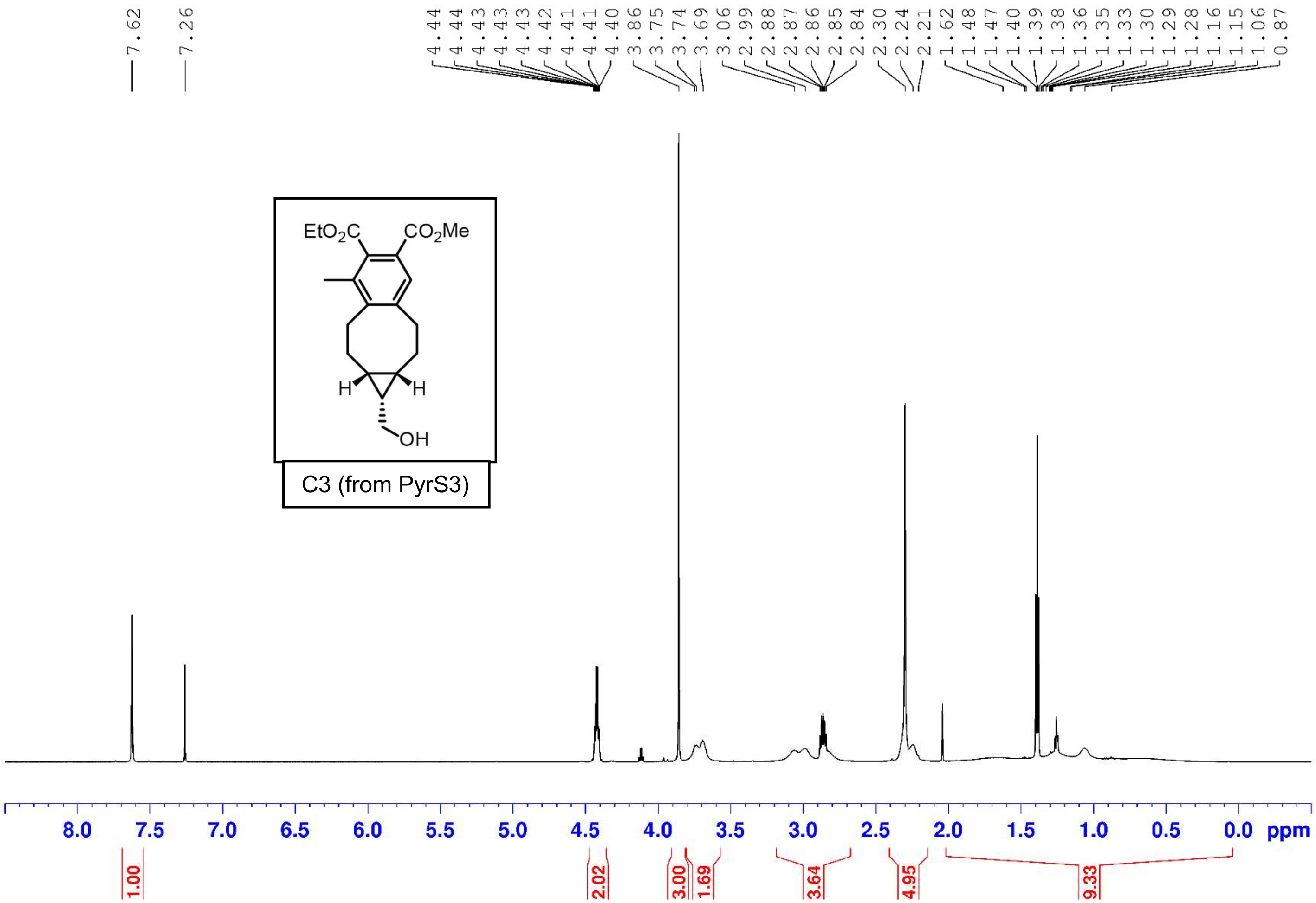
7.62
7.26
4.44
4.44
4.43
4.43
4.43
4.42
4.41
4.41
4.40
3.86
3.75
3.74
3.69
3.06
2.99
2.88
2.87
2.86
2.85
2.84
2.30
2.24
2.21
1.62
1.48
1.47
1.40
1.39
1.38
1.36
1.35
1.33
1.30
1.29
1.28
1.16
1.15
1.06
0.87
$EtO_2C$
$CO_2Me$
H
H
OH
C3 (from PyrS3)
8.0
7.5
7.0
6.5
6.0
5.5
5.0
4.5
4.0
3.5
3.0
2.5
2.0
1.5
1.0
0.5
0.0
ppm
1.00
2.02
3.00
1.69
3.64
4.95
9.33

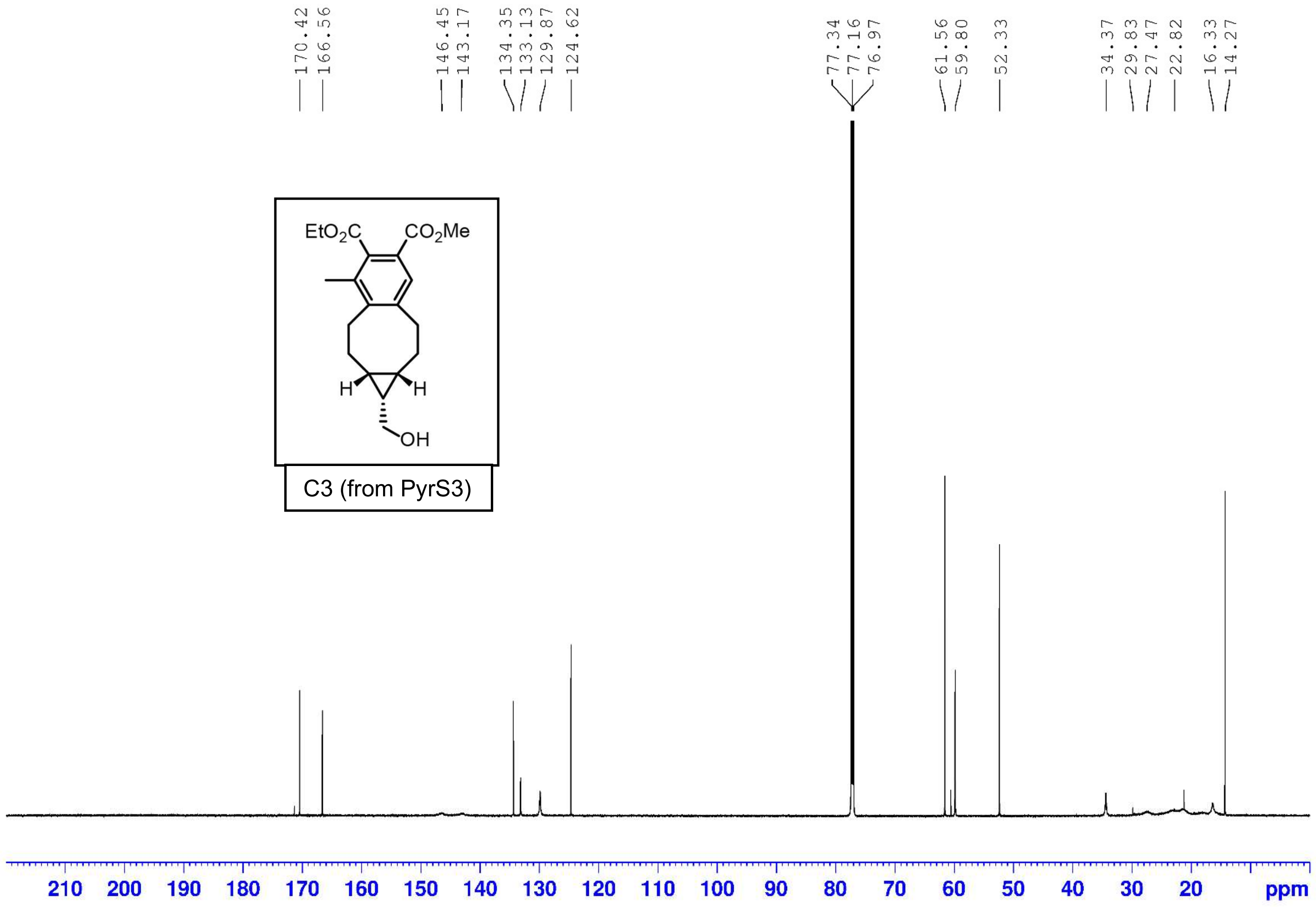
170.42
166.56
146.45
143.17
134.35
133.13
129.87
124.62
77.34
77.16
76.97
61.56
59.80
52.33
34.37
29.83
27.47
22.82
16.33
14.27
$EtO_2C$
$CO_2Me$
H
H
OH
C3 (from PyrS3)
210
200
190
180
170
160
150
140
130
120
110
100
90
80
70
60
50
40
30
20
ppm

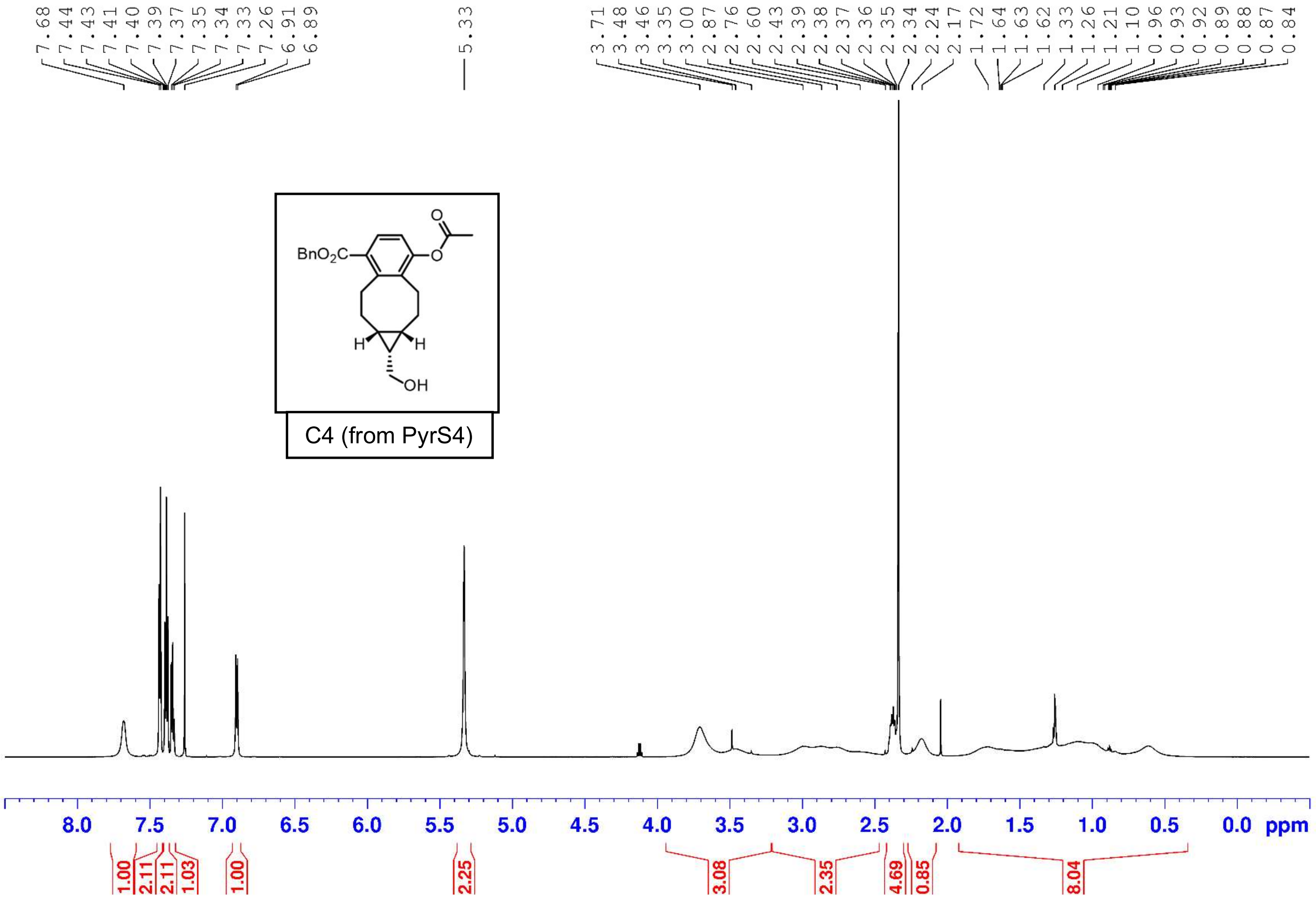
7.68
7.44
7.43
7.41
7.40
7.39
7.37
7.35
7.34
7.33
7.26
6.91
6.89
5.33
3.71
3.48
3.46
3.35
3.00
2.87
2.76
2.60
2.43
2.39
2.38
2.37
2.36
2.35
2.34
2.24
2.17
1.72
1.64
1.63
1.62
1.33
1.26
1.21
1.10
0.96
0.93
0.92
0.89
0.88
0.87
0.84
BnO2C
O
O
H
H
OH
C4 (from PyrS4)
8.0
7.5
7.0
6.5
6.0
5.5
5.0
4.5
4.0
3.5
3.0
2.5
2.0
1.5
1.0
0.5
0.0 ppm
1.00
2.11
2.11
1.03
1.00
2.25
3.08
2.35
4.69
0.85
8.04

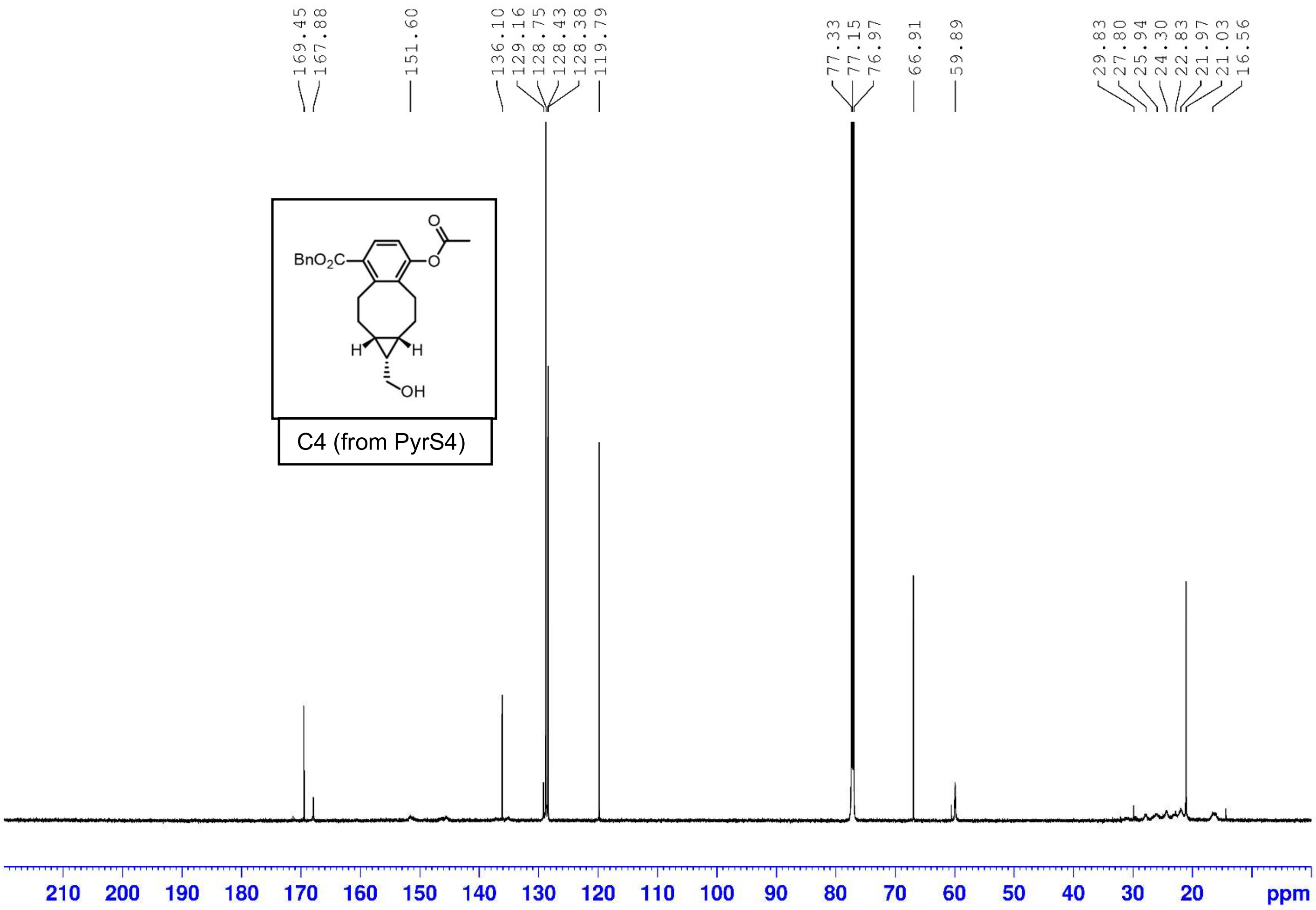
169.45
167.88
151.60
136.10
129.16
128.75
128.43
128.38
119.79
77.33
77.15
76.97
66.91
59.89
29.83
27.80
25.94
24.30
22.83
21.97
21.03
16.56
BnO2C
O
O
H
H
OH
C4 (from PyrS4)
210
200
190
180
170
160
150
140
130
120
110
100
90
80
70
60
50
40
30
20
ppm

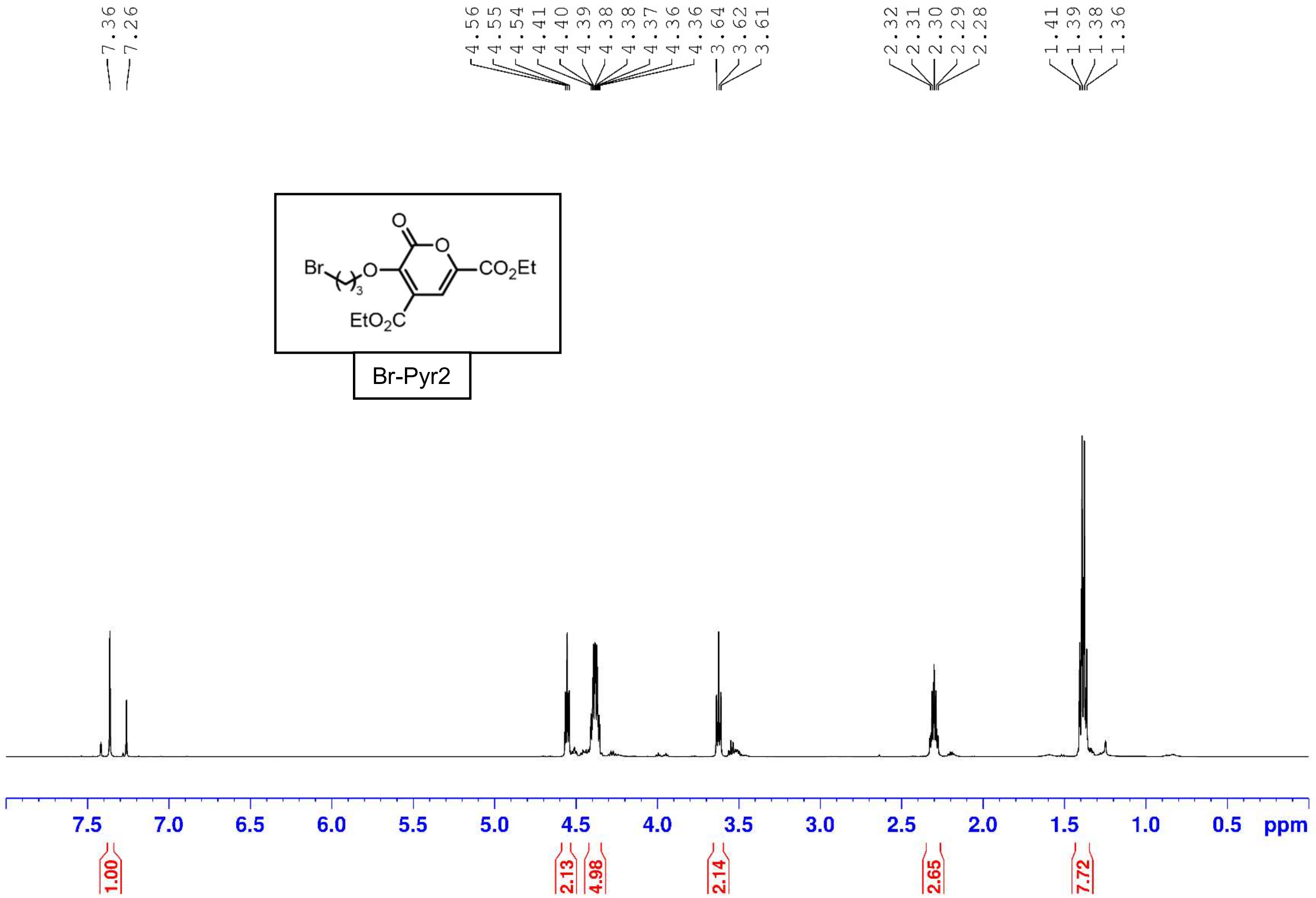
7.36
7.26
4.56
4.55
4.54
4.41
4.40
4.39
4.38
4.38
4.37
4.36
4.36
3.64
3.62
3.61
2.32
2.31
2.30
2.29
2.28
1.41
1.39
1.38
1.36
Br
O
O
O
CO2Et
EtO2C
3
Br-Pyr2
7.5
7.0
6.5
6.0
5.5
5.0
4.5
4.0
3.5
3.0
2.5
2.0
1.5
1.0
0.5
ppm
1.00
2.13
4.98
2.14
2.65
7.72

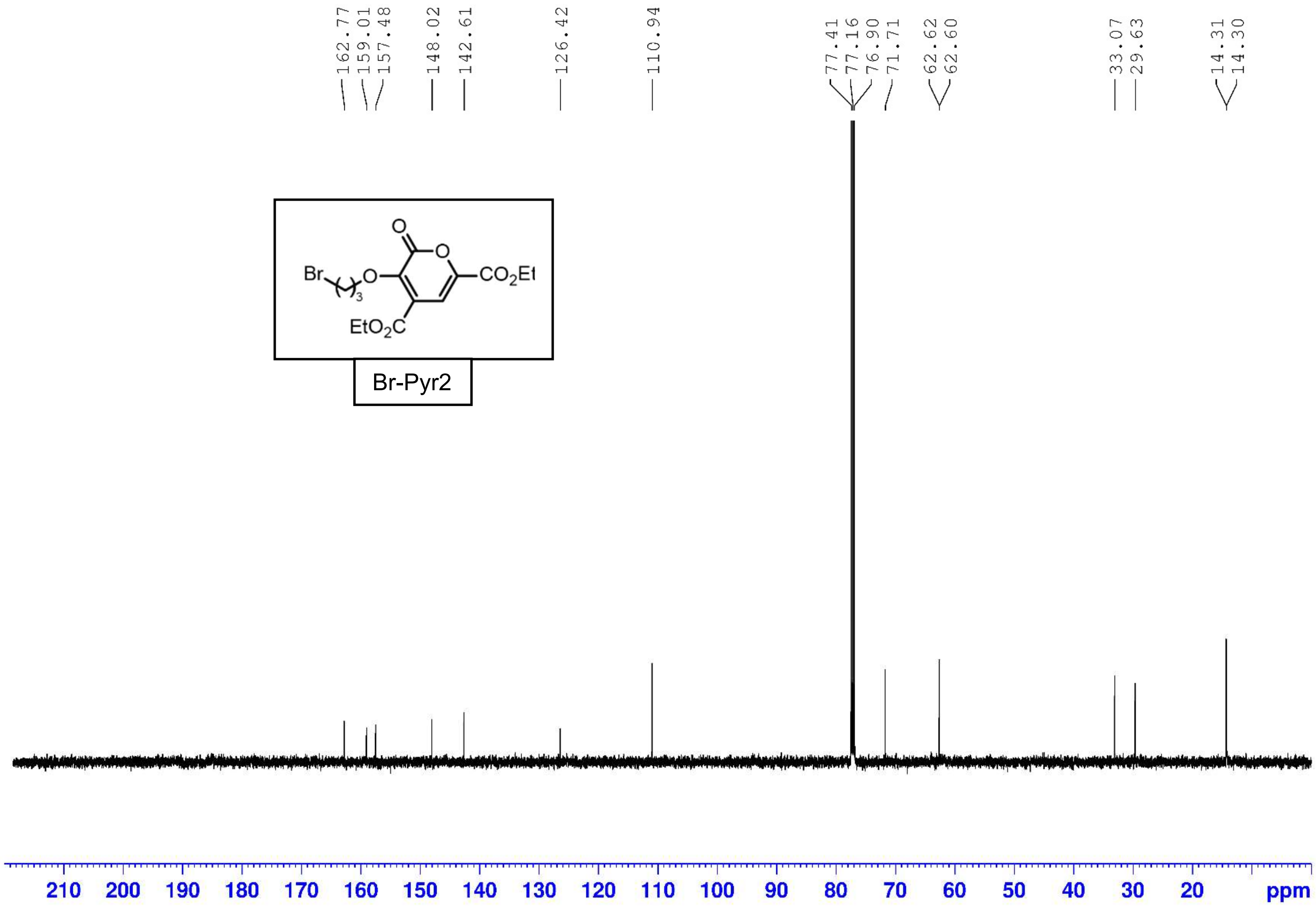
162.77
159.01
157.48
148.02
142.61
126.42
110.94
77.41
77.16
76.90
71.71
62.62
62.60
33.07
29.63
14.31
14.30
Br
O
O
O
CO2Et
EtO2C
3
Br-Pyr2
210 200 190 180 170 160 150 140 130 120 110 100 90 80 70 60 50 40 30 20 ppm

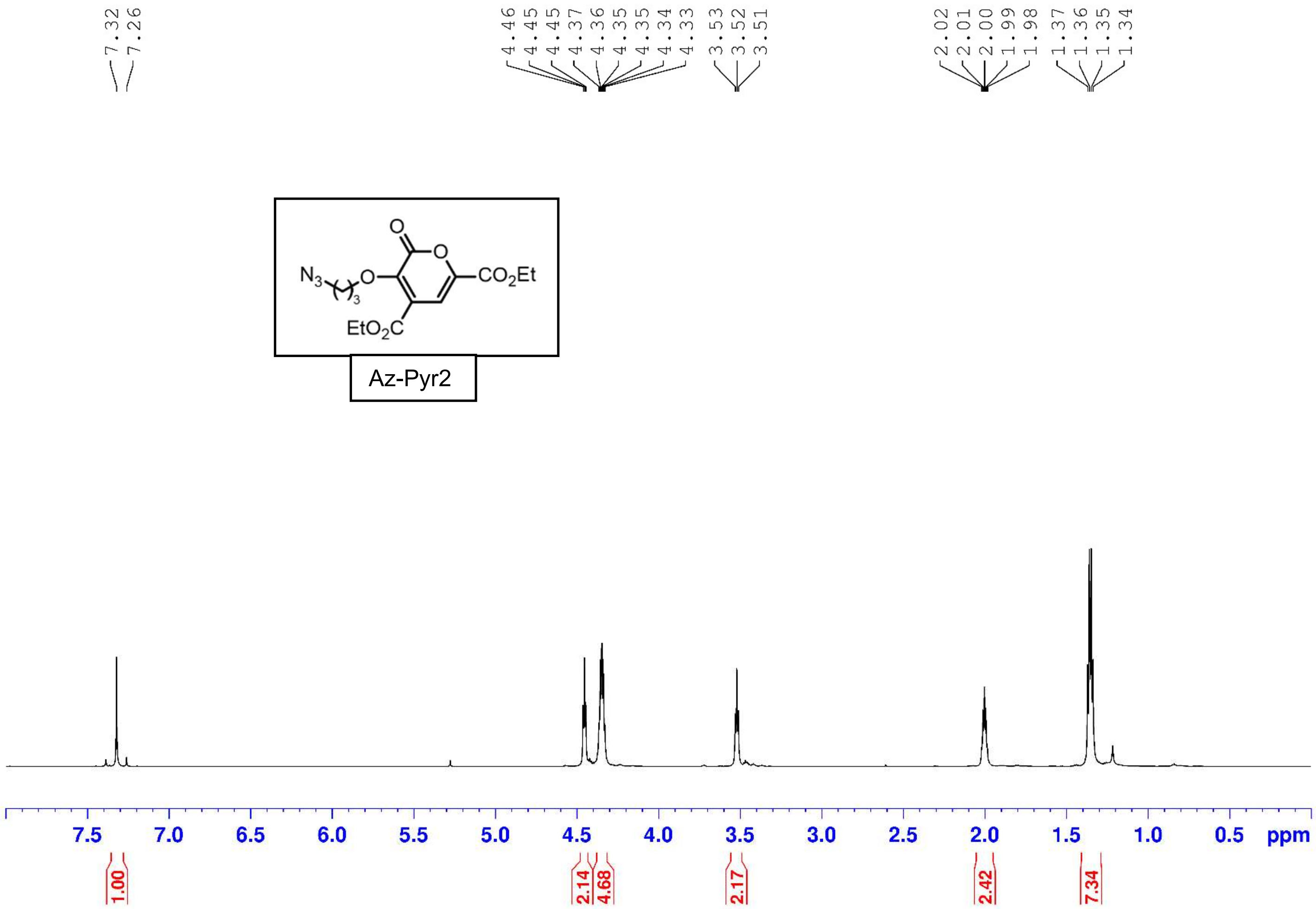
7.32
7.26
4.46
4.45
4.45
4.37
4.36
4.35
4.35
4.34
4.33
3.53
3.52
3.51
2.02
2.01
2.00
1.99
1.98
1.37
1.36
1.35
1.34
Az-Pyr2
7.5 7.0 6.5 6.0 5.5 5.0 4.5 4.0 3.5 3.0 2.5 2.0 1.5 1.0 0.5 ppm
1.00
2.14
4.68
2.17
2.42
7.34

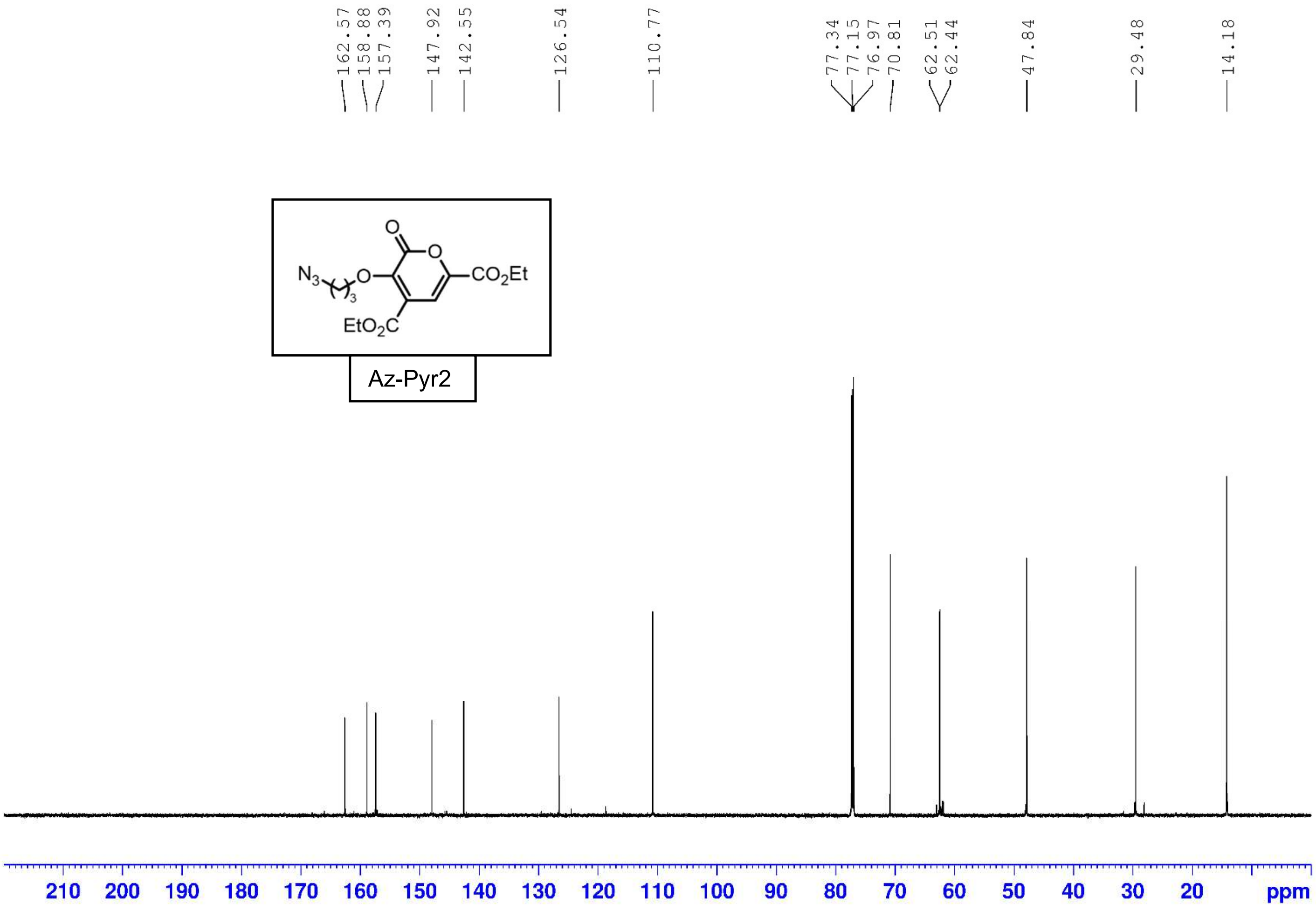
162.57
158.88
157.39
147.92
142.55
126.54
110.77
77.34
77.15
76.97
70.81
62.51
62.44
47.84
29.48
14.18
N3
O
O
O
CO2Et
EtO2C
3
Az-Pyr2
210
200
190
180
170
160
150
140
130
120
110
100
90
80
70
60
50
40
30
20
ppm

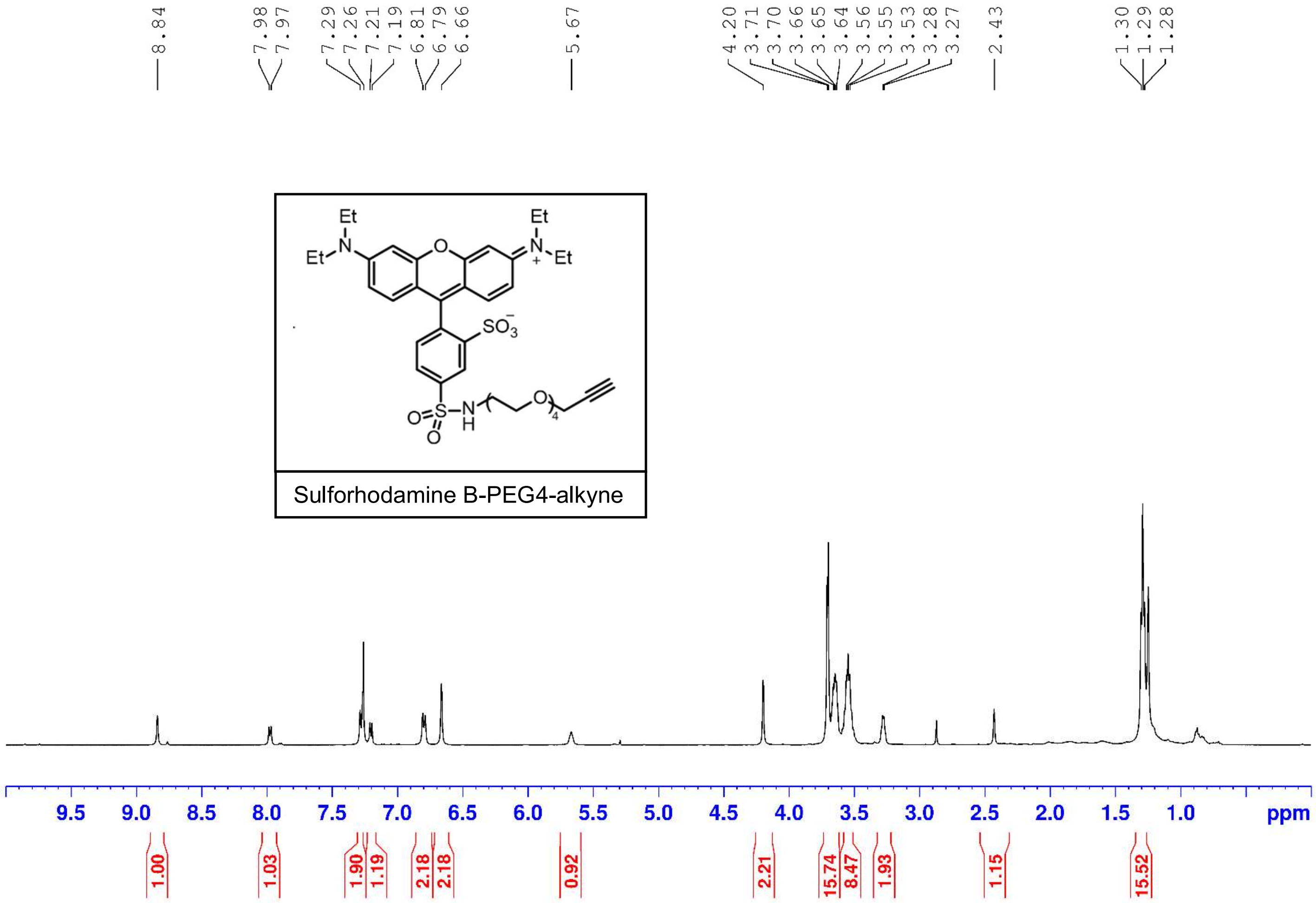
8.84
7.98
7.97
7.29
7.26
7.21
7.19
6.81
6.79
6.66
5.67
4.20
3.71
3.70
3.66
3.65
3.64
3.56
3.55
3.53
3.28
3.27
2.43
1.30
1.29
1.28
Et
N
Et
O
Et
N
+
Et
SO3
O=S
O
N
H
O
4
Sulforhodamine B-PEG4-alkyne
9.5
9.0
8.5
8.0
7.5
7.0
6.5
6.0
5.5
5.0
4.5
4.0
3.5
3.0
2.5
2.0
1.5
1.0
ppm
1.00
1.03
1.90
1.19
2.18
2.18
0.92
2.21
15.74
8.47
1.93
1.15
15.52

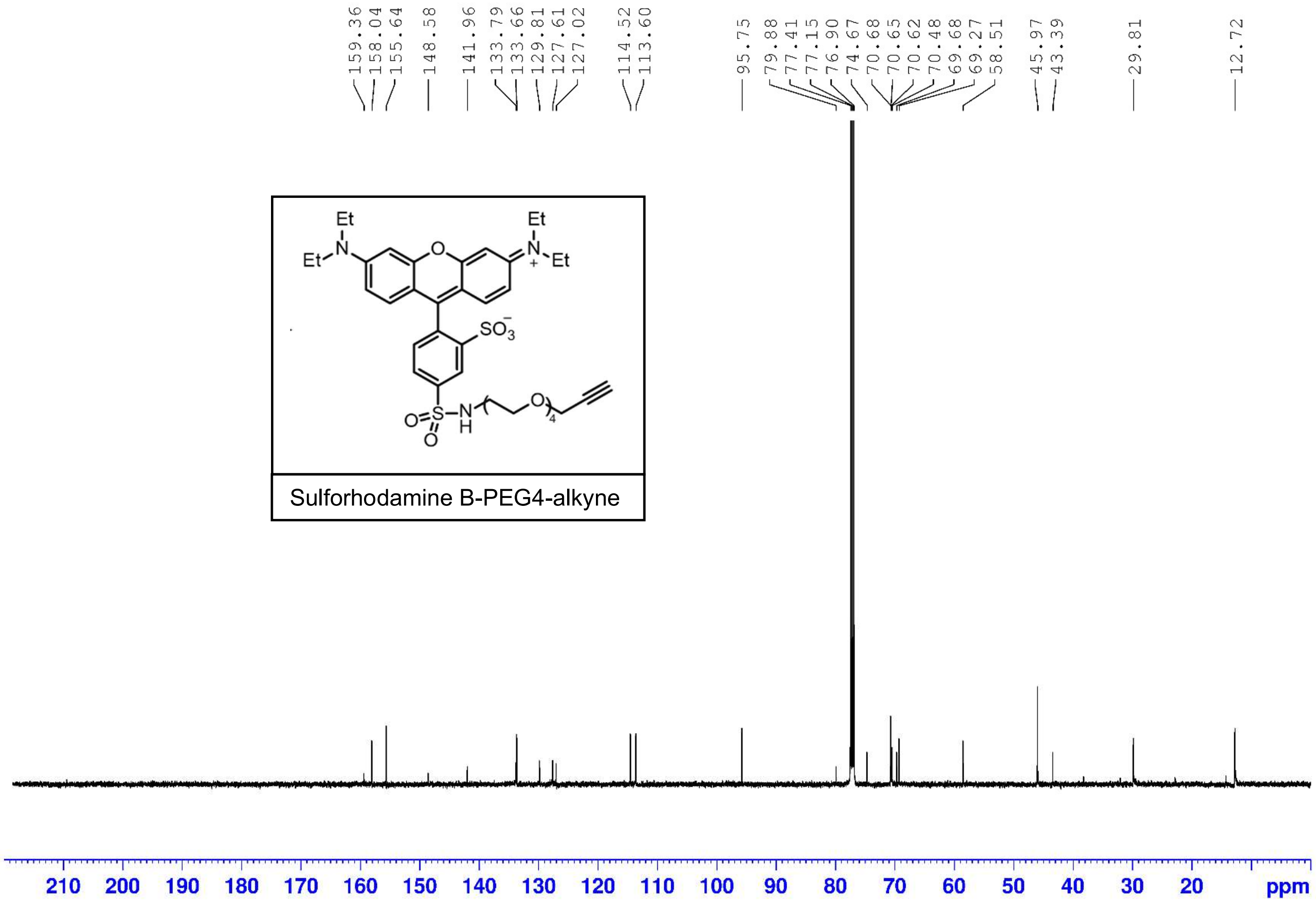
159.36
158.04
155.64
148.58
141.96
133.79
133.66
129.81
127.61
127.02
114.52
113.60
95.75
79.88
77.41
77.15
76.90
74.67
70.68
70.65
70.62
70.48
69.68
69.27
58.51
45.97
43.39
29.81
12.72
Et
N
O
SO3
S
NH
4
Sulforhodamine B-PEG4-alkyne
210 200 190 180 170 160 150 140 130 120 110 100 90 80 70 60 50 40 30 20 ppm

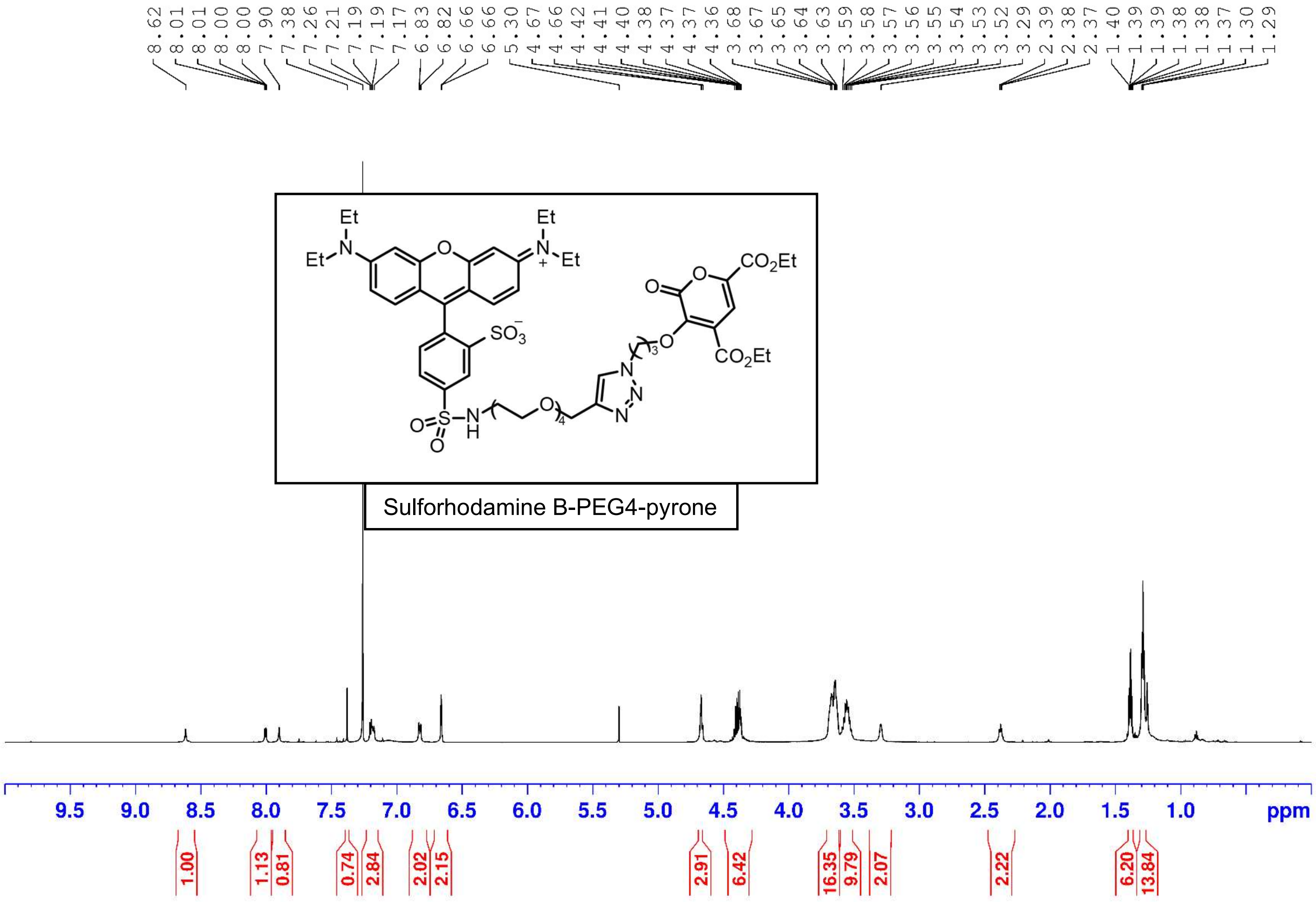
8.62 8.01 8.01 8.00 8.00 7.90 7.38 7.26 7.21 7.19 7.19 7.17 6.83 6.82 6.66 6.66 5.30 4.67 4.66 4.42 4.41 4.40 4.38 4.37 4.37 4.36 3.68 3.67 3.65 3.64 3.63 3.59 3.58 3.57 3.56 3.55 3.54 3.53 3.52 3.29 2.39 2.38 2.37 1.40 1.39 1.39 1.38 1.37 1.30 1.29
Sulforhodamine B-PEG4-pyrone
9.5 9.0 8.5 8.0 7.5 7.0 6.5 6.0 5.5 5.0 4.5 4.0 3.5 3.0 2.5 2.0 1.5 1.0 ppm
1.00 1.13 0.81 0.74 2.84 2.02 2.15 2.91 6.42 16.35 9.79 2.07 2.22 6.20 13.84

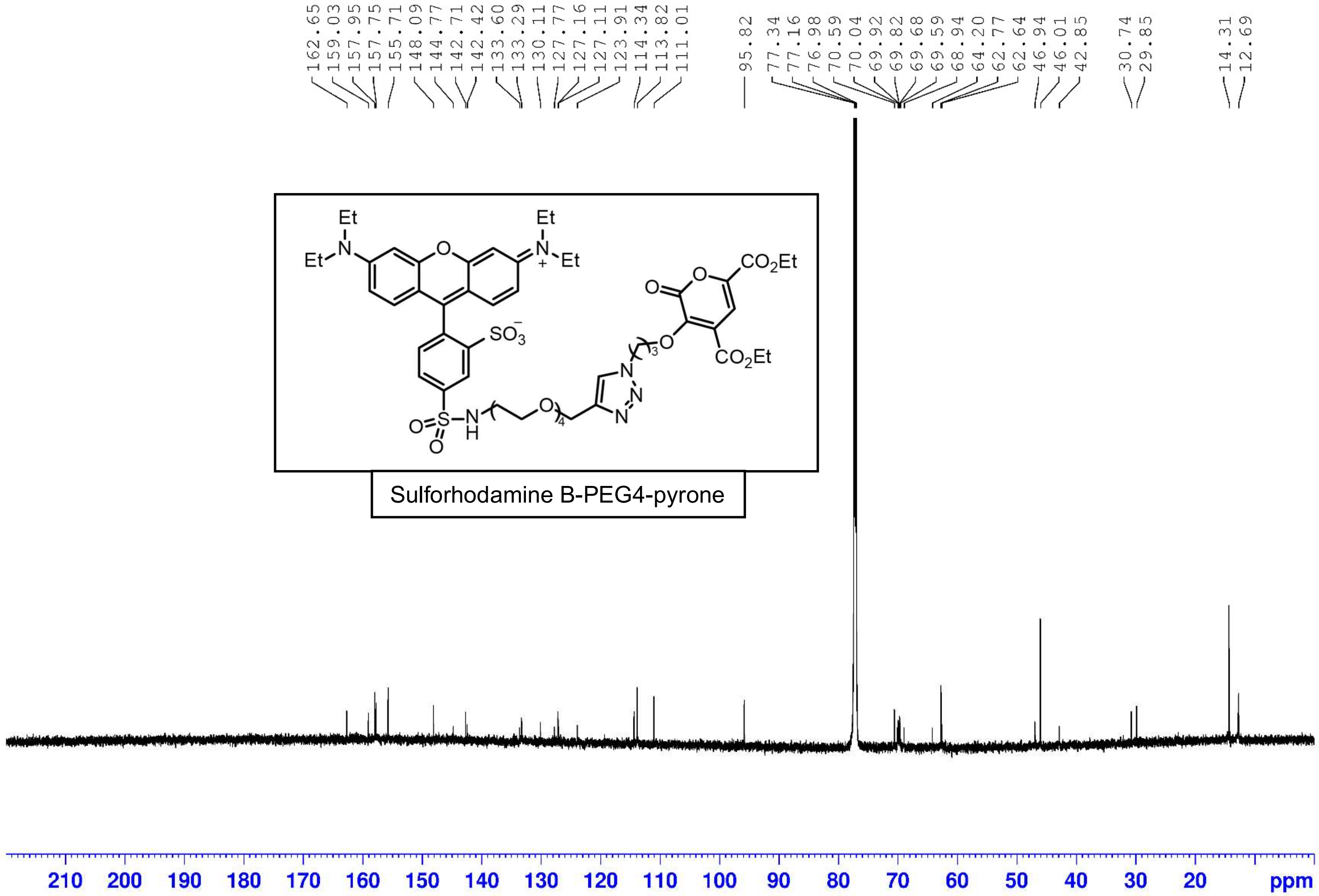
162.65
159.03
157.95
157.75
155.71
148.09
144.77
142.71
142.42
133.60
133.29
130.11
127.77
127.16
127.11
123.91
114.34
113.82
111.01
95.82
77.34
77.16
76.98
70.59
70.04
69.92
69.82
69.68
69.59
68.94
64.20
62.77
62.64
46.94
46.01
42.85
30.74
29.85
14.31
12.69
Sulforhodamine B-PEG4-pyrone
210 200 190 180 170 160 150 140 130 120 110 100 90 80 70 60 50 40 30 20 ppm

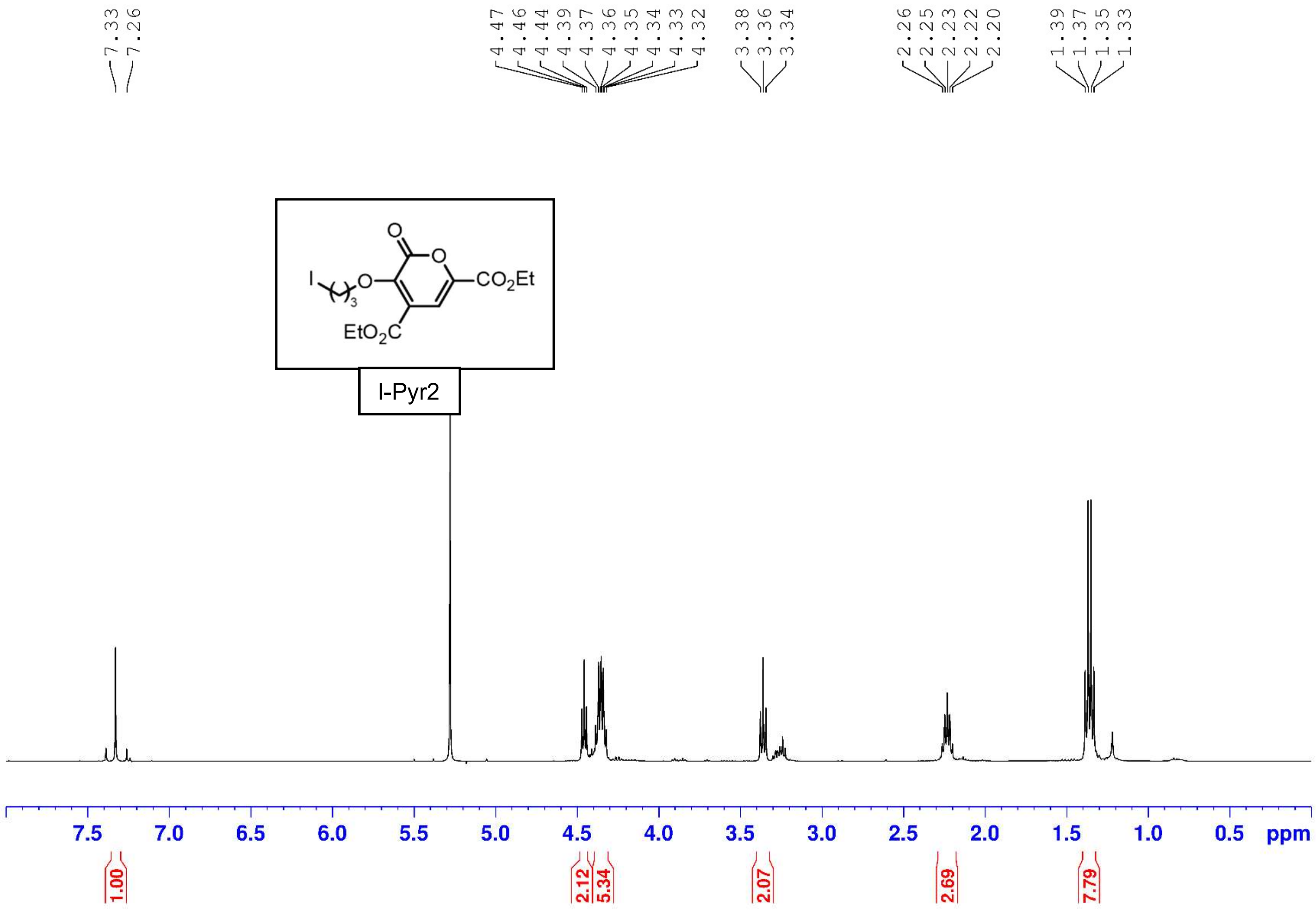
7.33
7.26
4.47
4.46
4.44
4.39
4.37
4.36
4.35
4.34
4.33
4.32
3.38
3.36
3.34
2.26
2.25
2.23
2.22
2.20
1.39
1.37
1.35
1.33
I-Pyr2
7.5 7.0 6.5 6.0 5.5 5.0 4.5 4.0 3.5 3.0 2.5 2.0 1.5 1.0 0.5 ppm
1.00
2.12
5.34
2.07
2.69
7.79

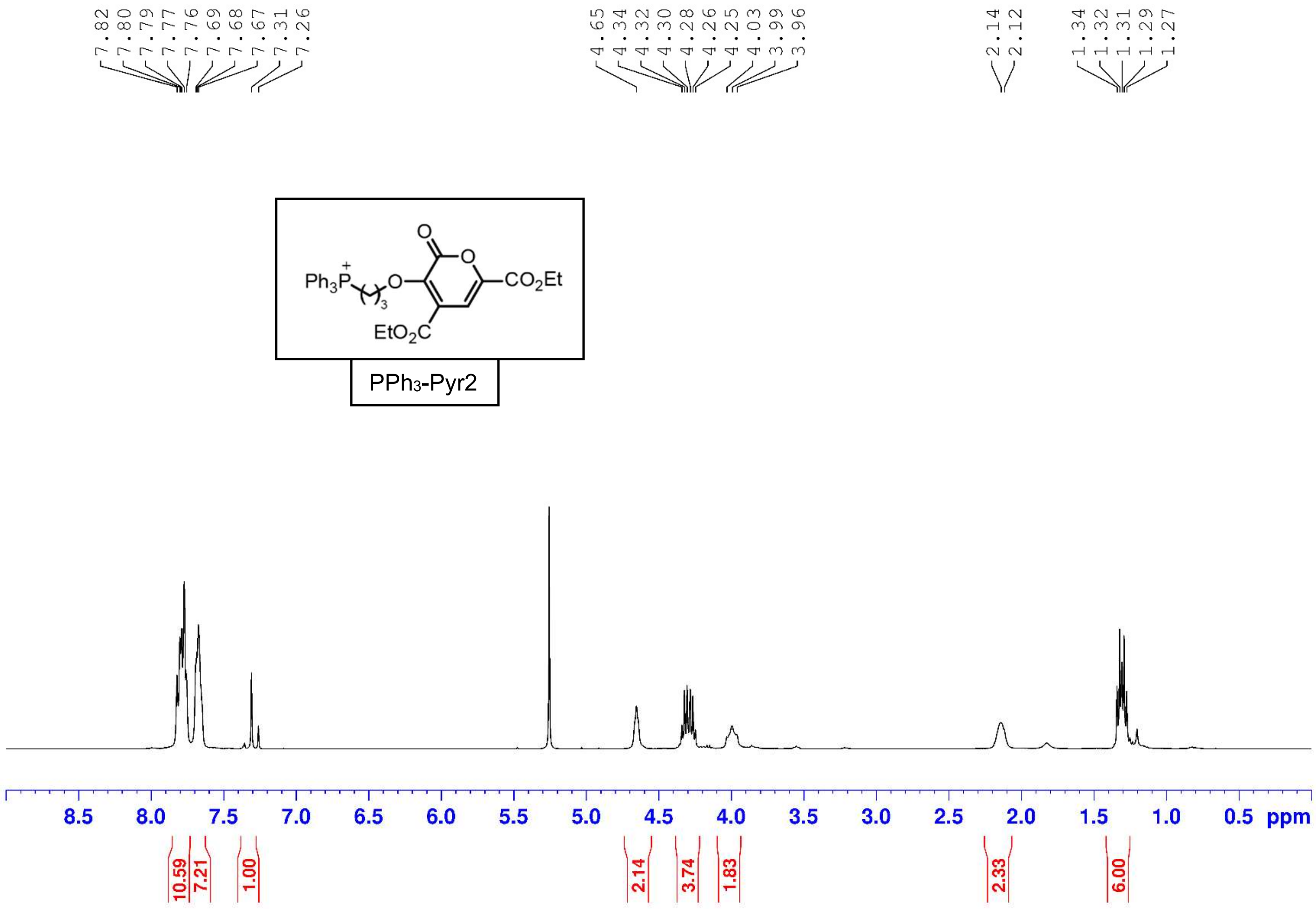
7.82
7.80
7.79
7.77
7.76
7.69
7.68
7.67
7.31
7.26
4.65
4.34
4.32
4.30
4.28
4.26
4.25
4.03
3.99
3.96
2.14
2.12
1.34
1.32
1.31
1.29
1.27
PPh3-Pyr2
8.5
8.0
7.5
7.0
6.5
6.0
5.5
5.0
4.5
4.0
3.5
3.0
2.5
2.0
1.5
1.0
0.5
ppm
10.59
7.21
1.00
2.14
3.74
1.83
2.33
6.00

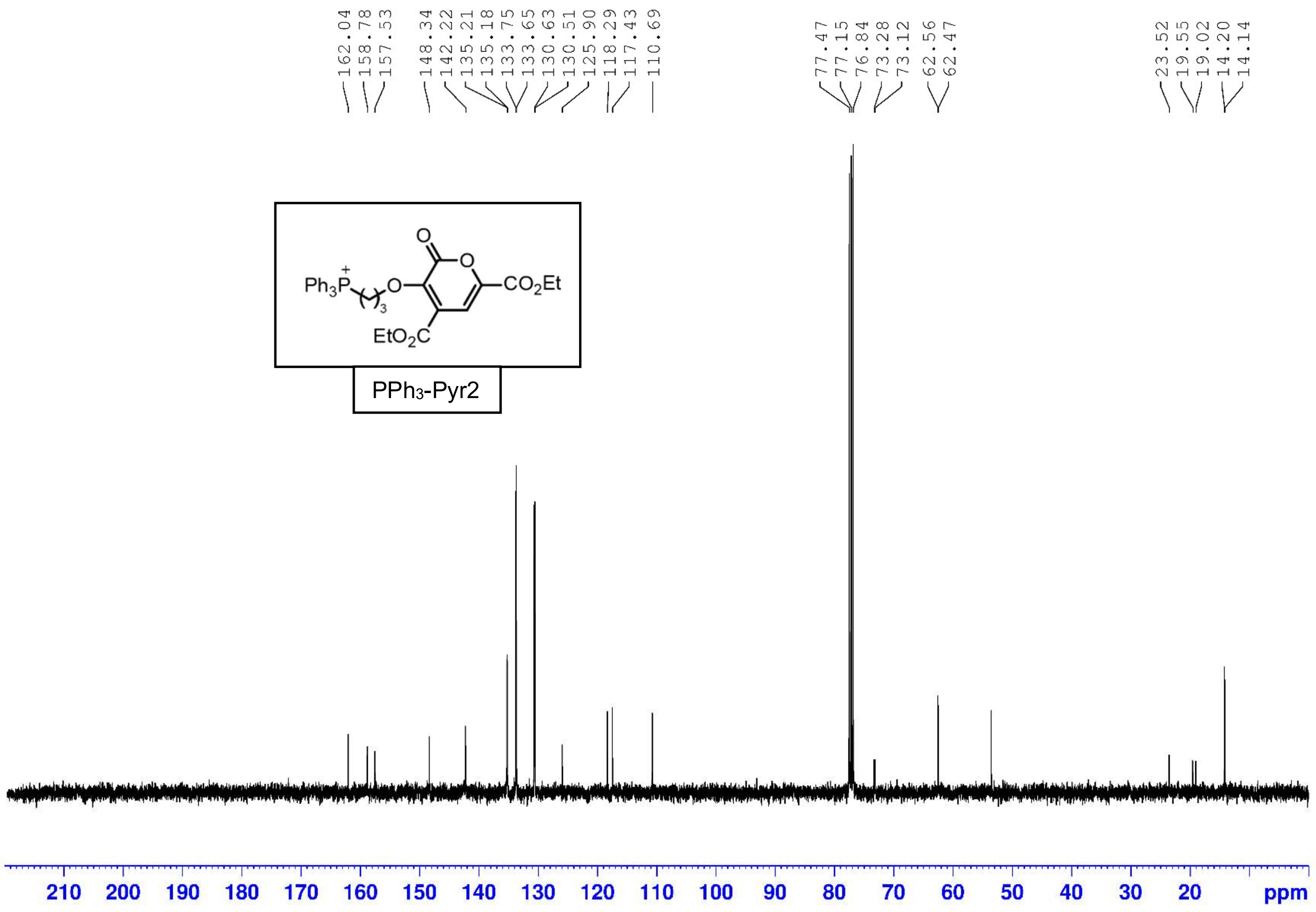
162.04
158.78
157.53
148.34
142.22
135.21
135.18
133.75
133.65
130.63
130.51
125.90
118.29
117.43
110.69
77.47
77.15
76.84
73.28
73.12
62.56
62.47
23.52
19.55
19.02
14.20
14.14
Ph3P
O
CO2Et
EtO2C
PPh3-Pyr2
210 200 190 180 170 160 150 140 130 120 110 100 90 80 70 60 50 40 30 20 ppm

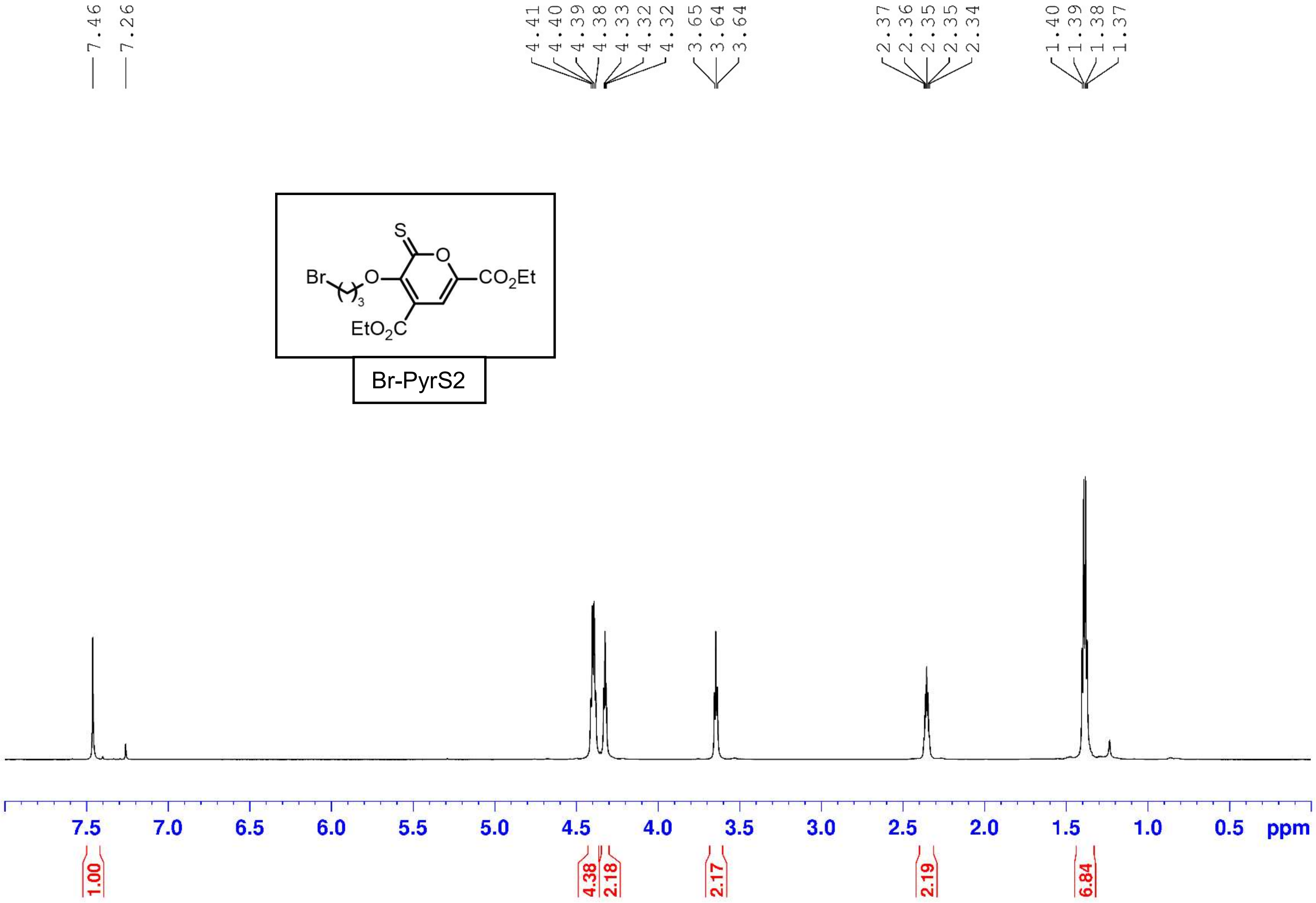
7.46
7.26
4.41
4.40
4.39
4.38
4.33
4.32
4.32
3.65
3.64
3.64
2.37
2.36
2.35
2.35
2.34
1.40
1.39
1.38
1.37
S
O
Br
O
3
$CO_2Et$
$EtO_2C$
Br-PyrS2
7.5
7.0
6.5
6.0
5.5
5.0
4.5
4.0
3.5
3.0
2.5
2.0
1.5
1.0
0.5
ppm
1.00
4.38
2.18
2.17
2.19
6.84

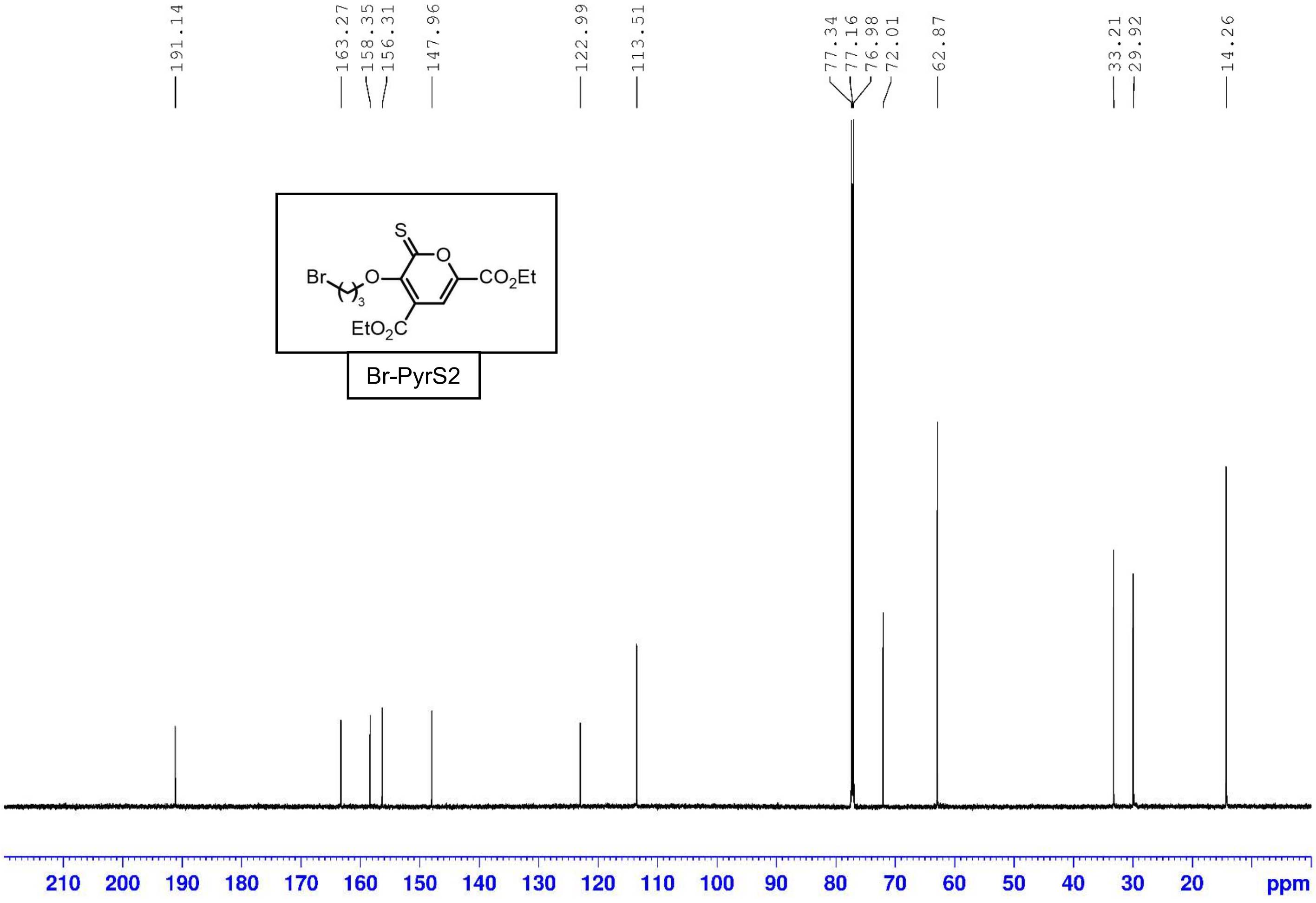
191.14
163.27
158.35
156.31
147.96
122.99
113.51
77.34
77.16
76.98
72.01
62.87
33.21
29.92
14.26
Br
O
S
O
CO2Et
3
EtO2C
Br-PyrS2
210 200 190 180 170 160 150 140 130 120 110 100 90 80 70 60 50 40 30 20 ppm

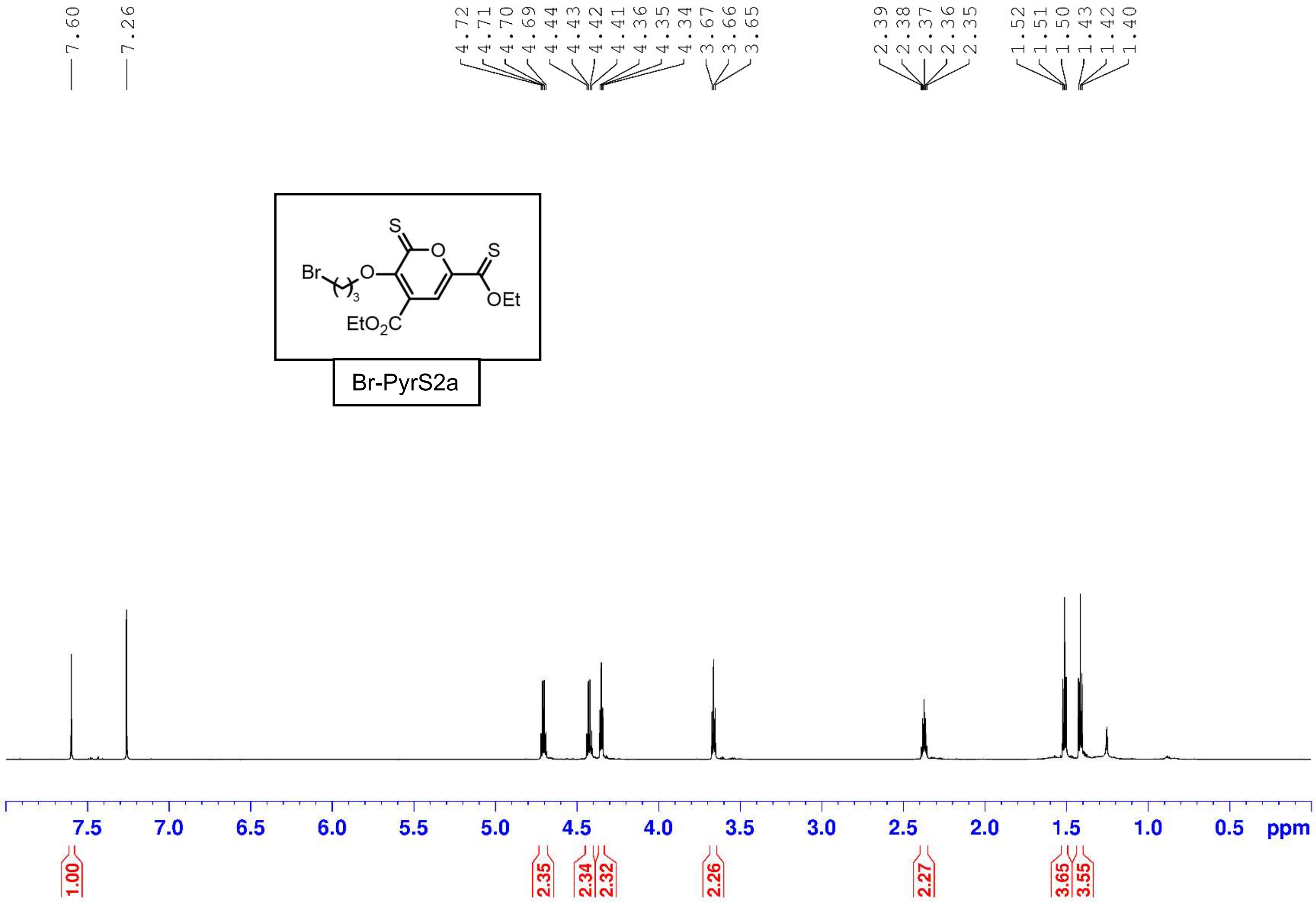

7.60
7.26
4.72
4.71
4.70
4.69
4.44
4.43
4.42
4.41
4.36
4.35
4.34
3.67
3.66
3.65
2.39
2.38
2.37
2.36
2.35
1.52
1.51
1.50
1.43
1.42
1.40
Br
O
S
O
S
OEt
EtO2C
3
Br-PyrS2a
7.5 7.0 6.5 6.0 5.5 5.0 4.5 4.0 3.5 3.0 2.5 2.0 1.5 1.0 0.5 ppm
1.00
2.35
2.34
2.32
2.26
2.27
3.65
3.55

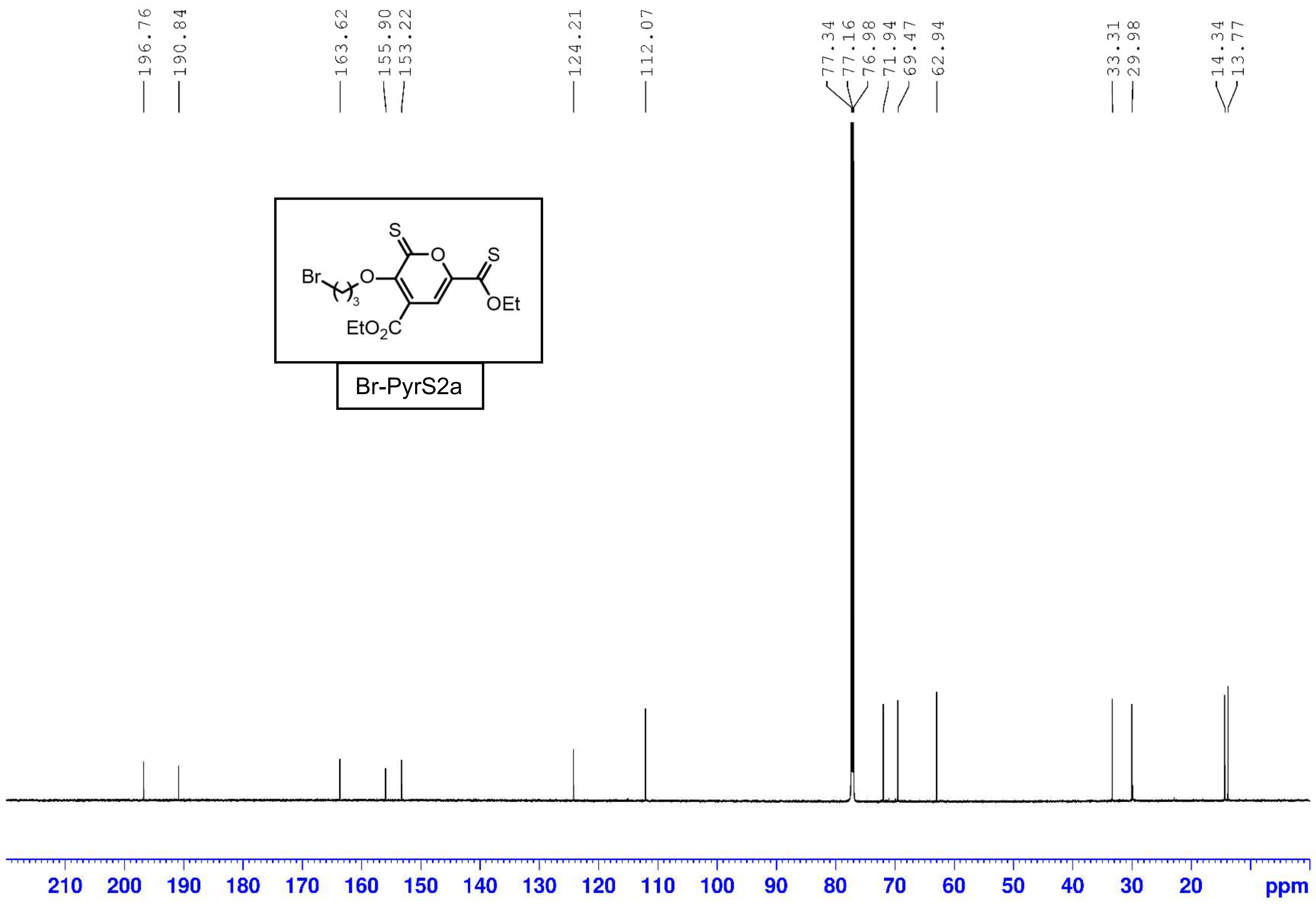
196.76
190.84
163.62
155.90
153.22
124.21
112.07
77.34
77.16
76.98
71.94
69.47
62.94
33.31
29.98
14.34
13.77
Br
O
S
O
S
OEt
3
EtO2C
Br-PyrS2a
210
200
190
180
170
160
150
140
130
120
110
100
90
80
70
60
50
40
30
20
ppm

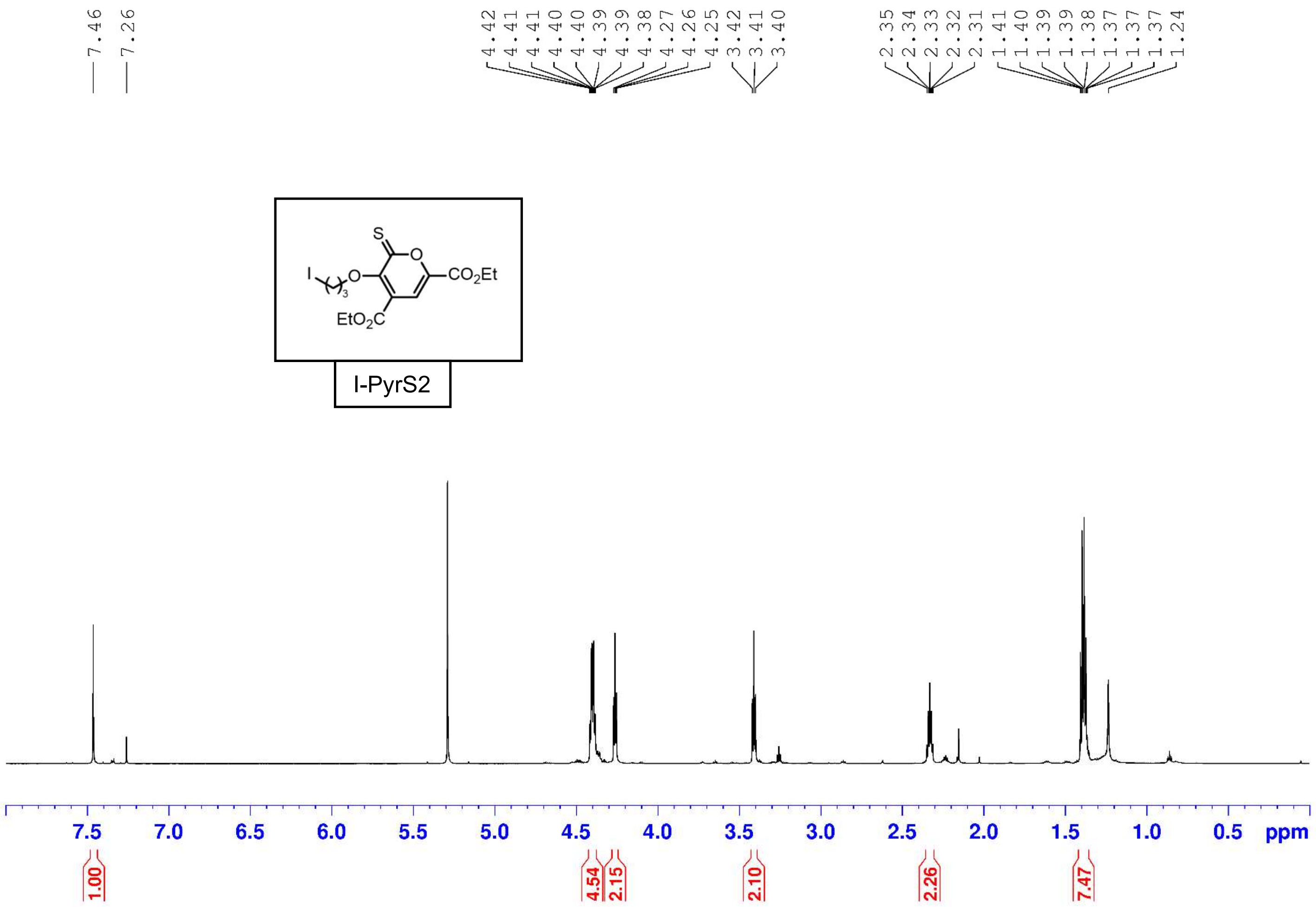
S
O
I
O
3
CO2Et
EtO2C
I-PyrS2
7.46
7.26
4.42
4.41
4.41
4.40
4.40
4.39
4.39
4.38
4.27
4.26
4.25
3.42
3.41
3.40
2.35
2.34
2.33
2.32
2.31
1.41
1.40
1.39
1.39
1.38
1.37
1.37
1.37
1.24
7.5
7.0
6.5
6.0
5.5
5.0
4.5
4.0
3.5
3.0
2.5
2.0
1.5
1.0
0.5
ppm
1.00
4.54
2.15
2.10
2.26
7.47

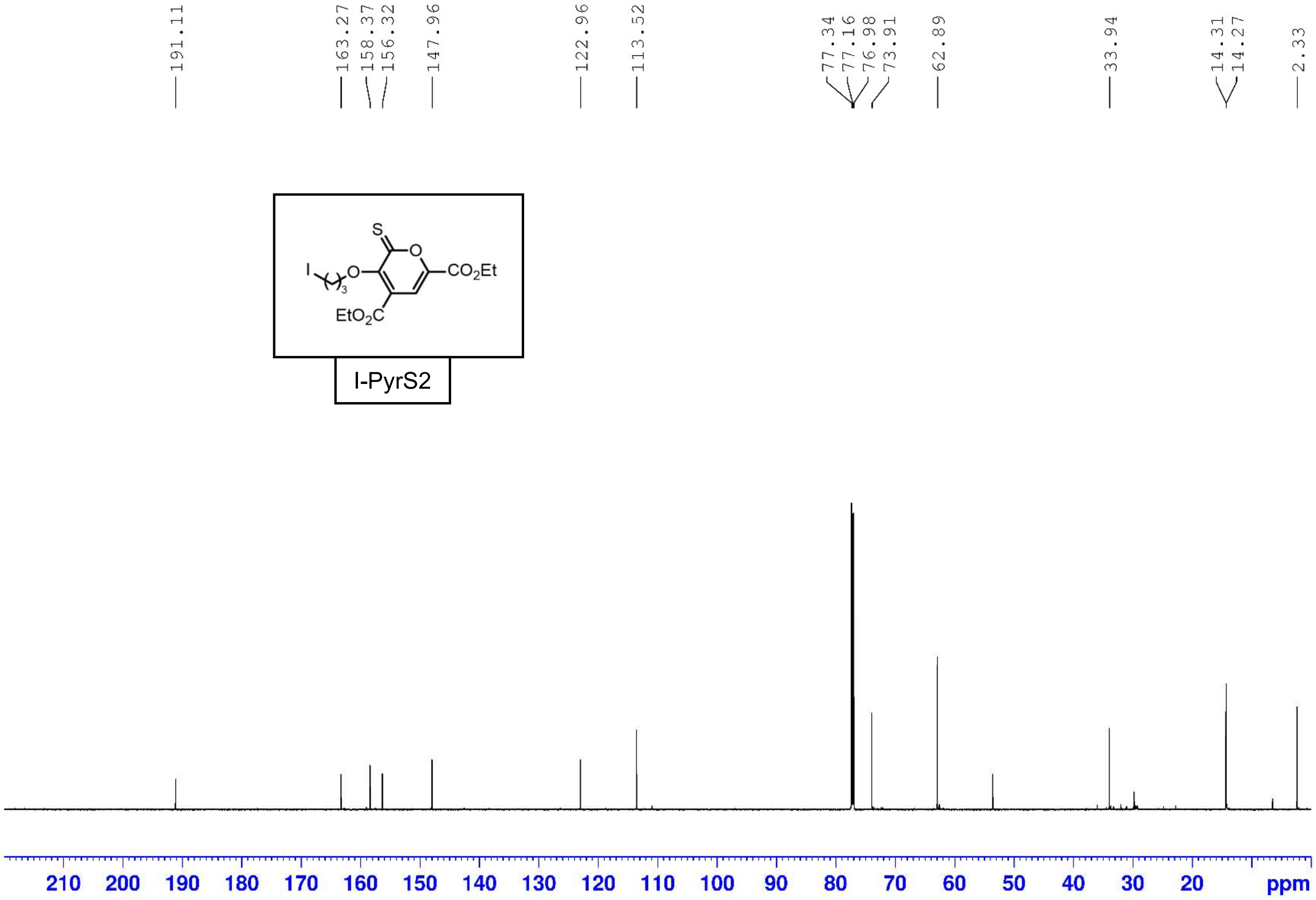
191.11
163.27
158.37
156.32
147.96
122.96
113.52
77.34
77.16
76.98
73.91
62.89
33.94
14.31
14.27
2.33
S
O
I
O
3
CO2Et
EtO2C
210 200 190 180 170 160 150 140 130 120 110 100 90 80 70 60 50 40 30 20 ppm

I-PyrS2

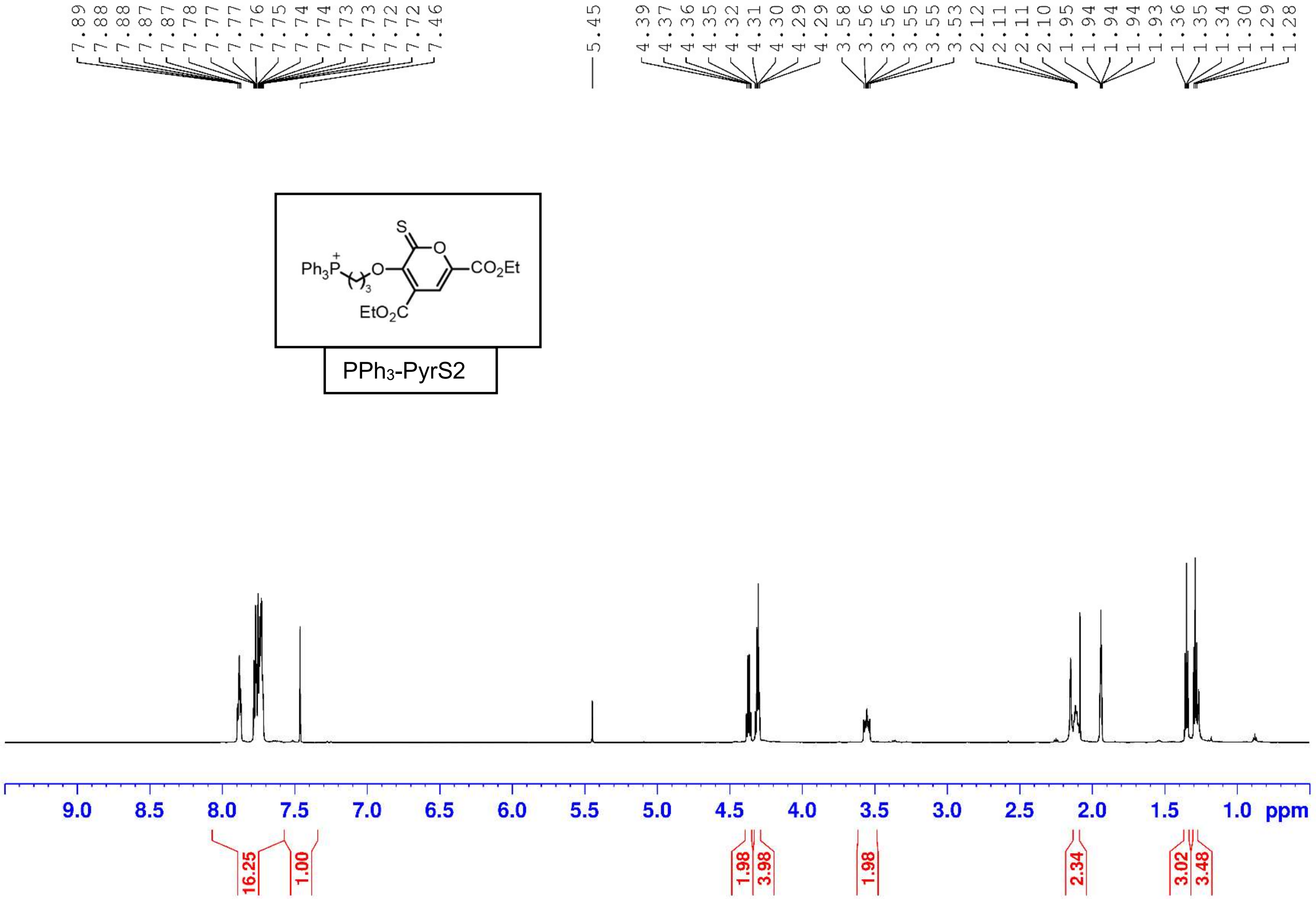
PPh3-PyrS2
ppm

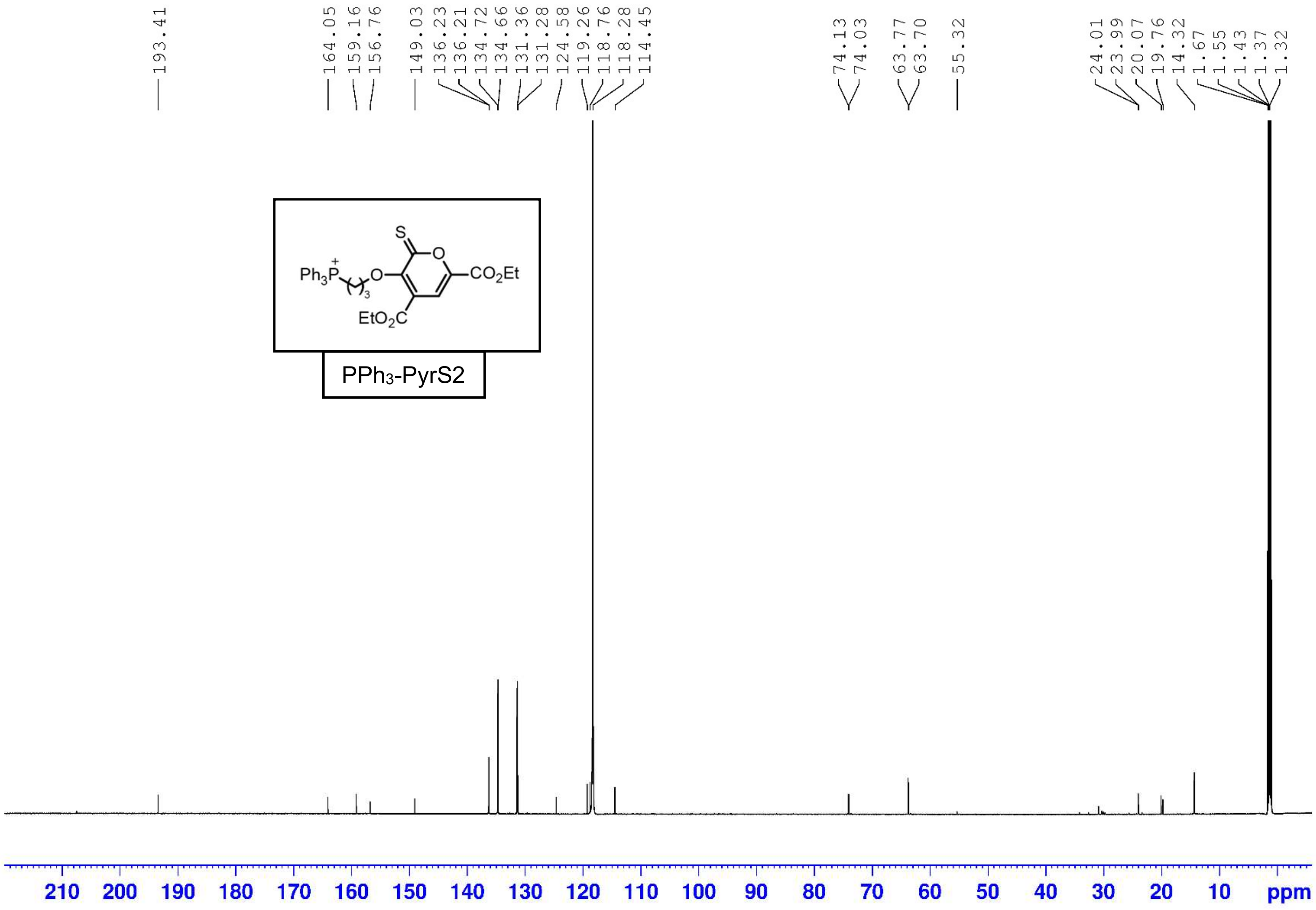
193.41
164.05
159.16
156.76
149.03
136.23
136.21
134.72
134.66
131.36
131.28
124.58
119.26
118.76
118.28
114.45
74.13
74.03
63.77
63.70
55.32
24.01
23.99
20.07
19.76
14.32
1.67
1.55
1.43
1.37
1.32
S
O
Ph3P
O
CO2Et
EtO2C
210 200 190 180 170 160 150 140 130 120 110 100 90 80 70 60 50 40 30 20 10 ppm

PPh3-PyrS2

(6) Gaussian 16, Revision C.01, Frisch, M. J., Trucks, G. W., Schlegel, H. B., Scuseria, G. E., Robb, M. A., Cheeseman, J. R., Scalmani, G., Barone, V., Petersson, G. A., Nakatsuji, H., Li, X., Caricato, M., Marenich, A. V., Bloino, J., Janesko, B. G., Gomperts, R., Mennucci, B., Hratchian, H. P., Ortiz, J. V., Izmaylov, A. F., Sonnenberg, J. L., Williams-Young, D., Ding, F., Lipparini, F., Egidi, F., Goings, J., Peng, B., Petrone, A., Henderson, T., Ranasinghe, D., Zakrzewski, V. G., Gao, J., Rega, N., Zheng, G., Liang, W., Hada, M., Ehara, M., Toyota, K., Fukuda, R., Hasegawa, J., Ishida, M., Nakajima, T., Honda, Y., Kitao, O., Nakai, H., Vreven, T., Throssell, K., Montgomery, J. A., Jr., Peralta, J. E., Ogliaro, F., Bearpark, M. J., Heyd, J. J., Brothers, E. N., Kudin, K. N., Staroverov, V. N., Keith, T. A., Kobayashi, R., Normand, J., Raghavachari, K., Rendell, A. P., Burant, J. C., Iyengar, S. S., Tomasi, J., Cossi, M., Millam, J. M., Klene, M., Adamo, C., Cammi, R., Ochterski, J. W., Martin, R. L., Morokuma, K., Farkas, O., Foresman, J. B., Fox, D. J. Gaussian, Inc., Wallingford CT, 2016.

(13) Huang, W., Wen, K. Q., Laughlin, S. T. Manuscript in preparation.

(14) Huang, W., Escoihula, J., Laughlin, S. T. Manuscript in preparation.
